# Transcriptome analysis reveals the differences between cellular response to ribosomal stress and translational stress

**DOI:** 10.1101/203356

**Authors:** Md Shamsuzzaman, Brian Gregory, Vincent Bruno, Lasse Lindahl

**Affiliations:** Department of Biological Sciences, University of Maryland Baltimore County (UMBC), 1000 Hilltop Circle, Baltimore, Maryland 21250, USA; Institute for Genome Sciences, University of Maryland School of Medicine, Baltimore, Maryland 21201, USA; Department of Microbiology and Immunology, University of Maryland School of Medicine, Baltimore, Maryland 21201, USA

## Abstract

Ribosome biogenesis is an essential metabolic process of a growing cell. Cells need to continuously synthesize new ribosomes in order to make new proteins than can support building biomass and cell division. It is obvious that in the absence of ribosome biogenesis, cell growth will stop and cell division will stall. However, it is not clear whether cell growth stops due to reduced protein synthesis capacity (translational stress) or due to activation of signaling specific to ribosome biogenesis abnormalities (ribosomal stress). To understand the signaling pathways leading to cell cycle arrest under ribosomal and translational stress conditions, we performed time series RNA-seq experiments of cells at different time of ribosomal and translational stress. We found that expression of ribosomal protein genes follow different course over the time of these two stress types. In addition, ribosomal stress is sensed early in the cell, as early as 2hr. Up-regulation of genes responsive to oxidative stress and over representation of mRNAs for transcription factors responsive to stress was detected in cell at 2hr of ribosomal protein depletion. Even though, we detected phenotypic similarities in terms of cell separation and accumulation in G1 phase cells during inhibition of ribosome formation and ribosome function, different gene expression patterns underlie these phenotypes, indicating a difference in causalities of these phenotypes. Both ribosomal and translational stress show common increased expression of stress responsive gene expression, like Crz1 target gene expression, signature of oxidative stress response and finally membrane or cell wall instability. We speculate that cell membrane and cell wall acts as major stress sensor in the cell and adjust cellular metabolism accordingly. Any change in membrane lipid composition, or membrane protein oxidation, or decrease or increase in intracellular turgor pressure causes stress in cell membrane. Cell membrane or cell wall stress activates and/or inactivates specific signaling pathway which triggers stress responsive gene expression and adaptation of cellular behavior accordingly.

## Introduction

Ribosome biogenesis is a complex but highly coordinated metabolic process involving assembly of 79 ribosomal proteins (RPs) and 4 rRNAs with the assistance of more than 200 assembly factor proteins (ref). In exponentially growing cells, the biogenesis process consumes significant amount of cellular resources with more than 50% of transcriptional capacity engaged in transcription of RP, ribosome biogenesis (Ribi) and rRNA genes, and 20% or more of translational capacity engaged in translation of RPs and Ribi proteins (Warner 1999). Imbalanced production of any essential component in ribosome biogenesis process stops the biogenesis process, decreases number of mature ribosomes per cell and increases accumulation and turnover of ribosomal precursor particles. Interrupted ribosome biogenesis is defined as ribosomal stress or nucleolar stress and this stress causes cell cycle arrest in eukaryotic cells (Pestov, Strezoska et al. 2001, Fumagalli and Thomas 2011, James, Wang et al. 2014). In mammalian cells the 5s RNP (5S rRNA/uL5/uL18) is one of the precursor particles that accumulates in cells during interrupted ribosome biogenesis (Sloan, Bohnsack et al. 2013). The elevation of the 5S RNP abundance results in binding of the 5S RNP to Mdm2, the ubiquitin ligase for p53, which in turn stabilizes p53 level. The increased level of p53 leads to arrest of the cell cycle (Fumagalli, Ivanenkov et al. 2012).

Ribosomal stress is also known to cause cell cycle arrest in *Saccharomyces cerevisiae* (yeast) (Bernstein and Baserga 2004, Gomez-Herreros, Rodriguez-Galan et al. 2013, Thapa, Bommakanti et al. 2013). However, there is no known p53 ortholog in yeast. How yeast cells are arrested in the absence of p53 under ribosomal stress is not well known. Understanding of the p53 independent mechanism of cell cycle arrest in yeast will shed light on p53 independent mechanism of cell cycle arrest in mammalian cells. This knowledge will be highly impactful in cancer treatment, because most of the cancer cells lack p53. It also seems very likely that the p53-independent mechanisms work in parallel with the 5SRNP-p53 pathway in p53 equipped organisms.

Since, ribosome biogenesis is a major metabolic event; its abrogation will have system wide effects and will probably impact multiple signaling pathways and metabolic processes. High-throughput transcriptome studies like microarray and RNA-seq are popular techniques used for understanding global gene expression changes in response to complex cellular stresses and the signaling pathways responsive to the stress (Gasch, Spellman et al. 2000, Belli, Molina et al. 2004).

Here, we report the results of RNA-seq analysis of mRNA samples collected from cells with and without ribosomal stress. To create ribosomal stress, we expressed L4 from gal promoter in a L4 null background and shifted to glucose for depleting L4 protein. We used Pgal-uL4 strain instead of Pgal-uL43 for gene expression study, since Pgal-uL43 and Pgal-uL4 had similar cell cycle arrest phenotypes described in chapter 2, and several parameters for uL4 has previously been investigated extensively in our lab. Since ribosomal stress also reduces the rate of ribosome formation while the mature ribosomes existing at the time of imposing ribosomal stress, the cellular protein synthesis capacity relative to total protein mass after the cessation of ribosome formation declined during ribosomal stress. No studies have previously been done to differentiate the effects of abnormal ribosome biogenesis process and reduced protein synthesis. To reduce protein synthesis capacity, here termed translational stress on cells, we have depleted translation elongation factor 3 (Tef3), which is essential in translation elongation, but does not affect ribosome biogenesis (Sasikumar and Kinzy 2014) (M. Shamsuzzaman, A. Bommakanti, A. Zapinsky, N. Rahman, C. Pascual, and L. Lindahl, submitted). To understand the differences between early stress and late stress response, we have done time course RNA-seq experiments. Our RNA-seq data analysis, shows that there are some similar and some dissimilar gene expression pattern in cells of these two stress types. Genes involving energy derivation by oxidation and mitochondrial organization have similar pattern between these two stresses whereas, genes involving ribosomal protein, cell wall stress response and oxidative stress response have mostly dissimilar pattern.

## Materials and Methods

### Strains and media

All Yeast strains used in this article were derived from BY4741. In the Pgal-uL4A strain the gene for uL4B (*RPL4B*) was deleted and the gene for uL4A (*RPL4A*) was expressed at its normal chromosomal location from the *GAL1/10* promoter (Pöll, Braun et al. 2009). *Pgal-TEF3* was derived from BY4741 in our lab by replacing its endogenous promoter with the *GAL1/10* promoter using homologous recombination KanMX selection. GFP-Ras2 was subcloned into PRS316(URA3) from pRS315-GFP-Ras2 (Luo, Vallen et al. 2004) (a gift from Dr. E. Bi).

Cultures were grown asynchronously in 1% yeast extract, 2% peptone, 2% galactose (YPGal) at 30°C until mid–log phase (OD600 0.8–1.0, corresponding to 1.5–2 × 10^7^ cells/ml) and then diluted 1:10 into 1% yeast extract, 2% peptone, 2% glucose (YPD). Alternatively, glucose was added to a galactose culture to a final concentration of 2%. Cells were harvested before and at the indicated times after the shift to glucose medium. Three biological replicates were collected for both of the strains at each time point. The graphical experimental schema for sample collection is shown in Fig 1E. Cultures were diluted as necessary with pre-warmed media to keep the OD600 <1.0 using a Hitachi U1100 spectrophotometer (Hitachi High-Technologies Corporation, Japan), which corresponds to approximately 1.5×10^7^ cells per ml in an exponentially growing glucose culture (L. Lindahl, personal communication).

**Fig 1:**
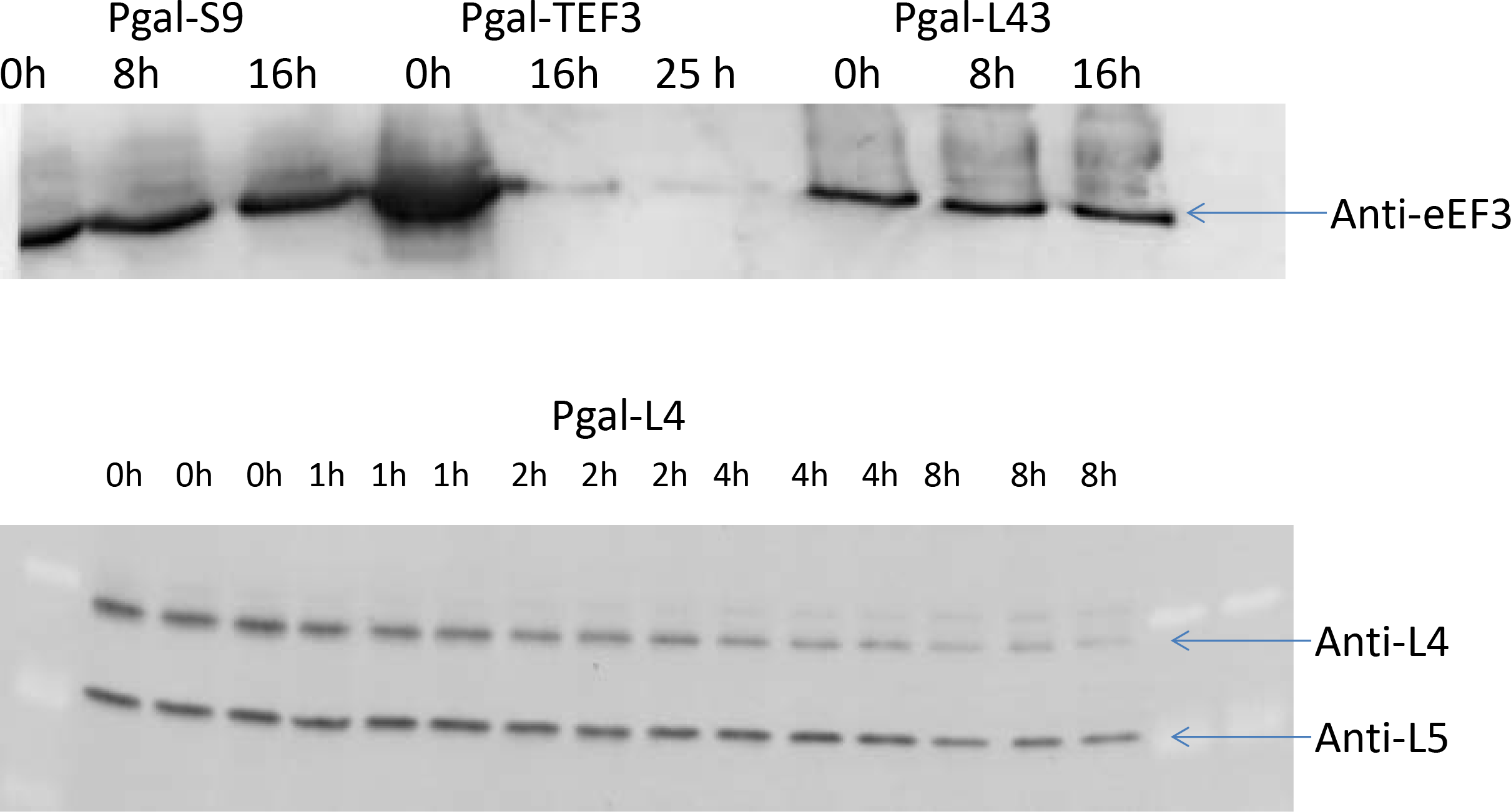
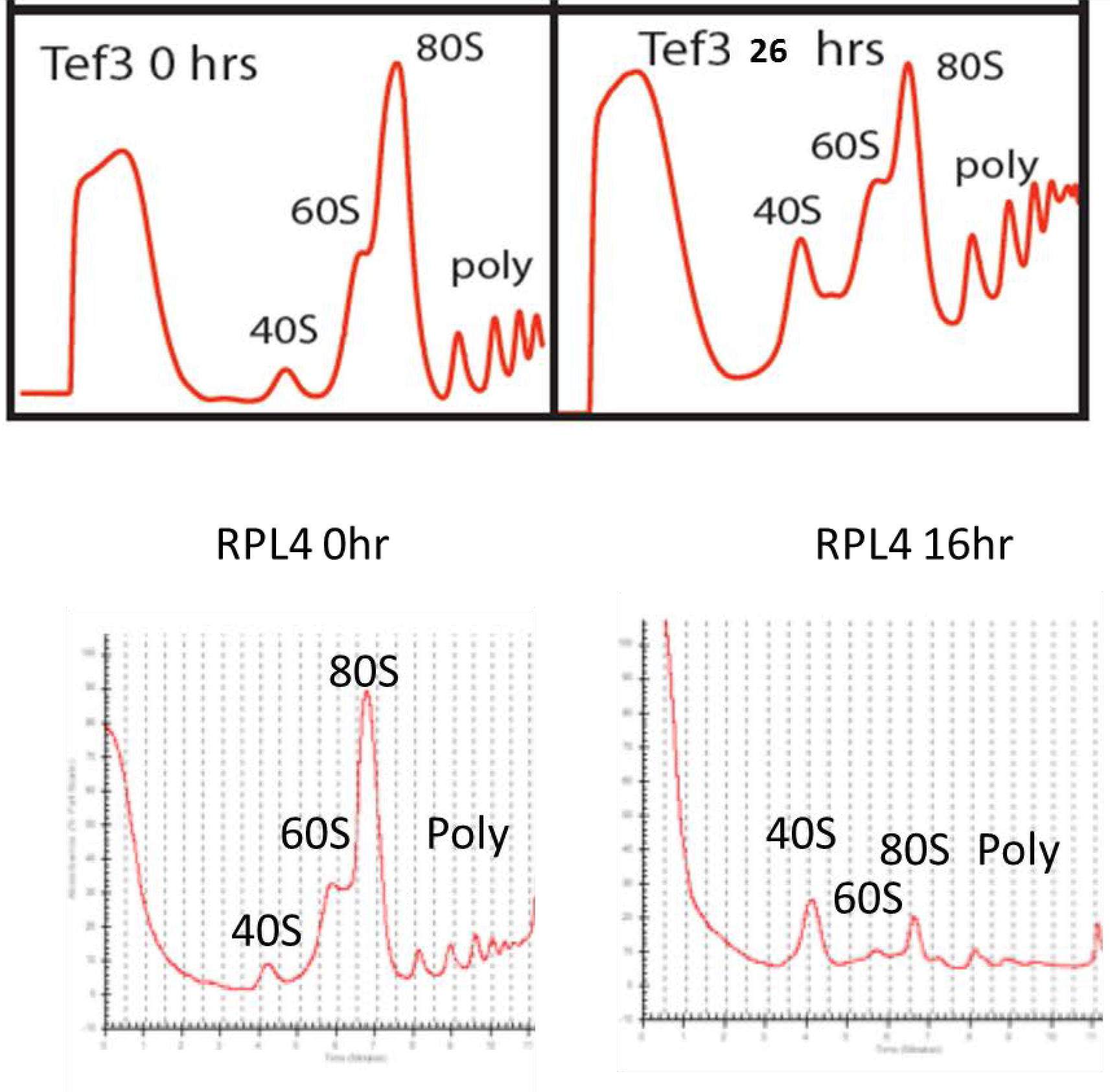
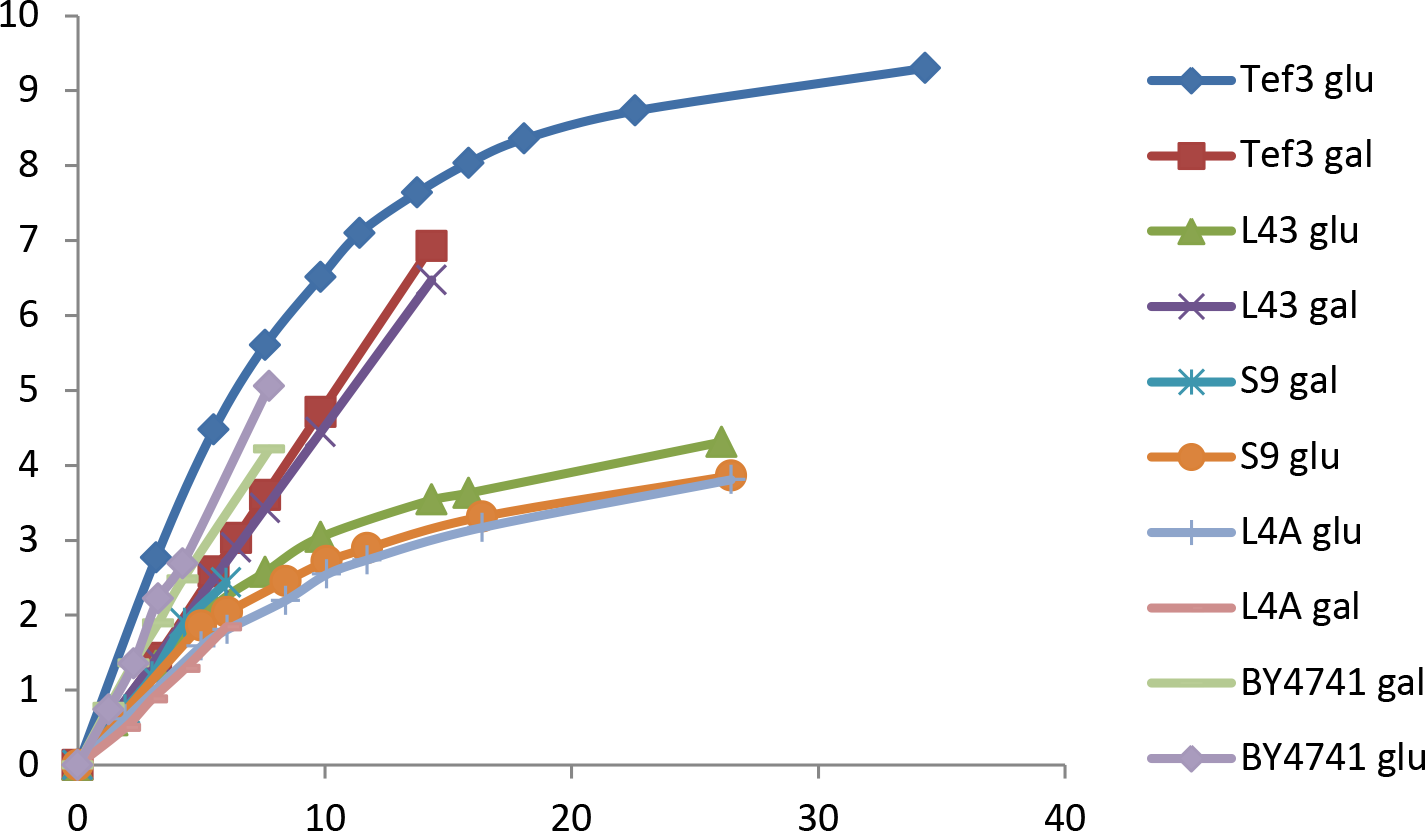
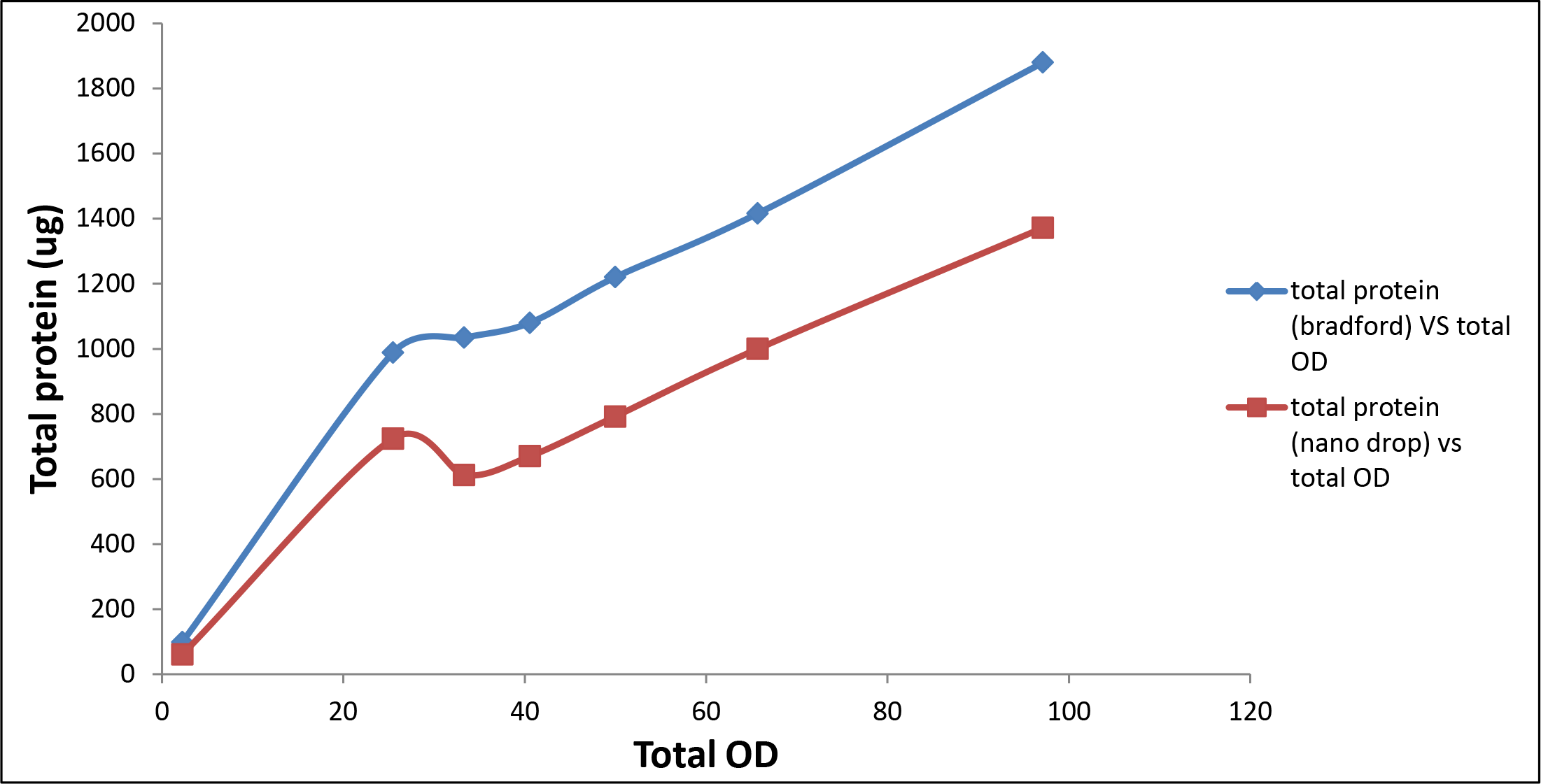
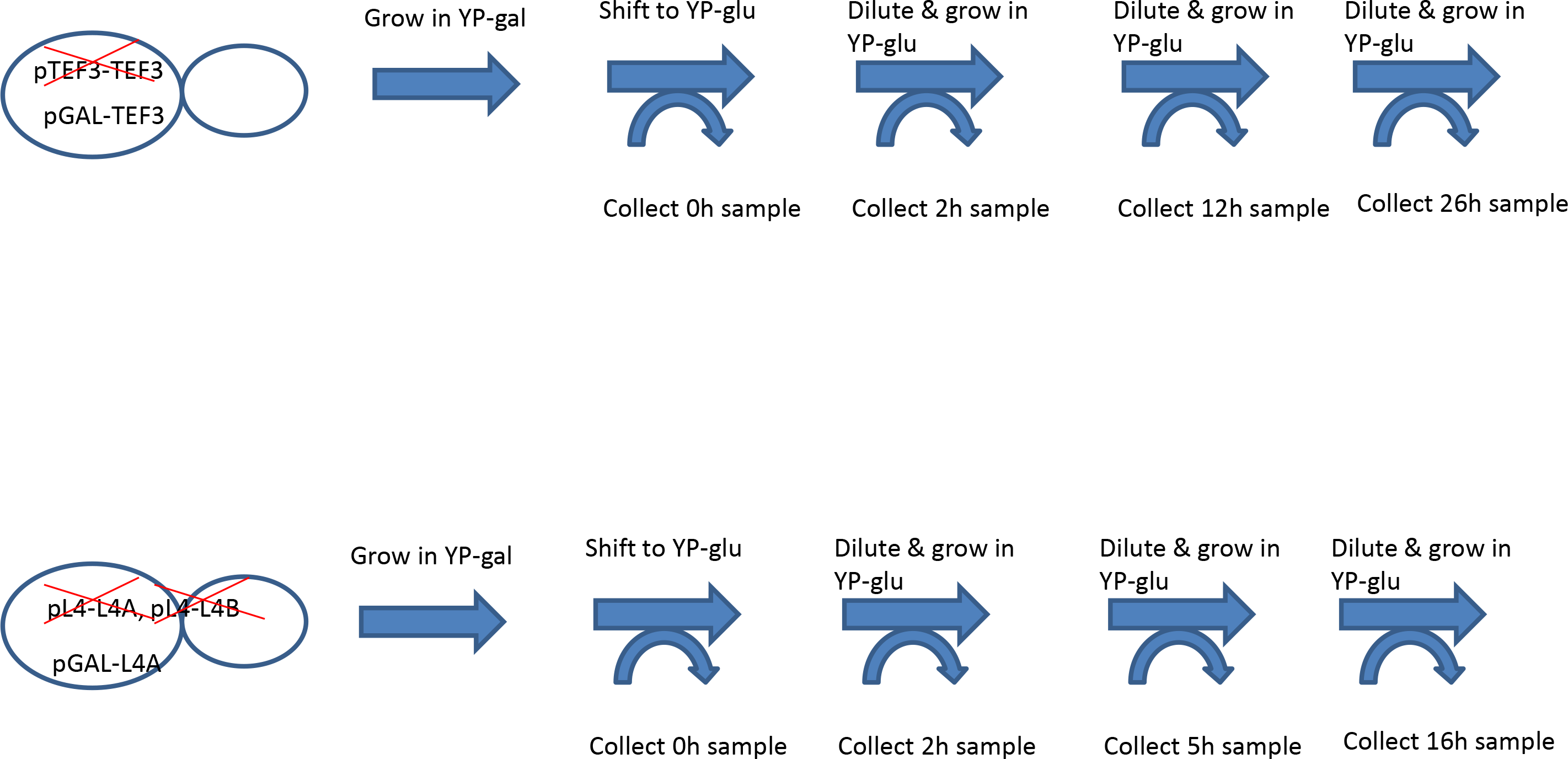
Effect of L4 and Tef3 depletion on growth and ribosome biogenesis. A) Western blots showing depletion of L4 and Tef3 proteins. Total cell lysates from uS4, uL43, and Tef3 depleted samples were run on the upper gel and probed with anti-eEF3 antibody. Tef3 does not significantly decrease in uS4 and uL43 depleted samples. But, only a trace amount of Tef3 protein was detected in Tef3 depleted samples. In the bottom gel, uL4 depleted samples were run in triplicates. By 8hr, most of the L4 protein was observed to be depleted. B) Polysome profiling of samples before and after depletion of L4 and Tef3 proteins. 60S peak is absent and 80S peak is short in L4 16 hr sample. Polysome profile of Tef3 depleted sample shows all 40S, 60S, and poly-ribosome peaks like Tef3 0h sample. 80S ribosomal peak was observed to be shorter in Tef3 depleted sample and concomitant increase in polysome particles was detected. C) Growth curves of Pgal-uL4, Pgal-uL43, Pgal-uS4 and Pgal-TEF3 samples grown in galactose and glucose. Cells with ribosomal protein depletion follow similar growth kinetics. By 6hr of ribosomal protein depletion, cell growth seemed to significantly reduce and deviate from the growth of the cells without ribosomal protein depletion. Interestingly, an increase in growth rate was observed for first six hours after shifting Pgal-Tef3 strain to glucose. By 12hr of Tef3 depletion, cells growth significantly slowed down. D) Relationship of total protein (measured using both Bradford assay and A280) content and cellular growth measured by OD^600^ while growing in galactose media. Cell growth increased linearly with the increase of total protein synthesized. E) Graphical experimental schema of sample collection for the RNA-seq experiment. Samples were collected in triplicates at four different time points over the course of uL4 and Tef3 depletion.

### Western analysis

Western analysis was performed as described previously (Lawrence, Shamsuzzaman et al. 2016) using a polyclonal anti-Tef3 (1:10,000 dilution) from Kerafast (http://www.kerafast.com/p-2206-anti-eef1a-antibody.aspx) and monoclonal anti-L4 (1:10,000 dilution).

### Sucrose gradient centrifugation

Cells were grown in media (YPGal or YPD) appropriate for the undepleted and depleted conditions (see above). Lysates were prepared for sucrose gradient centrifugation essentially as described by (Baim, Pietras et al. 1985), except that cycloheximide was not added to the culture before harvest. Briefly Quick-chilled cells were then spun down and washed with PA Lysis Buffer (0.05M Tris HCl at pH 7.5, 30 mM MgCl2, 0.1M NaCl and 200 μg/mL heparin). The cell pellet was resuspended in 1.25 mL of PA Lysis Buffer. The mixture was transferred to a tube containing 3g of glass beads (0.5 mm diameter) and vortexed eight times for thirty seconds at 4°C interrupted by cooling in ice water for 1 minute, and diluted with 1.2 mL PA Lysis Buffer. The cells were spun in 4°C twice at 10,000 g and the supernatant was transferred to a new precooled tube each time. Twenty A^260^ units of lysate were loaded onto a 10-50% sucrose gradient bed. The gradients were spun at 40,000 rpm for 4 hours at 4°C in a SW40 Beckman rotor. Gradients were collected using an ISCO Foxy Jr. sucrose gradient collector pumping at 1 ml/min.

### Confocal microscopy

Cells were fixed for 30 min by adding 3% paraformaldehyde 150 (final concentration) to one OD600 unit of culture. Cells were then collected by centrifugation at 5000 rpm for 5 min and washed twice with 0.1 M potassium phosphate pH 7. Cells were sonicated before being viewed under a Leica TCS SP5 confocal microscope equipped with LAS AF Lite Software using the 63× oil immersion lens. The filter was set at 488 nm for the excitation and 525 nm for the emission.

### RNA purification

Total RNA samples were extracted from yeast cell culture using Ribopure Yeast (ThermoFisher, USA) kit following protocol provided by the manufacturer (https://tools.thermofisher.com/content/sfs/manuals/1926ME.pdf). Briefly, ~2×10^8^ cells were pelleted by centrifugation. Cells were lysed by vortexing with Zirconium beads (0.5 mm) in presence of lysis buffer, 10% SDS and Phenol:Choloform:Isoamyl alcohol. After centrifugation, the aqueous phase was collected and mixed with binding buffer and 100% ethanol. The mixed solution was passed through the filter cartridge membrane. The filter cartridge membrane was washed with two washing solutions provided with kit. In final step, RNA was eluted from the membrane using elution buffer.

### Library preparation, sequencing and analysis

RNA library preparation and sequencing was performed by the sequencing core facility at the Institute for Genome Science at the University of Maryland Medical School. Briefly RNA samples were checked for quality by running on 1% agarose gel and for high RNA integrity number by Bio-Analyzer (Agilent Technologies, USA). One Tef3 12h depletion sample did not pass quality control and was not sequenced. Other 23 RNA samples were first enriched for mRNA using poly-A attached magnetic beads. Paired end RNA seq libraries were prepared from the enriched RNA samples using TruSeq RNA Sample Prep kit (Illumina). One hundred nucleotides were sequenced using HiSeq platform (Illumina) from both ends of each cDNA fragment following manufacturer instructions. Sequence reads were aligned against the *Saccharomyces cerevisiae* S288c reference genome using Tophat2 aligner (Kim, Pertea et al. 2013). Sequence annotation and read counts per gene were generated using HTseq (Anders, Pyl et al. 2015). Calculation of differential gene expression analysis between any RNA samples of 0hr and any specific hour was done using DESeq2. A gene was termed as differentially expressed gene (DEG) if the false discovery rate (FDR) value was <0.05 and the absolute log_2_(fold change) value was ≥ 1. For principal component analysis Python Sci-kit learn library was used and for all heat maps Python Seaborn library was used.

### Gene ontology enrichment and over-representation analysis

Gene ontology (GO) annotation of Yeast genes was developed by Saccharomyces Genome Database project. GO annotations were accessed from geneontology.org (http://www.geneontology.org/) supported by gene ontology consortium (Consortium 2015).

GO annotation database is constantly updated. GO annotation used in this article had GOC validation date 04/28/2017 and gaf-version 2.0. This version had total 110792 GO annotations of yeast genes and had total of 5791 unique GO terms. GO annotations are of three different types – biological process, molecular function and cellular component.

For over-representation analysis, differentially expressed genes (DEGs) between stress sample of any specific time and control sample (0hr) were mapped to GO terms and for pathway over-representation to pathway terms available in KEGG, Reactome, Wikipathways and YeastCyc using the interface of ConsensusPathDB (Kamburov, Pentchev et al. 2011). Number of DEGs mapped to each term were counted and compared to the counts for all genes of yeast genome. Hypergeometric test was performed and the terms with adjusted p value / q value / False discovery rate < 0.05 were defined as over-represented terms.

For enrichment analysis, all genes detected by RNA-seq were ranked by the Log_2_(fold change) value between stress sample of any specific time and control sample (0hr) and mapped to GO terms. Pathway over-representation analysis to pathway terms available in KEGG, Reactome, Wikipathways and YeastCyc was done using the interface of ConsensusPathDB (Kamburov, Pentchev et al. 2011). The number of DEGs mapped to each term were counted and compared to the counts for all genes of yeast genome. Wilcoxon signed-rank test was performed and probabilities were calculated to find whether any observation of combined difference (log fold change) between genes of two samples belonging to a functional group were by chance. The terms with adjusted p value / q value / false discovery rate < 0.05 were defined as over-enriched terms.

To find the significantly enriched terms between all three time points of L4 and Tef3 depletion, we performed the following analysis. First, we filtered the output list of DESEq2 analysis with Log_2_(fold change) between two samples to exclude the genes with fewer than 50 read counts. The genes in the filtered list were ranked from 1 to remaining number of genes in ascending order of fold change of expression. Then, rank value of each gene at L4 2h sample was subtracted from that of Tef3 2h sample to calculate rank value difference (RVD). RVD was calculated in the same way from L4 5h and Tef3 12h samples and from L4 16h and Tef3 26h samples. Absolute values and actual values of RVDs of each gene were summed over three time points to calculate the absolute rank sum and rank sum, respectively. Finally, absolute rank sum was subtracted from absolute value of rank sum to calculate the rank sign. The genes with highest absolute rank sum values are the genes, which differs most in terms of fold change value between these strains collectively over three time points. The genes with high rank sign values are the genes which had higher rank in L4 relative to Tef3 at a specific time point, but changes to lower rank relative to Tef3 at other time points and vice versa. The genes were ranked according to the absolute rank sum of differences between L4 and Tef3 depletion samples. Wilcoxon signed-rank test was performed and probabilities were calculated to find whether any observation of combined difference (log fold change) between genes of two samples belonging to a functional group were by chance. The terms with adjusted p value / q value / false discovery rate < 0.05 were defined as enriched terms. Python Pandas library was used for this analysis.

The list of significant GO terms received from each analysis is long. To better understand the functional concepts, list of GO terms were summarized visually In Treemap structure based on semantic similarity of GO terms after removing redundant GO terms from this list. Revigo package from Bioconductor was used in this analysis (Supek, Bošnjak et al. 2011).

### Transcription factor (TF) over-representation analysis

Yeastract database was used for TF over-representation analysis (Monteiro, Mendes et al. 2008). Genes with known experimental evidence of expression for a given TF was used in this analysis. List of DEGs up regulated between two samples were used to find enrichment of TF targets where TF is acting as an activator and for down regulated DEGs, TF acting as inhibitor.

## Results

### Growth kinetics and ribosome biogenesis after repressing ribosomal protein and translational elongation factor synthesis

We used yeast strains in which the only gene for a particular protein is transcribed from the *GAL1/10* promoter. These strains are referred to as “Pgal-Protein Name” (Pgal-uL4 and Pgal-Tef3). Changing the carbon source from galactose to glucose represses the gene transcribed from the galactose promoter, which immediately stops the synthesis of mRNA for the gene expressed from the *GAL* promoter. However, since yeast messengers have a half-life of about 20 minutes (and 14 minutes for r-proteins) (Kim and Warner 1983) the synthesis of the protein of interest will decline over approximately one hour. Westerns show that one hour after repressing uL4 synthesis the cell content of uL4 relative to total cell protein is reduced by about 20% and continues to fall for at least 16 hours (Fig. 1A). The content of other ribosomal proteins also declines (Brian Gregory, PhD Thesis, UMBC, 2017), presumably because the repression of uL4 synthesis prevents their incorporation into stable ribosomal particles and free ribosomal proteins are degraded with a half-life of a few minutes (Woolford Jr. and Warner 1991). Thus, the specific translation capacity (translation capacity/total cell protein) after repression of a r-protein declines gradually as the ribosomes existing at the time of the shift to glucose medium continue to synthesize protein, while the biogenesis of new ribosomes is blocked. The translation capacity of ribosomes also declines gradually as Tef3/ribosome changes. Furthermore, there appears to be a pool of Tef3 not associated with ribosomes, which can serve as a reservoir during the initial depletion of the total Tef3 (Sasikumar and Kinzy 2014).

To determine if abrogation of Tef3 synthesis affects ribosome content we compared sucrose gradients of equal numbers of A^260^ units of total cell lysates of Pgal-Tef3 and Pgal-L4 before and after switching from galactose to glucose medium. Twenty six hours after repression of *TEF3* the ribosome level was not affected. As expected, an increase in polysome amount was observed resulting from arrest of mRNA bound ribosomes in the middle of translation elongation step in the absence of elongation factor Tef3. In contrast, the ribosome content in Pgal-L4 was nearly obliterated after 16 hours in glucose medium (Fig 1B). Thus, as expected, depletion of Tef3 did not affect the ribosome content, while the cell ribosome content decreased after repression of r-protein synthesis. The change in ribosome content during repression of ribosome formation is further discussed elsewhere (Brian Gregory, UMBC, Thesis to be submitted, 2017)

Prior to the shift both strains grew at a doubling time of about 110 minutes measured by the increase on OD600 (Fig. 1C). After the shift to glucose Pgal-L4 continued to grow at the pre-shift rate until it slowed at 4-6 hours; by 16hr Pgal-L4 growth was essentially stopped (Fig. 1C). Interestingly, the Pgal-Tef3 culture almost doubled its OD600 growth rate after the shift, i.e. it showed a classic shift up growth phenotype, but after 4-6 hours this culture also slowed and after 26 hour growth of Pgal-TEF3 strain had essentially plateaued (Fig. 1C). Difference in growth kinetics can be partially explained by the depletion kinetics of the corresponding proteins (see above). It is also conceivable that blocking r-protein synthesis prevents a shift up of the growth rate via a mechanism that is not triggered by repressing the *TEF3* gene.

### Introduction to analysis of global impact on cell functions after Inhibiting ribosome formation and ribosome function

Ribosome biogenesis is a major metabolic activity in the cell, which accounts for ~60% of total RNA transcription and more than 20% of total protein synthesis process (Warner 1999). Interruption of ribosome biogenesis will therefore have multiple systemic effects on the cell. To understand how abrogation of ribosome biogenesis affects the global gene expression level we did an RNA-seq experiment after repressing uL4 synthesis (see Materials and Methods for details). To distinguish between early and late effects and cellular adaptation to this ribosomal stress, we collected at 0h (pre-shift), 2h (early), 5h (mid) and 16h (late) time points after shift to glucose.

Abrogation of ribosome biogenesis will begin to reduce the cellular translation capacity shortly after repression (Fig. 1A). To discern which effects of reduced ribosome biogenesis originates from dismantling the ribosome assembly pathways and which are results from the general reduced translation activity we also did RNA-seq after repressing the synthesis of the translation elongation factor Tef3 using Pgal-Tef3. Since we wanted to compare gene expression data at time points between these two strains when translation capacity will be similar, samples from Pgal-Tef3 were collected at 0h (pre-shift), 2h (early), 12h (mid), and 26h (late) after the shift to glucose. The slope of the OD600 growth curve for Pgal-Tef3 at these times corresponded approximately to the slops of the OD600 curve for Pgal-L4 at 0h (pre-shift), 5h, and 16h except 2h sample. We collected 2h sample from both strains to distinguish the effects of glucose shift on gene expression from the effect of depletion. Each sample was collected in biological triplicates.

For convenience, we estimated the protein synthesis rates from the OD600 curves. However, since OD 600 depends not only on cell density, but also on cell size and shape, we first determined if OD600 is proportional to translational capacity. We measured total amount of protein by Bradford assay and A^280^ of crude lysates cleared of debris in Pgal-L4 at different times and compared that with the OD600 curve for the strain growing in glucose. We found a linear relationship between total protein and the OD600 measurements indicating the OD600 curve is an appropriate indicator for comparison of translation capacity between these two strains (Fig. 1D). In the following sections and in all figures, we annotated L4 0h and Tef3 0h sample as pre-shift, L4 2h and Tef3 2h samples as early, L4 5h and Tef3 12h samples as mid, L4 16h and Tef3 26h samples as late stress samples. The design of the RNA-seq experiment is illustrated in (Fig. 1E).

### General features of transcriptome measurement at different times after repressing transcription of L4 or TEF3 mRNA

A total of 773 million reads were generated from 23 samples from the Pgal-L4 and Pgal-TEF3 strains averaging 33.7 million reads per sample. Total number of reads generated was similar from sample to sample and 20 out of 23 samples had more than 30 million reads (Table 1). The percentage of reads that aligned to reference genome ranged from 83 to 86%, and 93 to 97% of the aligned reads mapped to exonic region (Table 1). Reads per kilobase per million mapped reads (RPKM) indicate average higher coverage of the gene features in L4 samples than the coverage of gene features in Tef3 samples (Fig 2A). Alignment summaries of the RNA-seq experiment for all replicates are shown in Table 1.

**Fig 2:**
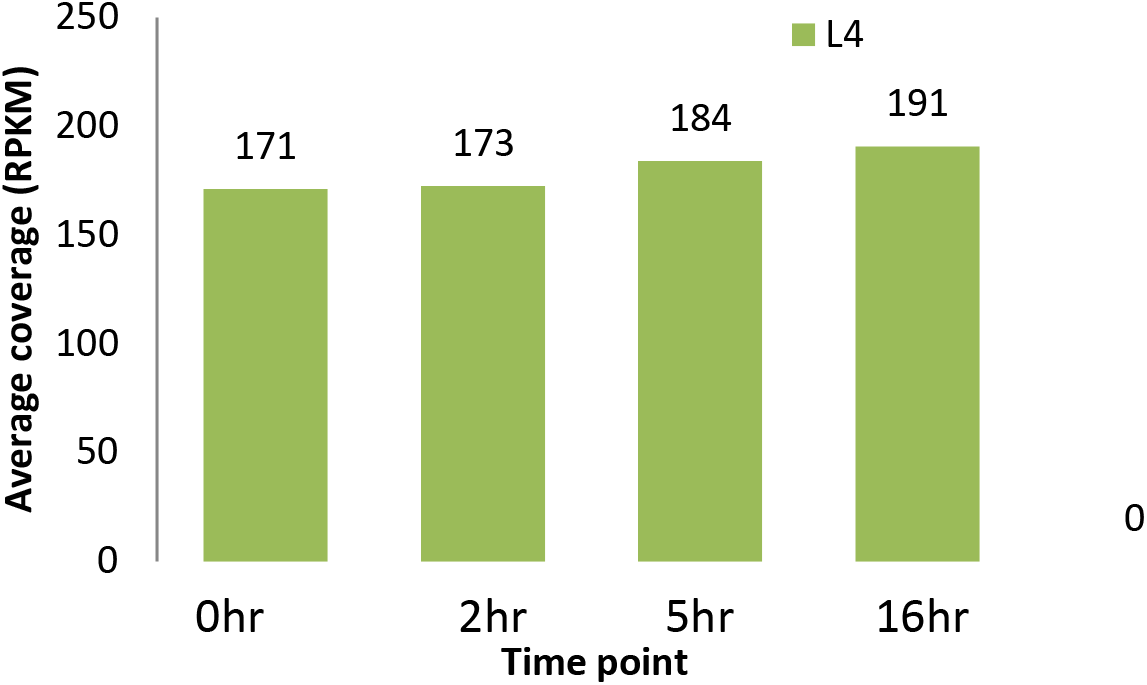
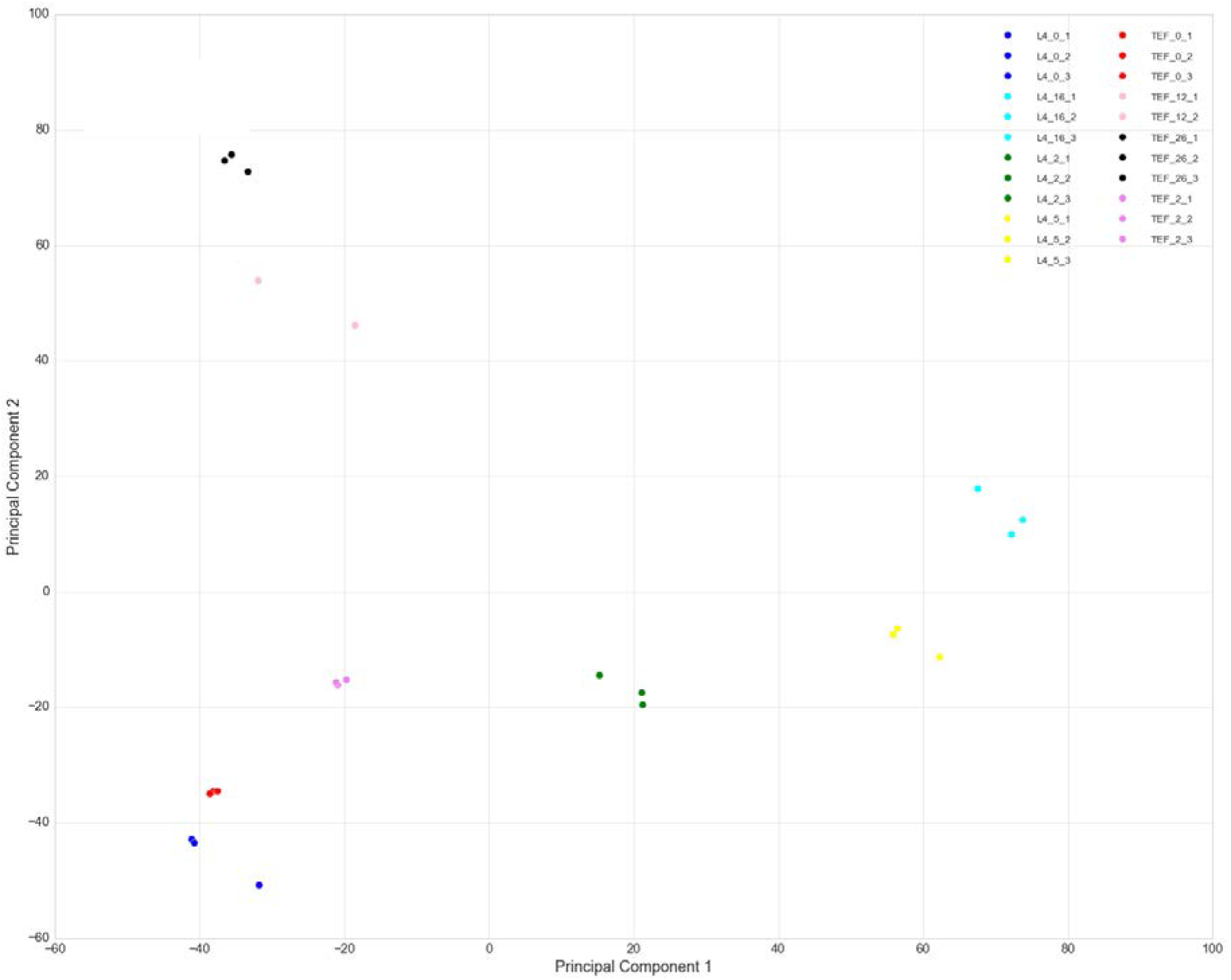
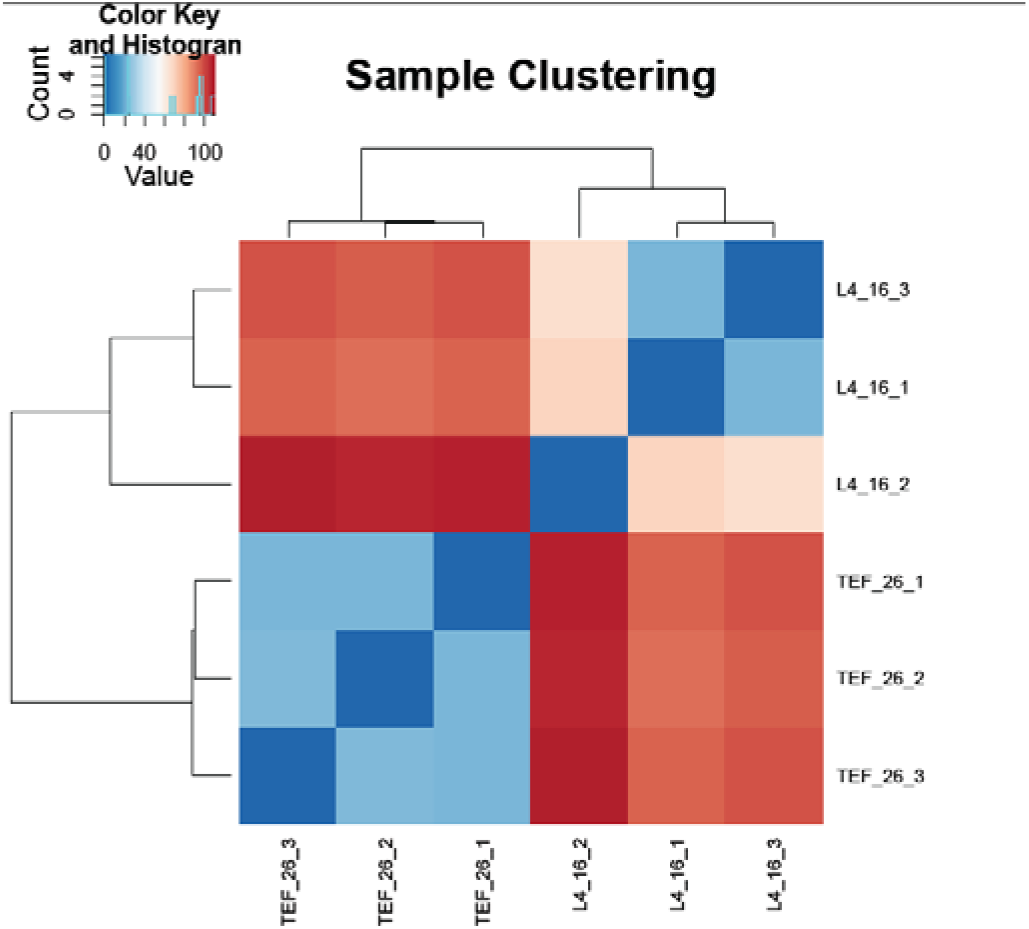
Quality check of the RNA-seq samples. A) Average coverage of transcripts detected in Pgal-L4 and Pgal-TEF3 samples at different points of shift to glucose media. Average coverage is little higher in Pgal-L4 samples. B) Two principal components showing variations in normalized read counts between replicates of samples, variations between samples collected at different time points and variation between two different types of depletion samples. 0hr replicates of both Pgal-L4 and Pgal-TEF3 samples cluster together. At 2hr, samples start to differentiate from each other even though replicates of each samples cluster tightly. Pgal-L4 and Pgal-TEF3 samples collected at 16hr and 26hr respectively differ significantly from each other and are located along two different components. C) Clustering of Pgal-L4 16hr glucose sample replicates and Pgal-TEF3 16hr glucose sample replicates showing again the replicates of each samples form tight and differentiating cluster.

**Table 1:** Alignment summary of RNA sequencing.

| #Sample_ID | Total.Reads | Total.Mapped.<br>Reads | Percent.Map<br>ped.Reads | Percent.Prop<br>erly.Paired | Uniquely.Map<br>ped.Reads | Percent.Ex<br>onic | Percent.In<br>tronic | Percent.In<br>tergenic | Total.Feat<br>ures | Features.With<br>.Coverage | Avg.RPKM |
| --- | --- | --- | --- | --- | --- | --- | --- | --- | --- | --- | --- |
| L4_0_1 | 31160098 | 26091108 | 83.73 | 81.41 | 24843193 | 94.43 | 0.33 | 5.23 | 7126 | 6959 | 175.557 |
| L4_0_2 | 30781140 | 26058135 | 84.66 | 82.07 | 24928504 | 94.81 | 0.28 | 4.91 | 7126 | 6958 | 168.076 |
| L4_0_3 | 26708550 | 22765778 | 85.24 | 84.69 | 21783187 | 94.85 | 0.31 | 4.84 | 7126 | 6938 | 169.921 |
| L4_16_1 | 36991550 | 30506119 | 82.47 | 74.39 | 29355476 | 93.66 | 0.67 | 5.67 | 7126 | 6959 | 183.784 |
| L4_16_2 | 31995934 | 27774929 | 86.81 | 85.99 | 26508361 | 97.01 | 0.39 | 2.6 | 7126 | 6906 | 198.837 |
| L4_16_3 | 36689130 | 30929680 | 84.3 | 80.94 | 29801308 | 93.65 | 0.7 | 5.65 | 7126 | 6973 | 189.542 |
| L4_2_1 | 27886160 | 23686502 | 84.94 | 82.54 | 22762919 | 94.05 | 0.23 | 5.71 | 7126 | 6963 | 170.237 |
| L4_2_2 | 34014132 | 28757250 | 84.55 | 81.47 | 27535637 | 94.11 | 0.23 | 5.66 | 7126 | 6977 | 173.129 |
| L4_2_3 | 31898502 | 26632952 | 83.49 | 80.81 | 25487021 | 94.13 | 0.23 | 5.64 | 7126 | 6990 | 174.199 |
| L4_5_1 | 31329378 | 26146877 | 83.46 | 79.86 | 25140341 | 94.08 | 0.31 | 5.62 | 7126 | 6964 | 180.279 |
| L4_5_2 | 36924836 | 30694342 | 83.13 | 80.25 | 29240834 | 95.05 | 0.27 | 4.68 | 7126 | 6987 | 188.077 |
| L4_5_3 | 35606604 | 30237874 | 84.92 | 81.71 | 29093196 | 94.18 | 0.3 | 5.52 | 7126 | 6973 | 183.353 |
| TEF_0_1 | 37346500 | 32217107 | 86.27 | 83.83 | 30805084 | 95.17 | 0.3 | 4.53 | 7126 | 6973 | 169.067 |
| TEF_0_2 | 34300980 | 29186649 | 85.09 | 81.62 | 27891837 | 95 | 0.31 | 4.69 | 7126 | 6959 | 166.823 |
| TEF_0_3 | 29945526 | 25410700 | 84.86 | 80.91 | 24309318 | 94.98 | 0.29 | 4.73 | 7126 | 6952 | 167.004 |
| TEF_12_1 | 37484454 | 32394050 | 86.42 | 83.63 | 30869484 | 95.55 | 0.25 | 4.19 | 7126 | 6965 | 159.694 |
| TEF_12_2 | 34380570 | 29375261 | 85.44 | 83.37 | 28058017 | 95.5 | 0.28 | 4.22 | 7126 | 6972 | 170.177 |
| TEF_26_1 | 32073470 | 27571486 | 85.96 | 84.9 | 26373958 | 95.32 | 0.21 | 4.47 | 7126 | 6972 | 160.663 |
| TEF_26_2 | 35181456 | 30299713 | 86.12 | 84.08 | 28940924 | 95.27 | 0.21 | 4.52 | 7126 | 6974 | 157.593 |
| TEF_26_3 | 32350742 | 28083916 | 86.81 | 85.17 | 26786000 | 95.36 | 0.17 | 4.47 | 7126 | 6966 | 157.391 |
| TEF_2_1 | 33958280 | 28793497 | 84.79 | 80.05 | 27450692 | 95.48 | 0.25 | 4.27 | 7126 | 6940 | 174.172 |
| TEF_2_2 | 38516970 | 33088402 | 85.91 | 82.63 | 31600268 | 95.28 | 0.3 | 4.42 | 7126 | 6972 | 179.033 |
| TEF_2_3 | 38590280 | 33067634 | 85.69 | 81.98 | 31539775 | 95.17 | 0.29 | 4.54 | 7126 | 6954 | 175.152 |

To determine the variance among biological replicates and samples collected from each strain, we did a principal component analysis (PCA) on the read counts per transcript/gene feature after size factor normalization using DESeq2 package. The replicate samples for a given time point in each strain clustered tightly indicating high reproducibility (Fig 2B). Furthermore, L4 0hr and Tef3 0hr replicates clustered together indicating a strong similarity. Thus the two strains growing uninhibitedly in galactose have very similar gene expression patterns. At 2hr after the shift to glucose the two strains began to differentiate from each other, a trend that was even more pronounced when comparing samples after 5h of L4 depletion with 12h of Tef3 depletion. Comparison of the samples from 16h of L4 depletion with the samples from the 26h time point for Tef3 depletion indicate that gene expression differ significantly from each other after long periods of depletion (Fig 2B).

### Comparative gene expression profile analysis

Differential gene expression analysis of L4 2hr, 5hr and 16hr samples relative to the 0hr sample demonstrated that the gene expression landscape changed over time. At 2hr, 44% of all the genes with FDR (False Discovery Rate) ≤0.05 were in between the +1.5 and -1.5 fold change, meaning that these genes were expressed at approximately the same level at 2 h as they were before the shift (Fig 3A). Fewer than 20% of the genes were either up or down regulated more than 50% at this time. After 5hr depletion, the gene expression landscape did not change that much relative to the 2hr time point either. However, after 16h of L4 depletion, roughly 15% of all genes were up regulated by more than 2.5 fold relative to the 0h values, and 15% of the genes were down regulated more than 2.5 fold. Most of the remaining 70% of genes were up regulated or down regulated between 1.5 and 2.5 fold. Thus the gene expression landscape changes dramatically between 5h and 16h of cessation of L4 synthesis. After depleting Tef3 for 2hr, 70% of total genes with FDR value smaller than 0.05 remained in between +/- 1.5 fold change. After 12hr and 26hr of Tef3 depletion, approximately 50% of genes were between +/- 1.5 fold change. Roughly 3% genes were up-regulated more than 2.5 fold and 10% genes were down-regulated by more than 2.5 fold at both 12hr and 26hr time points (Fig 3A). From these data it appears that the cell responds to ribosomal stress by changing expression of more genes than it does in response to translational stress. In fact, by 16hr of repression of L4 synthesis almost all genes are regulated ≥1.5 fold up or down. A plot of log normalized read counts at 2h vs log normalized read counts at 0h samples and 16h vs 0h samples of L4 depletion indicates similar global expression pattern. Genes which are upregulated (more than 2 fold) at 2hr and 16hr time points ranged from low to high read counts, i.e. the DEG does not depend on the preshift level of expression. However, genes downregulated by at least 2 fold after 16hr of L4 depletion were missing in the high end read counts. At 2hr Tef3 depletion read count points were less scattered in comparison to L4 2hr time point and the up regulated genes with high end read counts were absent in TEF 26hr depletion point (Fig 3B).

**Fig 3:**
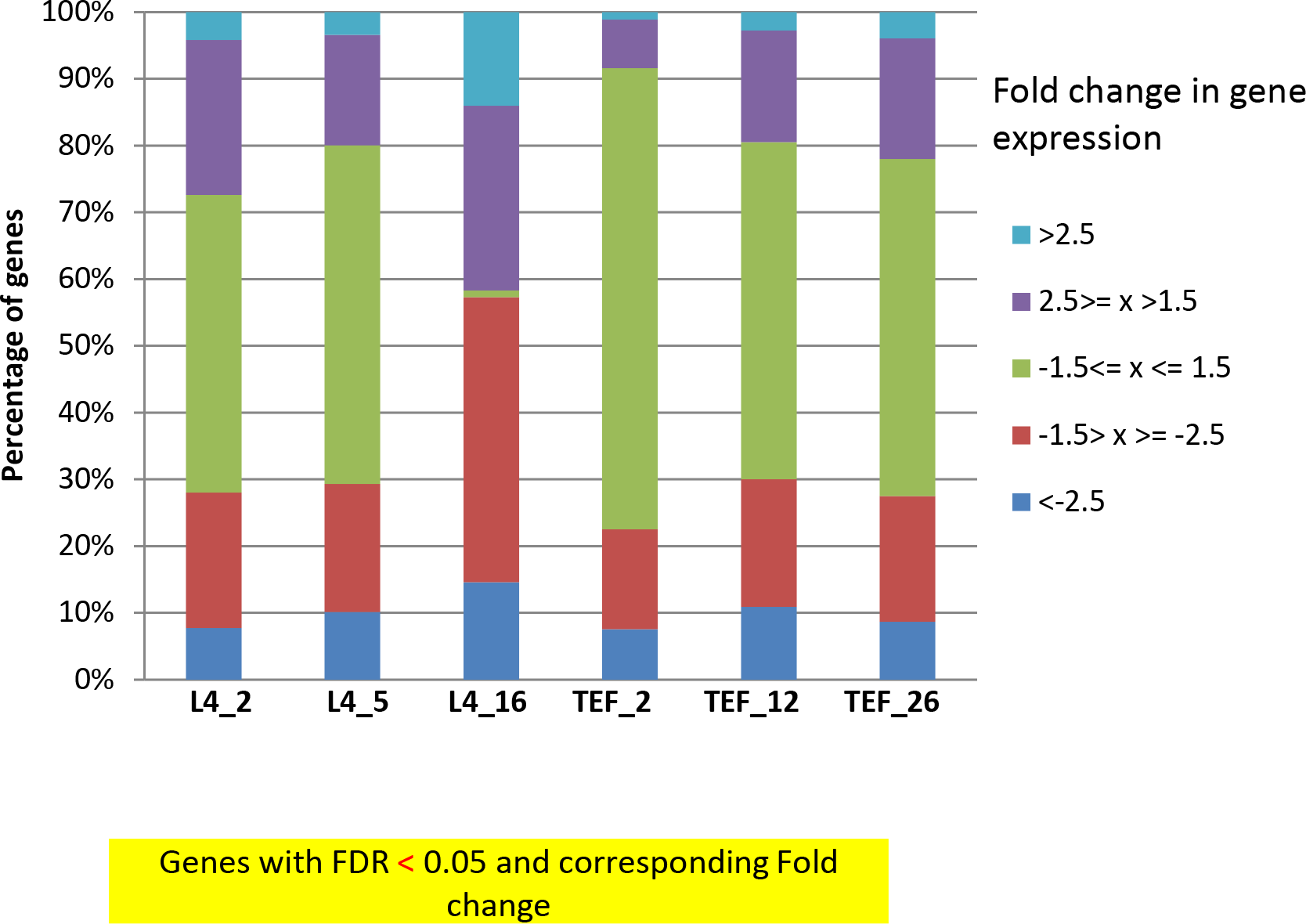
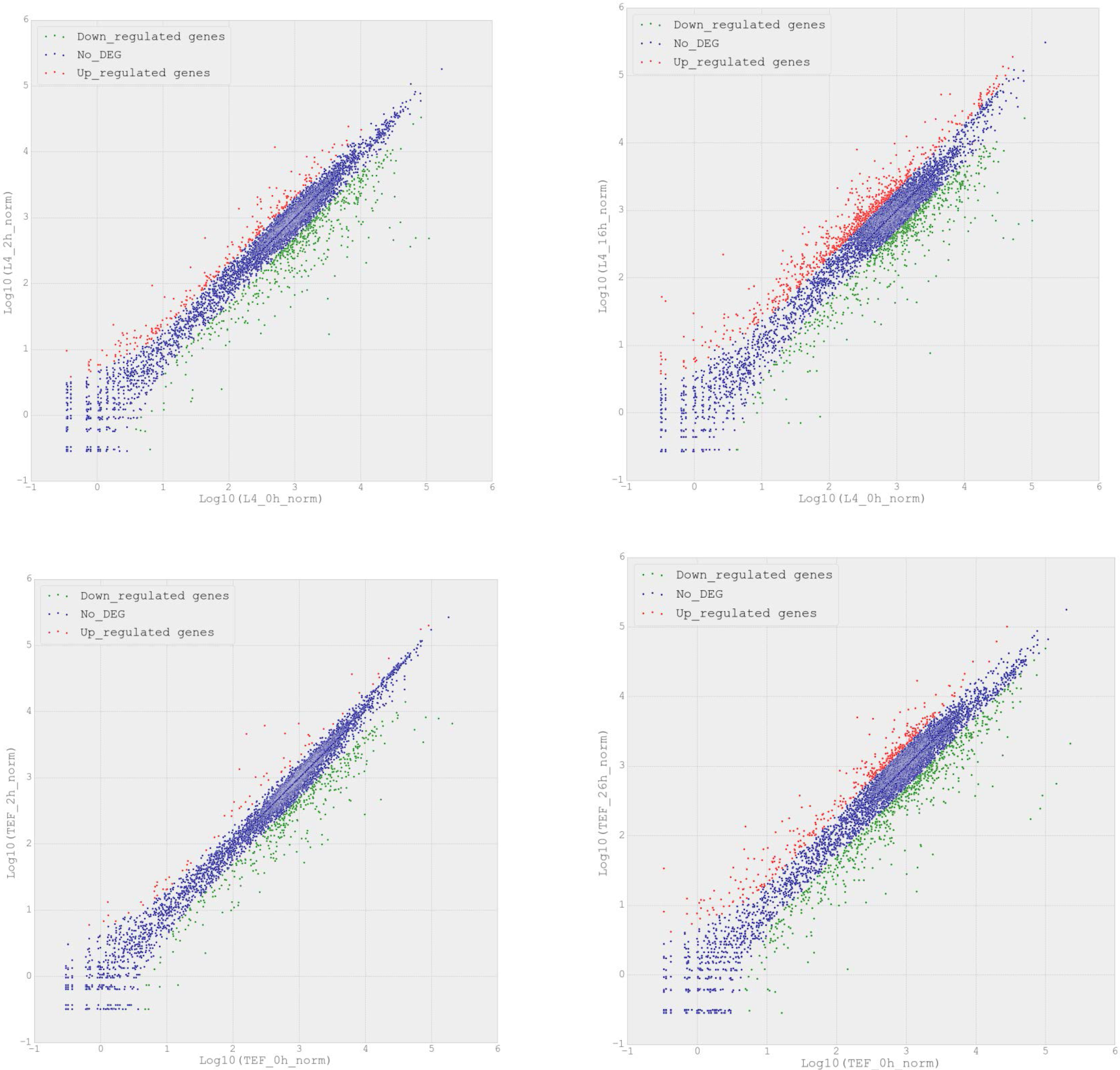
Distribution of gene expression fold changes. A) Bar plots showing distribution of fold changes in gene expression after depleting L4 and Tef3 proteins with respect to without depletion (0hr sample). After 2hr and 5hr depletion of L4, a small portion of genes are up or down regulated by more than 2.5 fold. About 50% of the total genes change less than 1.5 fold at 2hr or 5hr from 0hr sample. After 16hr of L4 depletion, most of the genes are up or down regulated more than 1.5 fold from 0hr sample. Tef3 depletion samples show gradual increase in genes with more than 1.5 fold differential expression than the 0hr sample. However, majority of the genes did not change more than 1.5 fold at any point of depletion. B) Scatter plot showing the distribution of normalized read counts of 0hr sample and normalized read counts of a specific depletion sample. Genes up regulated more than 2 fold are colored as red and genes down regulated more than 2 fold are colored green. Genes with less than 2 fold change are colored as blue. These plots show the distribution of DEGs in terms of read counts. Pgal-L4 16hr sample has more red and green dots implying higher number of DEGs. Read counts of most of the genes including DEGs range between 100 and 10000.

We further analyzed the time course of differentially expressed genes (FR ≤0.05 and fold change ≥2.5). In Pgal-L4 strain, 508 genes were changed at all time points, of which 144 DEGs (Differentially Expressed Genes) are up regulated and 362 DEGs are down regulated at all time points (Fig 4). However, out of total 493 DEGs unique to L4 16hr, majority of the DEGs (341) are upregulated and only 158 DEGs are down regulated. That is, by 16hr time point when the translational capacity has decreased significantly, cells respond by upregulating significantly more genes rather than they repress (Fig 4). We did not observe the same pattern in pGAL-TEF3 strain. Out of total 449 DEGs unique to 26hr Tef3 depletion, 238 DEGs were upregulated and 241 DEGs were downregulated (Fig 4), i.e. almost equal numbers of genes were up and down regulated.

**Fig 4:**
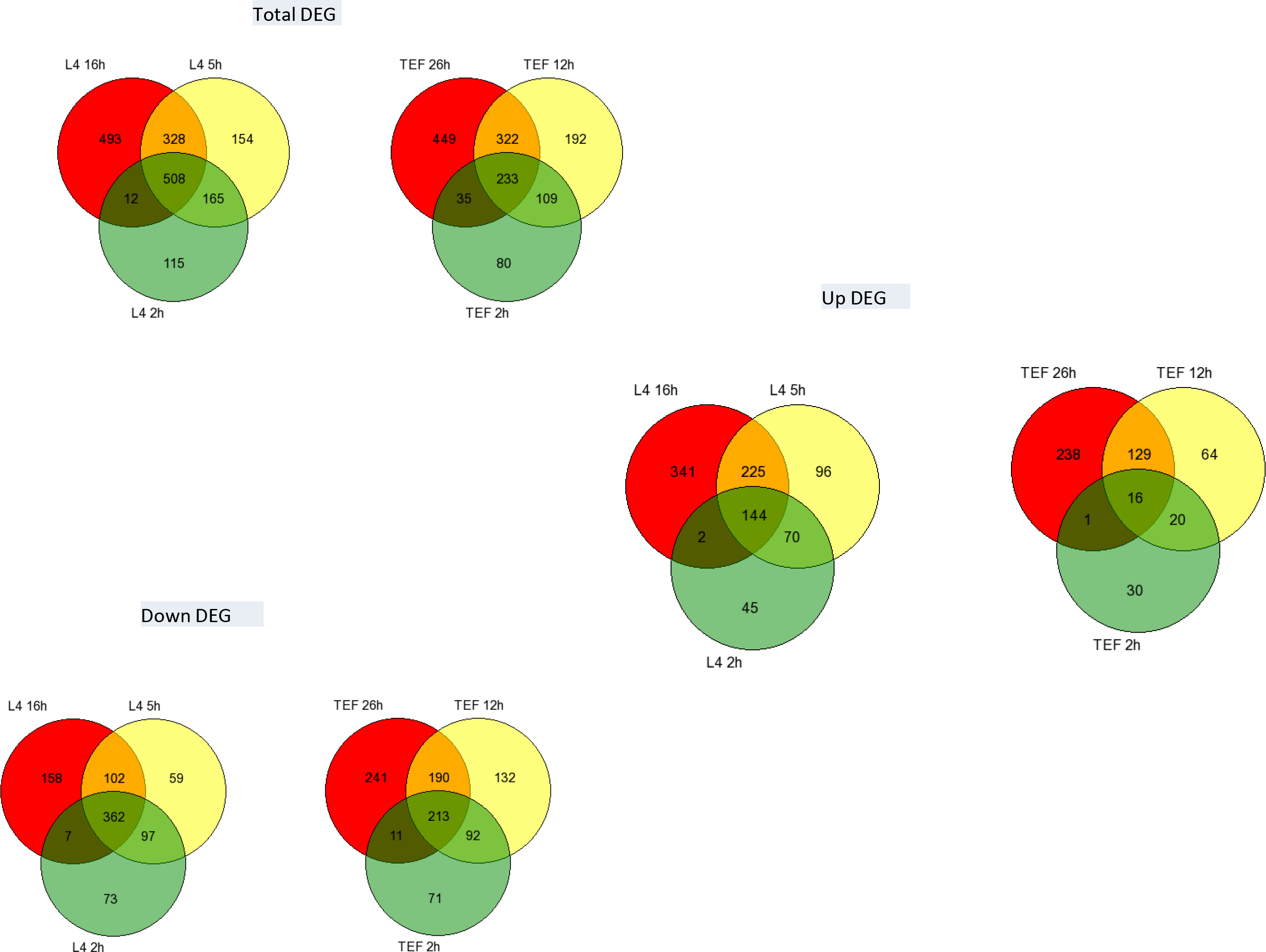
Venn diagram showing number of unique DEGs and common DEGs between samples. Almost twice as many as unique but down regulated DEGs were uniquely upregulated in 16hr uL4 depletion sample. On the other hand, the number of Up and unique DEGs were equal to the number of down and unique genes in 26hr Tef3 depleted sample.

### Enrichment and over-representation analysis of the GO terms

To understand the biological significance of the gene expression changes, we performed Gene Ontology (GO) enrichment and over-representation analyses. First, to determine the difference between early stress and late stress response during uL4 depletion, we performed an enrichment analysis on the 2h and 16h gene expression data using ConsensusPathDB Yeast (http://cpdb.molgen.mpg.de/YCPDB). The enriched GO terms were further analyzed and grouped together based on their semantic similarity using Revigo package. Enrichment treemaps were constructed by grouping related GO terms together and marking them with the same color (Fig 5). The size of each block corresponds to the statistical significance of the GO terms that make up block based on negative log10 of the p values. This analysis showed the functional annotation of the gene expression changes during the stress period. Fig 5A shows an enrichment treemap of GO terms after repression of L4 transcription for 2hr. The strongest regulation was in the groups of “energy derivation by oxidation of organic materials” and “mitochondrion organization”. Although “mitochondrion organization” was also strongly represented after 16h, two other groups not found at 2h appeared. These were “cell cycle”, the most significant group at 16h, and “negative regulation of biological process”. This observation correlates with our prior observation that the cell cycle progression stops in the cells under ribosomal stress (M. Shamsuzzaman, A. Bommakanti, A. Zapinsky, N. Rahman, C. Pascual, and L. Lindahl, submitted), but also suggests that cell cycle genes are not directly affected initially during ribosomal stress. The inclusion of the GO term “mitochondrion organization”, which includes any gene relevant to mitochondrion function and structure and other mitochondrial processes, at both 2h and 16h suggests that cells modify its energy deriving process very early under stress condition in response to sensing the ribosomal stress.

**Fig 5:**
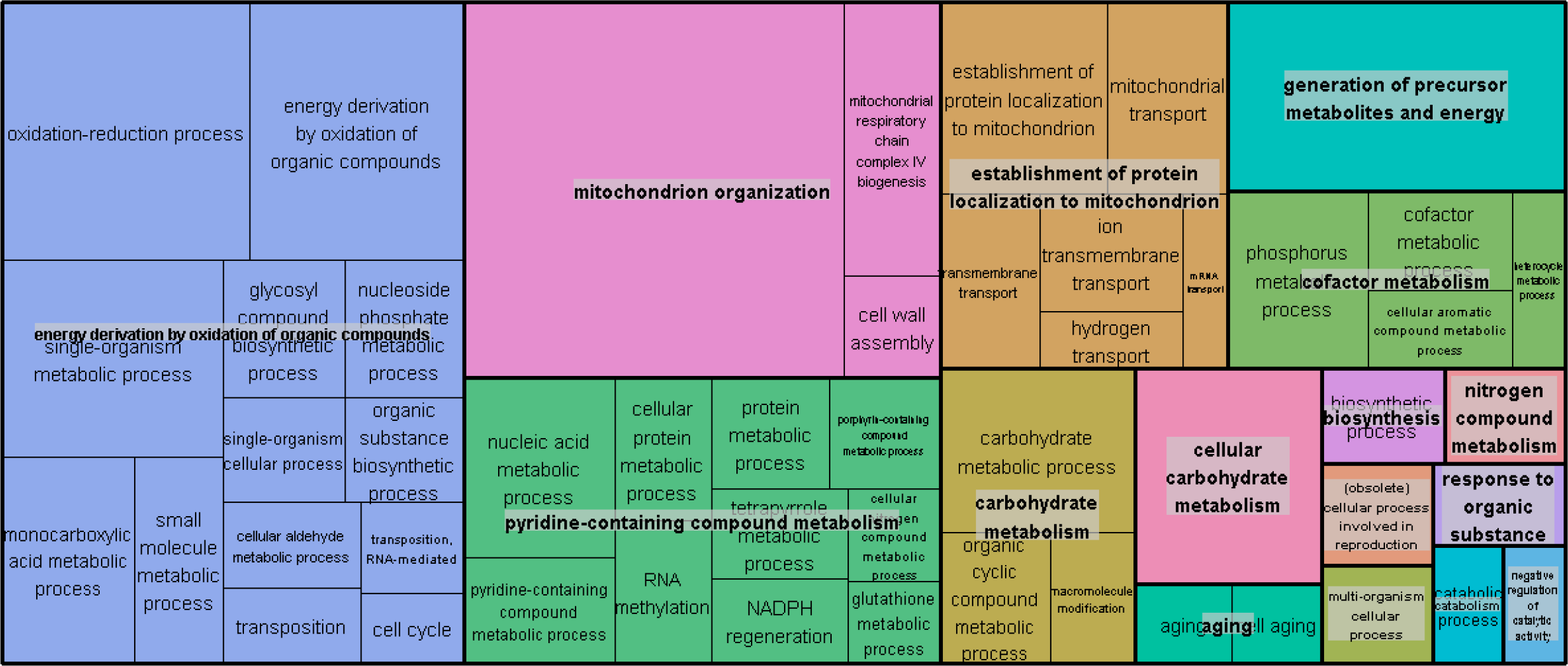
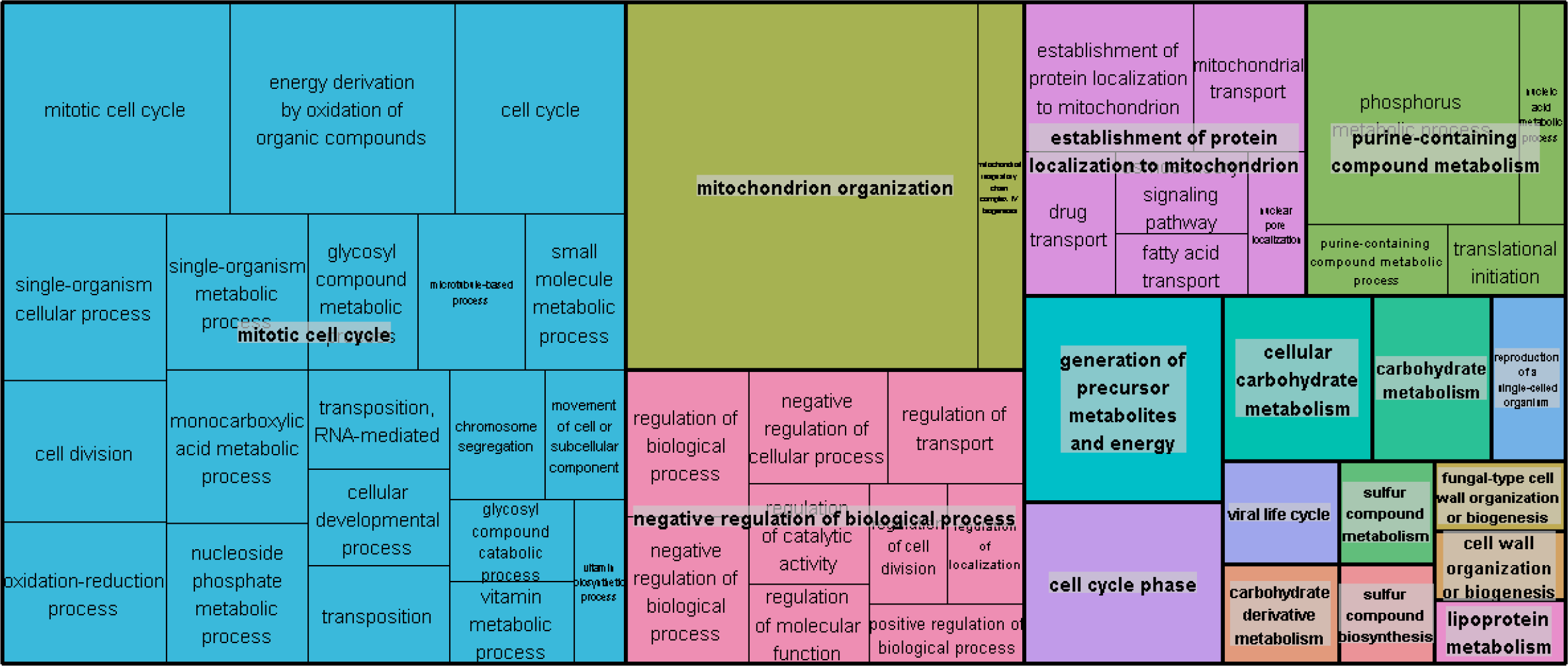
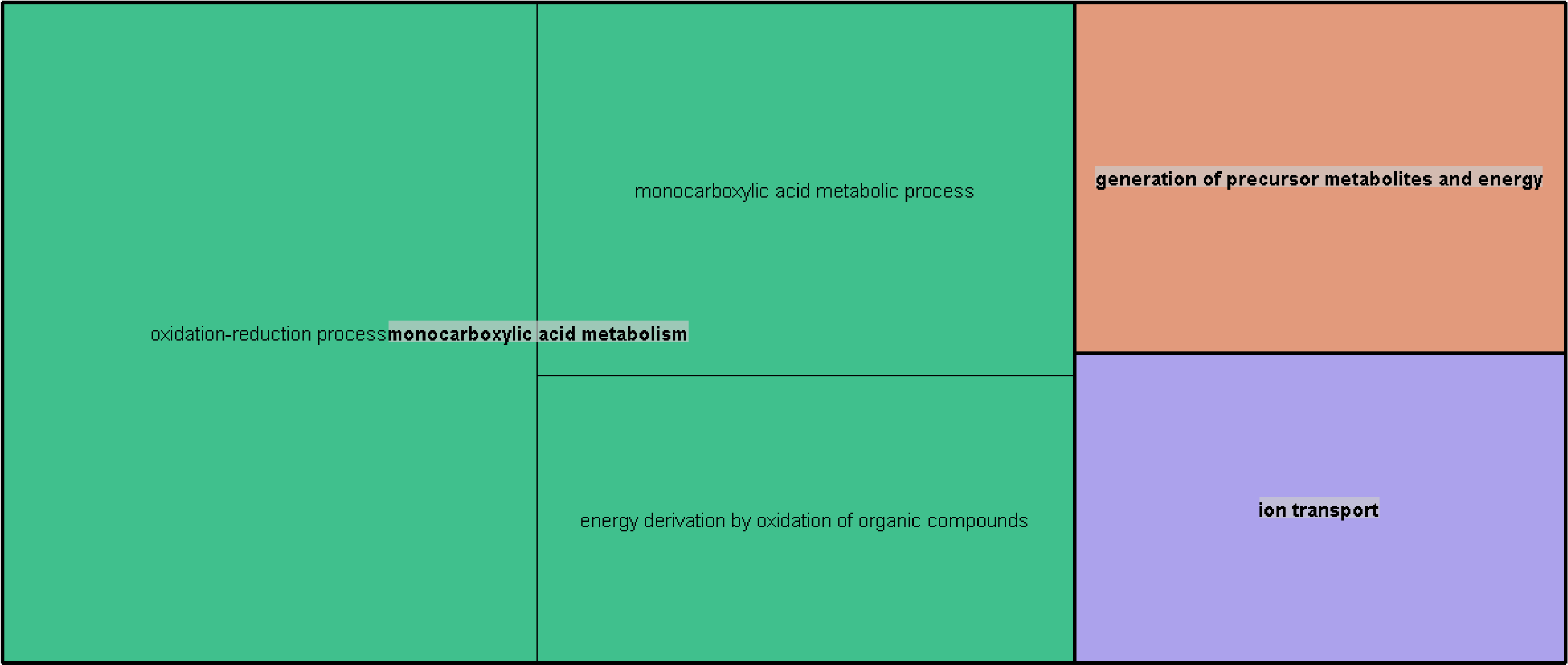
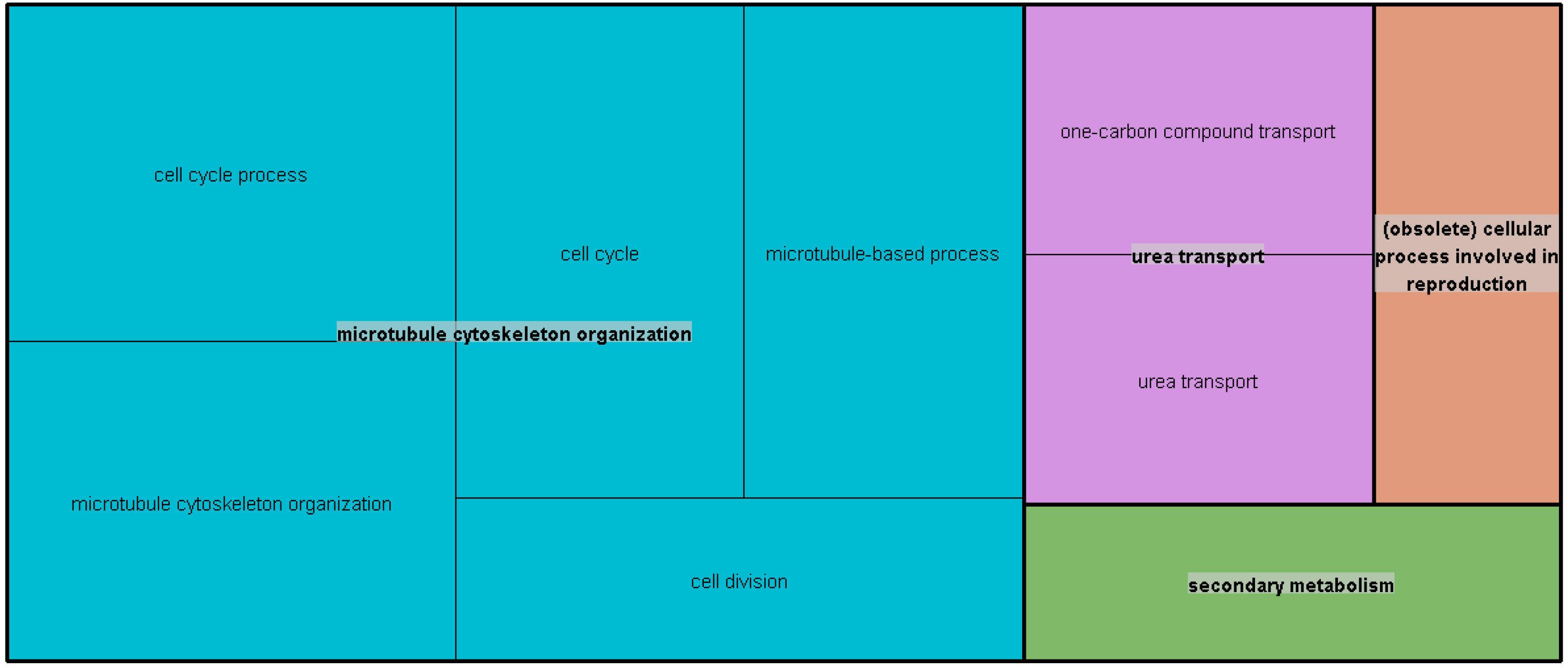
Gene enrichment analysis and over representation analysis of DEGs of early (2hr) and late (16h) L4 depletion samples. A) Treemap showing GO terms enriched in L4 2hr gene expression data with respect to 0hr data. GO terms acquired from gene enrichment analysis are grouped together based on their semantic similarity. The bigger the size of each square, the more significant that term is. Energy derivation and mitochondrion organization are the two most significant terms here. B) Treemap showing GO terms enriched in L4 16hr gene expression data with respect to 0hr data. Mitotic cell cycle and mitochondrion organization are the two most significant terms here. C) Over representation analysis of 115 differentially expressed genes unique to L4 2hr sample. Most significant terms is monocarboxylic metabolism for energy derivation D) Over representation analysis of 493 differentially expressed genes unique to L4 16hr sample. Most significant term here is cell cycle process and associated microtubule cytoskeleton organization.

To further explore the regulation of gene expression during ribosomal stress we did an over-representation analysis of the 115 and 493 genes that were uniquely overexpressed at 2hr and 16hr after abrogation of uL4 synthesis, respectively (Fig 5C and Fig 5D). This analysis also showed the genes involved in cell cycle process and cell division are over-represented in the DEGs list unique to 16hr of stress, which is coherent with the observation from enrichment analysis. Treemaps for over-representation analysis of total DEGs expressed at 2h and 16h after L4 depletion does not show much difference because of prevalence of highly significant similar terms between these two samples (Supplementary Fig 1 and Supplementary Fig 2).

### Comparison of ribosomal stress and translational stress

#### uL4 depletion for 16 hours compared with Tef3 depletion for 26 hours

We next focused on understanding the difference between ribosomal stress and translational stress. A total of 496 DEGs are shared after 16h of repression of L4 synthesis and 26h of Tef3 synthesis (Fig 6). Of these 136 are up regulated, while 315 are down regulated. Out of the 845 DEGs unique to the Pgal-L4 16h sample, 576 DEGs are upregulated, whereas only 248 DEGs unique to the Pgal-TEF3 26h sample are up regulated. Similarly, 314 DEGs unique to L4 16h and 340 DEGs unique to Tef3 26 h are down regulated (Fig 6). Enrichment analysis of GO terms for Tef3 2h (Supplementary Fig 3) and 26h sample (Supplementary Fig 4) showed that “vesicle mediated transport” and “cellular macromolecule metabolism” terms are enriched in the Tef3 26h sample which is different from the enrichment of “mitotic cell cycle” and “negative regulation” of biological process enriched in L4 depletion sample (Fig 5B). In both of the samples, genes affiliated with mitochondrion organization are enriched, indicating that the adjustment of energy derivation of cell most likely occurs in response to decreased translational capacity. However, since “energy derivation by oxidation of organic compounds” appear in both L4 and Tef3 depletion samples at 2h, it is likely that this response is observed due to shift from galactose to glucose. It is known that yeast cells shifted to glucose from non-favorable media upregulate glycolysis and repress TCA cycle and electron transport genes (Rolland, Winderickx et al. 2002).

**Fig 6:**
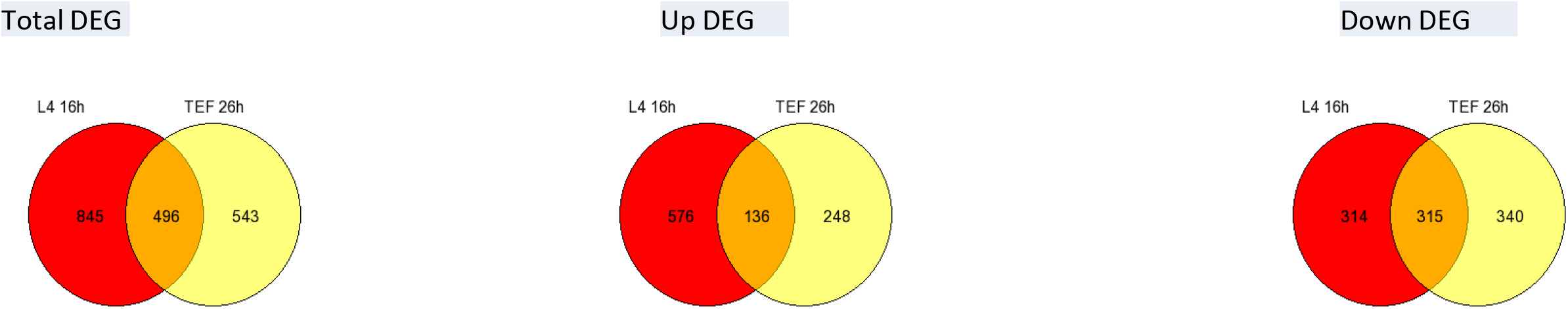
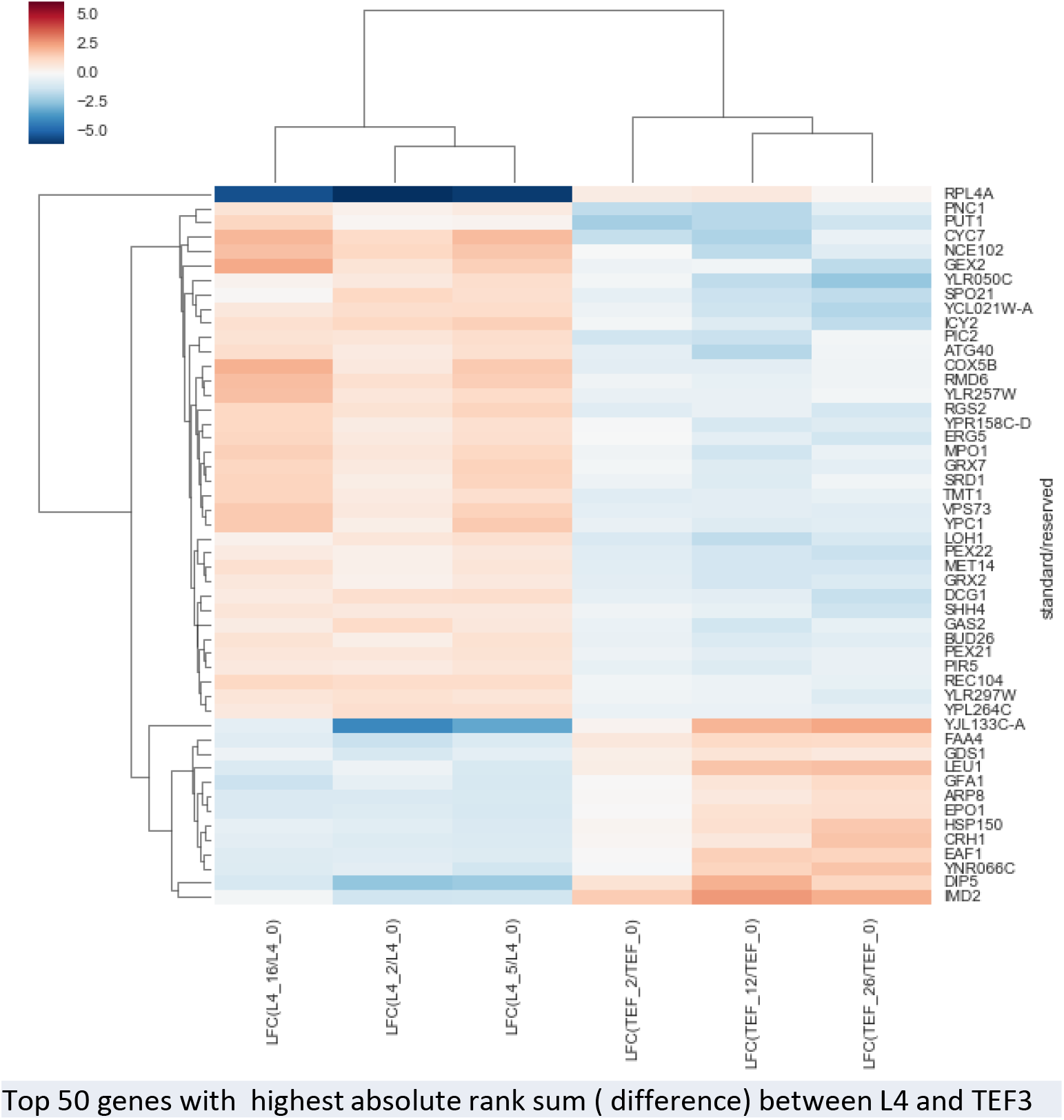
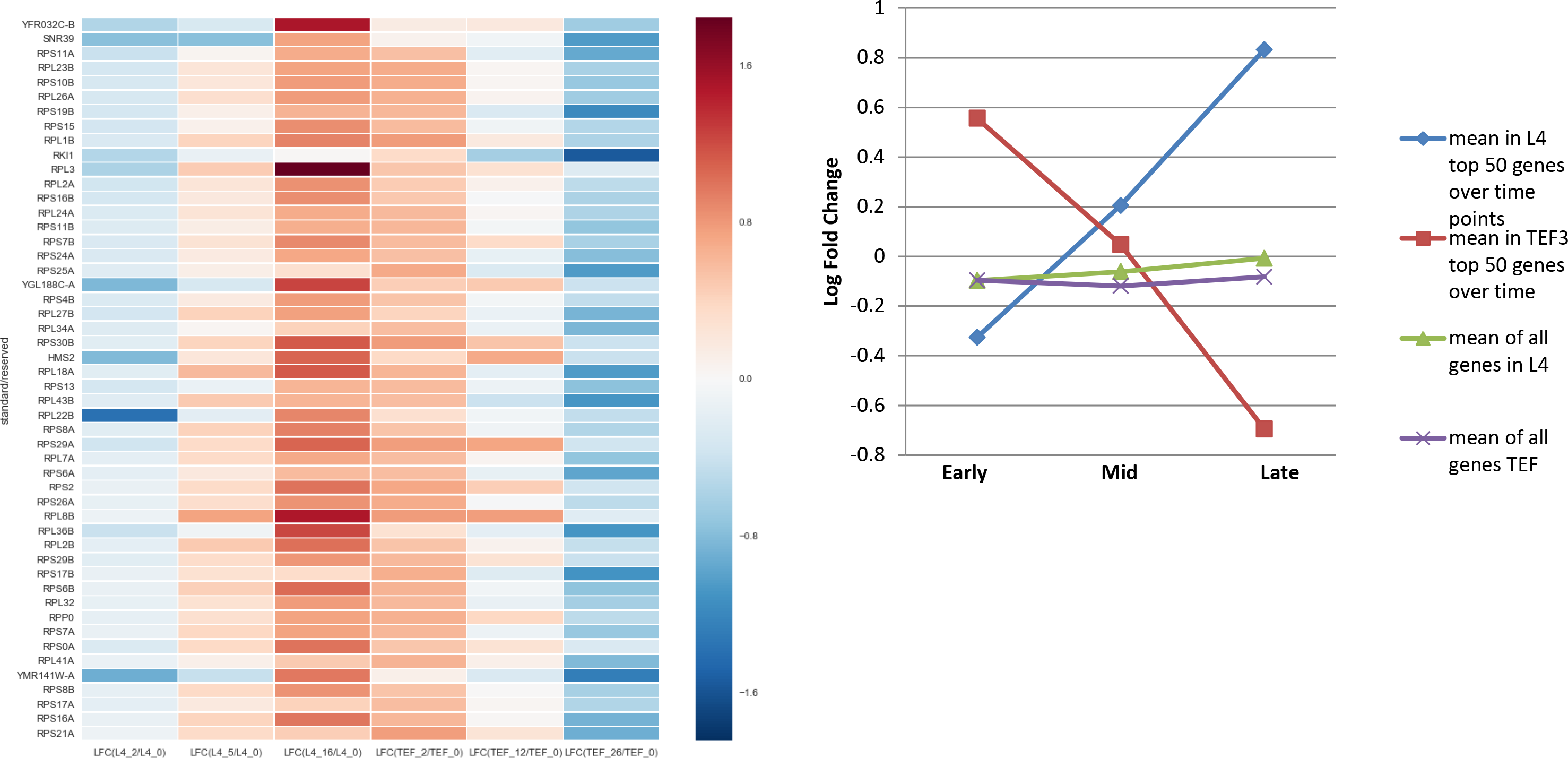
Genes differentially expressed between L4 and Tef3 depletion samples. A) Venn diagram showing unique and common DEGs in L4 16hr and Tef3 26hr samples. DEGs uniquely up regulated in L4 16hr samples are more than double than it is in Tef3 26hr sample. B) Heatmap showing expression of top 50 genes which have highest sum of absolute rank differences (cumulative difference) in expression between these two samples. C) Heatmap and scatter plot showing expression of top 50 genes which have highest sum of absolute rank differences (cumulative difference) in expression between these two samples and were first up in Tef3 early depletion sample but went down in Tef3 late depletion sample.

#### Time trends of Differential Gene Expression

The comparison of gene expression for similar degrees of the change in the slope of the OD600 curves for repression of L4 and Tef3 synthesis did not capture the changes in gene expression over time. As it is clear from GO Tree maps in Fig 5, Supplementary fig 3 and 4; the stress response evolves with time after repression of either L4 or Tef3 synthesis. This may have several causes, including the fact that translation capacity after repression of either L4 or Tef3 declines gradually. Furthermore, it is likely that the true stress response evolves over time, as the cells sense the effects of actin depolarization and pileup of cells in G1 and in mother-daughter complexes (M. Shamsuzzaman, A. Bommakanti, A. Zapinsky, N. Rahman, C. Pascual, and L. Lindahl, submitted). The gene expression pattern during stress also changes with time after imposition of some relatively simple environmental stresses, such as adding a final concentration of a noxious chemical. This often provokes an immediate change in the gene expression pattern which then gradually fades over time (Gasch, Spellman et al. 2000). Furthermore, the shift from galactose to glucose, which is the basis for repressing the genes of interest, also invokes a shift-up response (Kief and Warner 1981), which results in relatively quick changes in gene expression and growth curves, which could complicate the analysis of the initial stress response.

For these reasons, it is important to try to isolate the changes in gene repression that can be attributed to the stress rather than the change in carbon source. Accordingly, we compared the gene expression patters at different times after the shift and ensuing repression of L4 or Tef3 synthesis. If we find a set of genes which has a positive log fold change of expression in response to glucose shift at 2hr time point, but changes to negative trend upon continued repression of uL4 or Tef3, or vice versa, the change of trend is a marker of differential expression due to stress. Furthermore, the differences of gene expression between L4 and Tef3 depletion will be informative, since both strains grow similarly in galactose and are shifted to glucose in similar way. Differences between the two strains can thus be attributed to the stress response, even though the response is different between repression of L4 and Tef3 repression as demonstrated by the Principle Component Analysis (Fig 2).

To accomplish these comparisons, we have calculated sum of absolute rank differences between L4 and Tef3 in early, mid, and late stress samples as described in Materials and Methods. We first identified the genes that differ most in terms of log fold change between these two strains over the time of depletion of L4 and Tef3 proteins. We sorted the list of genes in descending order of sum of absolute rank differences (cumulative difference) and took the top 500 genes. This set of genes are the most likely to have significant difference in log fold change value between the two strains at all three time points and is referred to as list of “top differentiating genes” onward from here. We then used the 50 genes with the highest sum of absolute rank differences (cumulative difference) from this list to generate a cluster-map between two samples (Fig 6B). The difference in gene expression is easily appreciated by looking at the difference of color in the L4 and the Tef3 side of the heat map. Interestingly, L4 2h sample forms a sub-cluster with L4 5h sample; but Tef3 12h sample forms sub-cluster with Tef3 26h sample instead of Tef3 2h sample. A quick look up of top 50 genes (Fig 6B) reveals that the genes which are up in L4 samples but down in Tef3 samples comprises of several oxidative stress responsive genes; whereas the genes which are up in Tef3 but down in L4 contains few cell wall stress genes.

We noticed a shift up response in Pgal-TEF3 strain in the growth curve but not in Pgal-L4A strain (Fig 1). We wanted to understand the difference in growth between these two strains in terms of gene expression. We thought that the genes which are up regulated at Tef3 2hr samples due to shift up response will probably go down as growth slows down and stress builds up. That is why, we next queried for genes, which are most different in terms of sum of absolute rank differences between two strains and also had positive log fold change in expression at 2hr of Tef3 depletion, but changed to negative trend (down regulated) at later time points of stress. Unsurprisingly, we noticed that most of the top 50 genes which met the above condition are ribosomal protein genes (Fig 6C). These genes were up regulated at 2hr after Tef3 repression, but down at 2h after L4 repression (Fig 6C). Surprisingly, during the stress period the expression of the r-protein genes changed strikingly in opposite direction. We compared the mean log fold change for 50 genes shown in Fig 6C with the average DEG for all genes for which significant read counts was generated (scatter plot in Fig. 6C). It is clear that from the plot that there is not a specific positive or negative trend in mean expression of all genes, while the genes shown in the heat map in Fig 6C went from positive to negative trend in Pgal-TEF3 and reverse in Pgal-L4 strain (Fig 6C).

Next, we did an enrichment analysis of all genes and identified GO terms significantly different between L4 and Tef3 samples by ranking the genes according to the sum of absolute rank differences and performing Wilcoxon rank test. Summary of significantly enriched GO terms is shown as treemap in Fig 7. Mitotic cell cycle, protein transport, fungal type cell wall biogenesis are some of the significant GO terms which are differentially enriched between these two strains. To understand the function of the top differentiating genes between L4 and Tef3 samples, we did an over-representation analysis of these genes using CPDB yeast. GO terms significantly over represented in the list of top differentiating genes consists mostly ribosome biogenesis, vesicles, fungal type cell wall biogenesis (Supplementary file 1). We then identified the gene set associated with these GO terms; in most cases from GO annotations of genes and in some cases from studies which attributed genes to a specific function or biological process; generated separate heat maps for the genes included in these GO terms to illustrate how expression changes over time under these two different types of stress (Fig 8-10). We excluded any ‘Inferred from electronic annotation’ (IEA) GO annotation of genes from trend analysis, since this is the least reliable annotation method.

**Fig 7:**
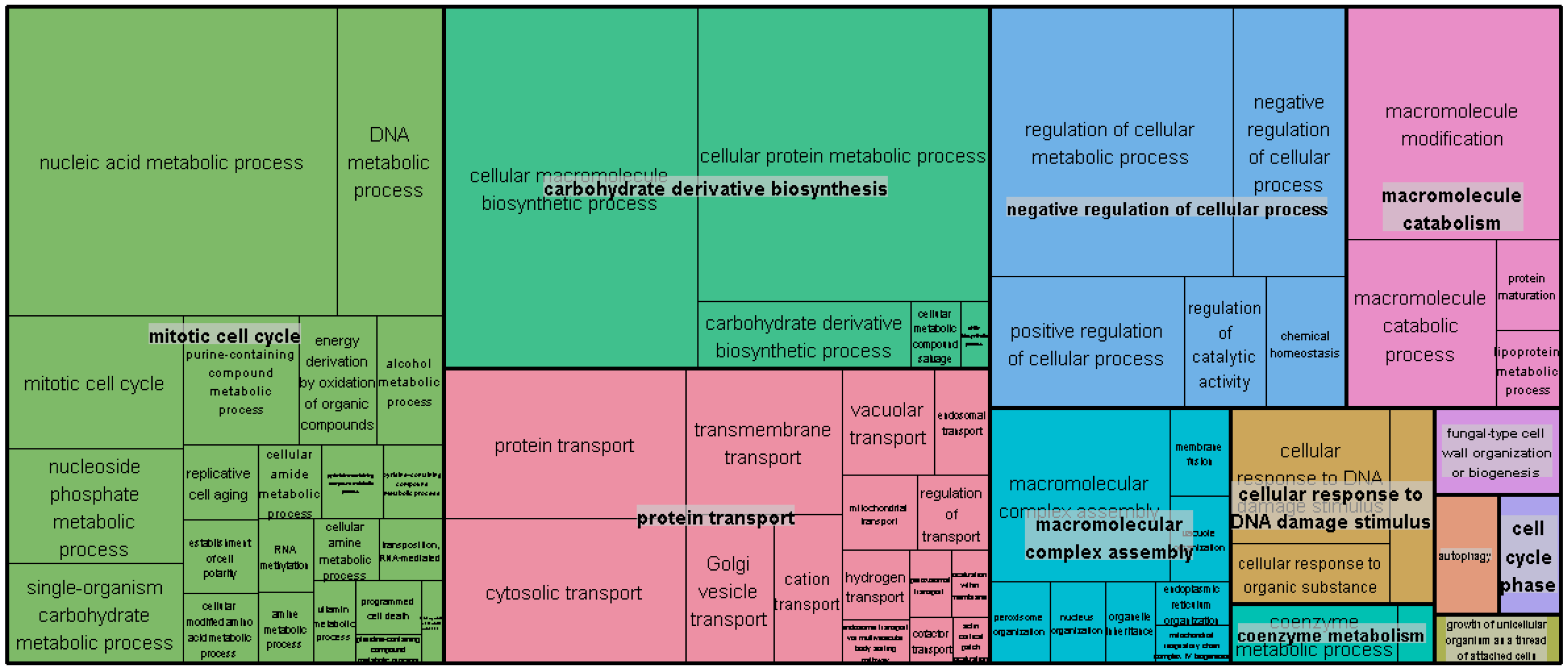
Treemap summarizing significant GO concepts from enrichment analysis of genes based on sum of absolute rank differences between L4 and Tef3 time course depletion samples. Among the GO terms differentially represented between uL4 and Tef3 depletion samples, mitotic cell cycle and protein transport are noteworthy.

**Fig 8:**
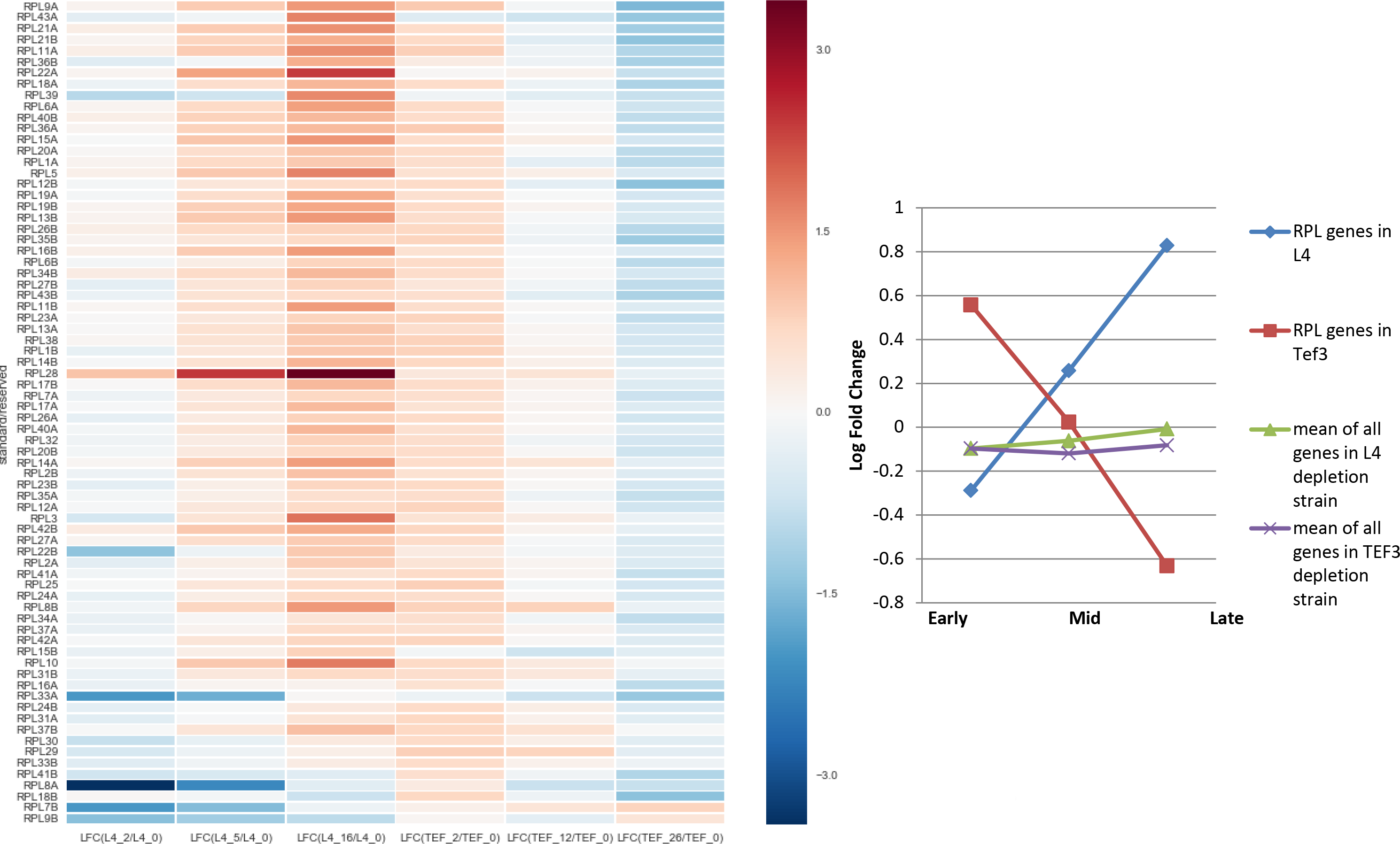
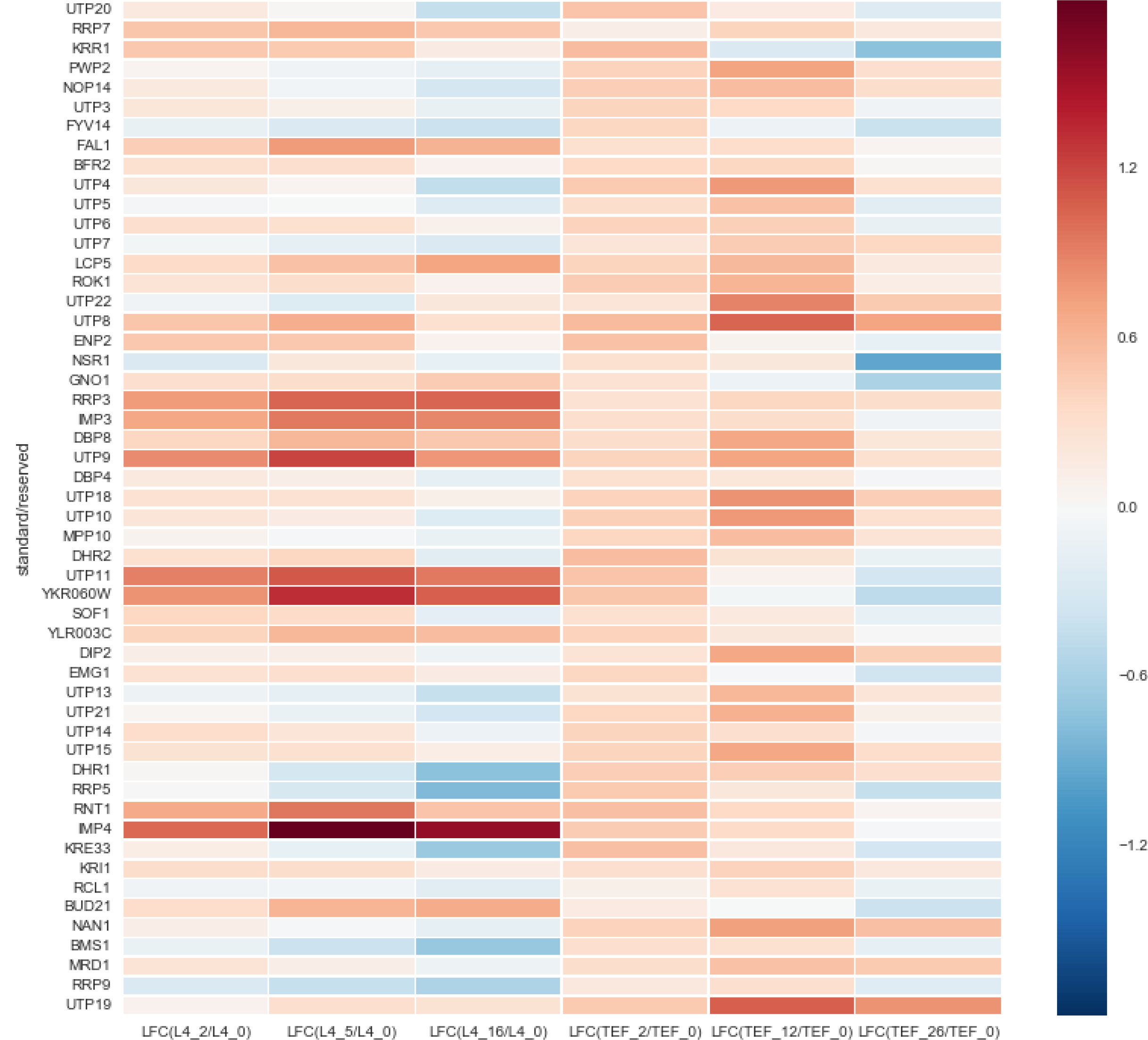
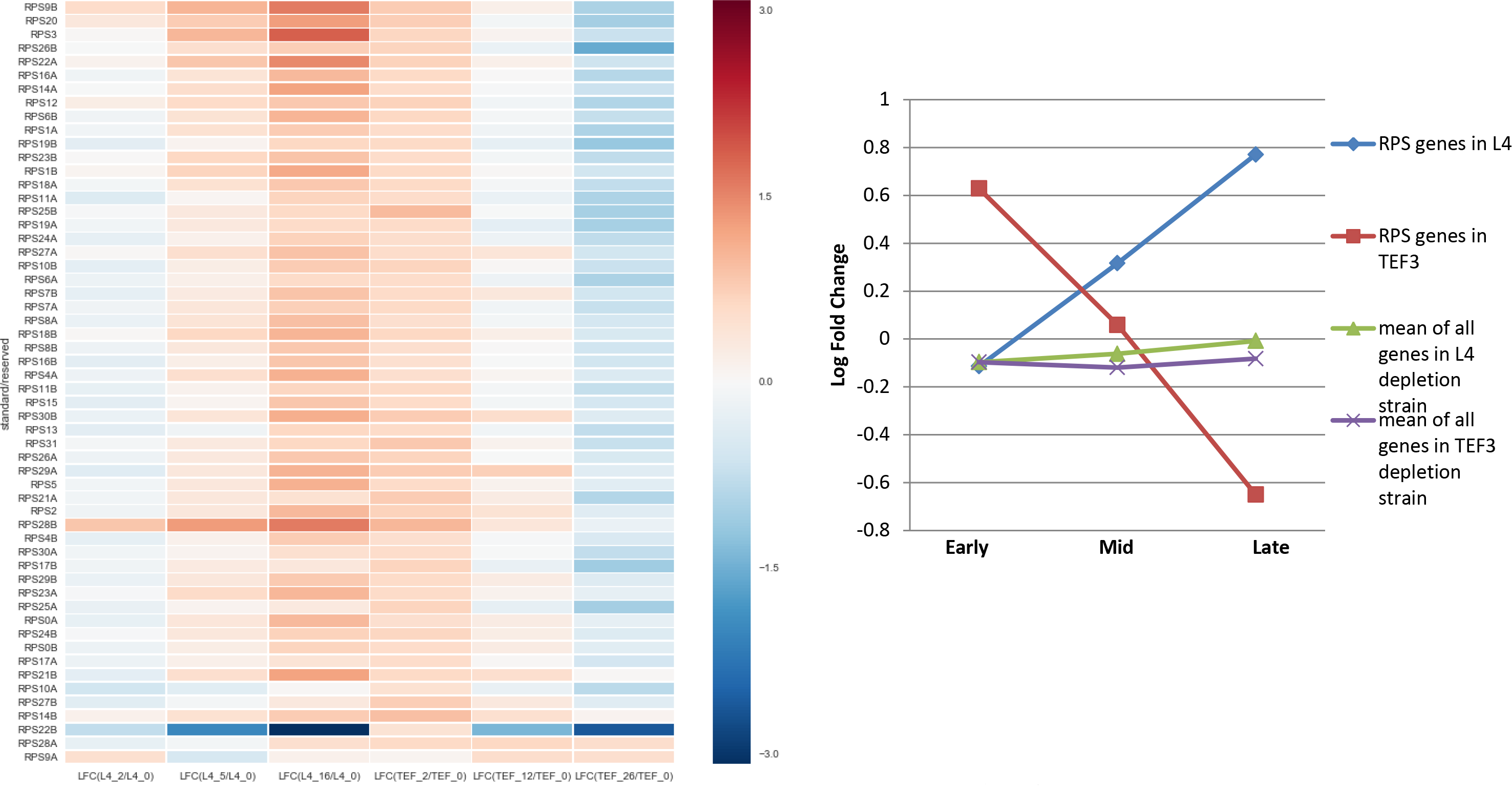
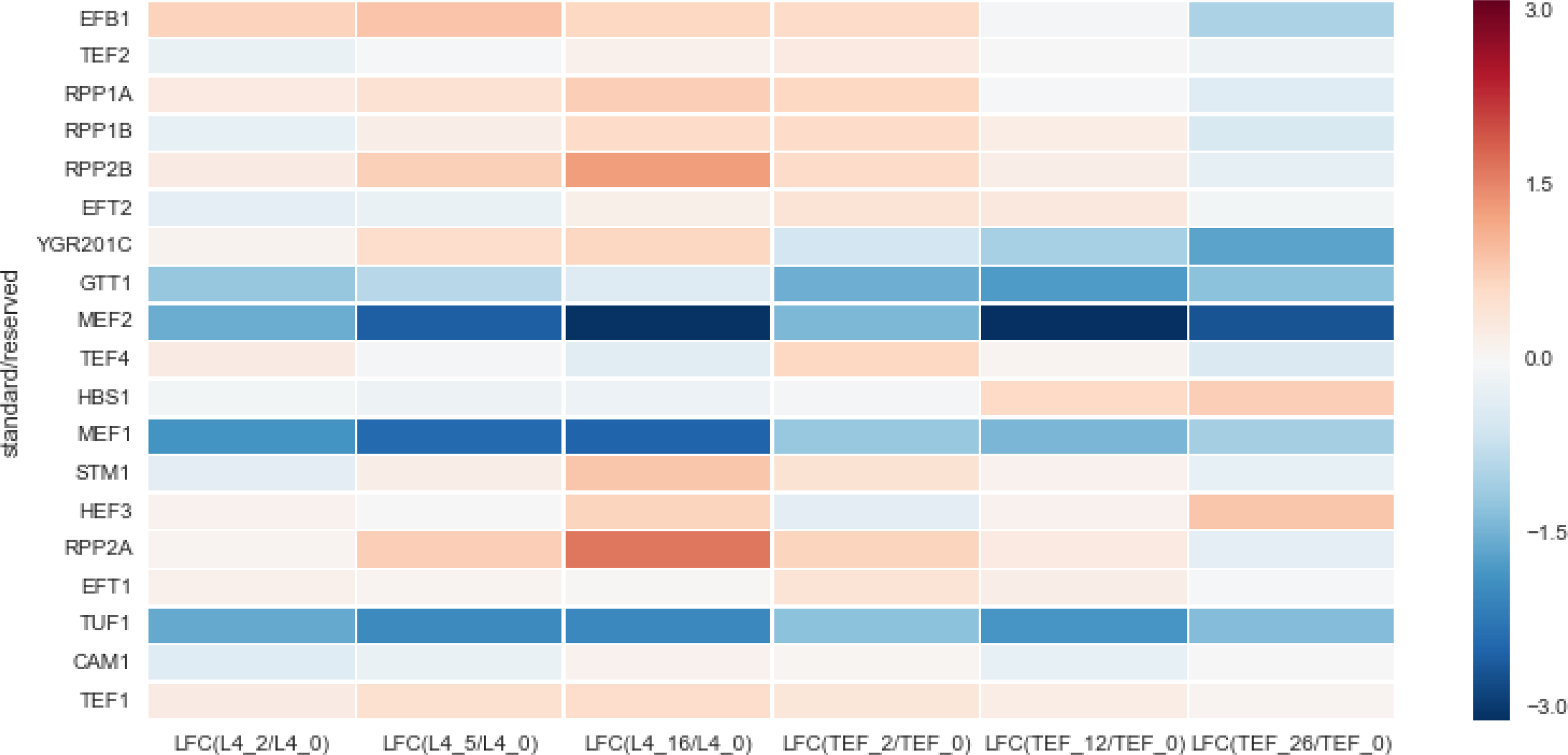
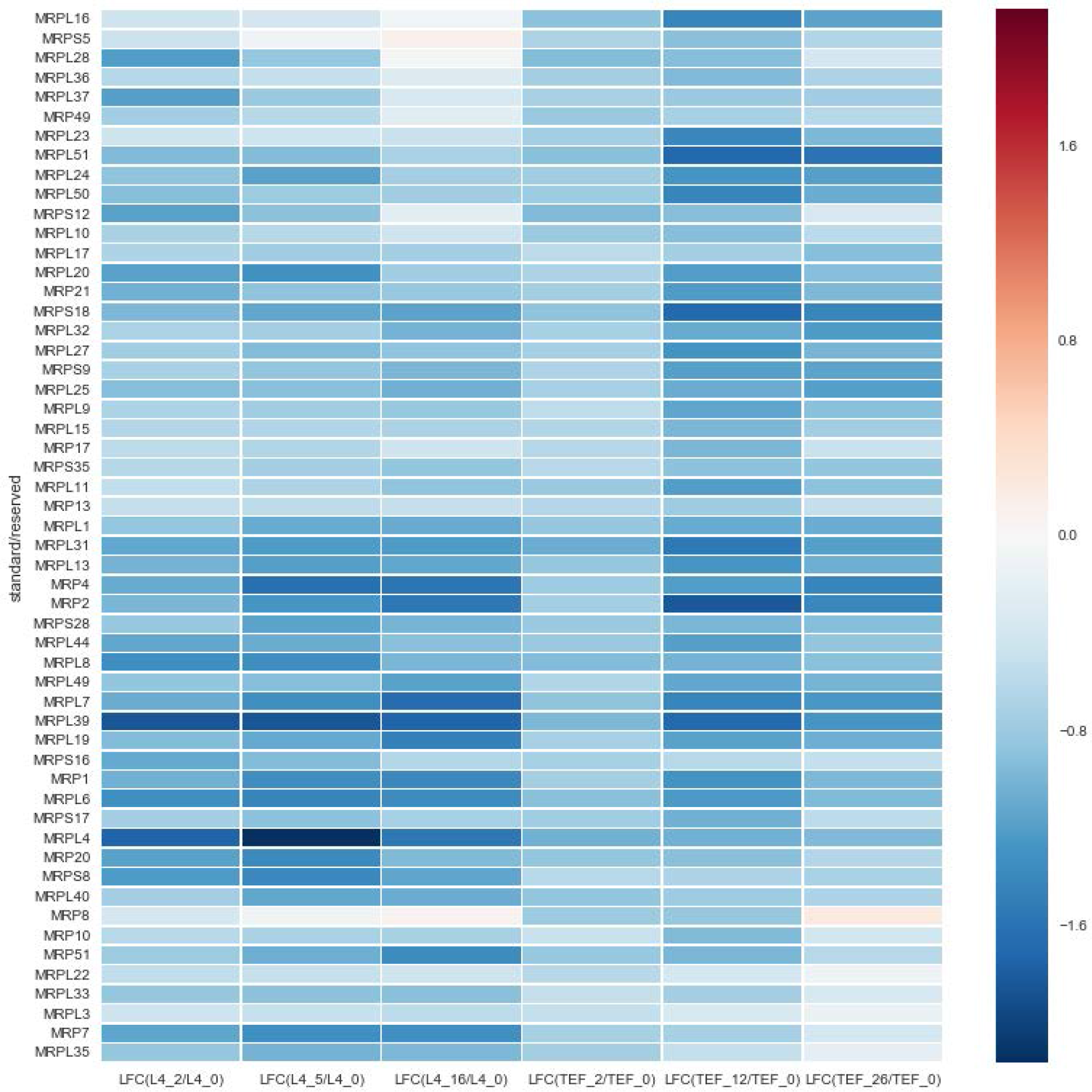
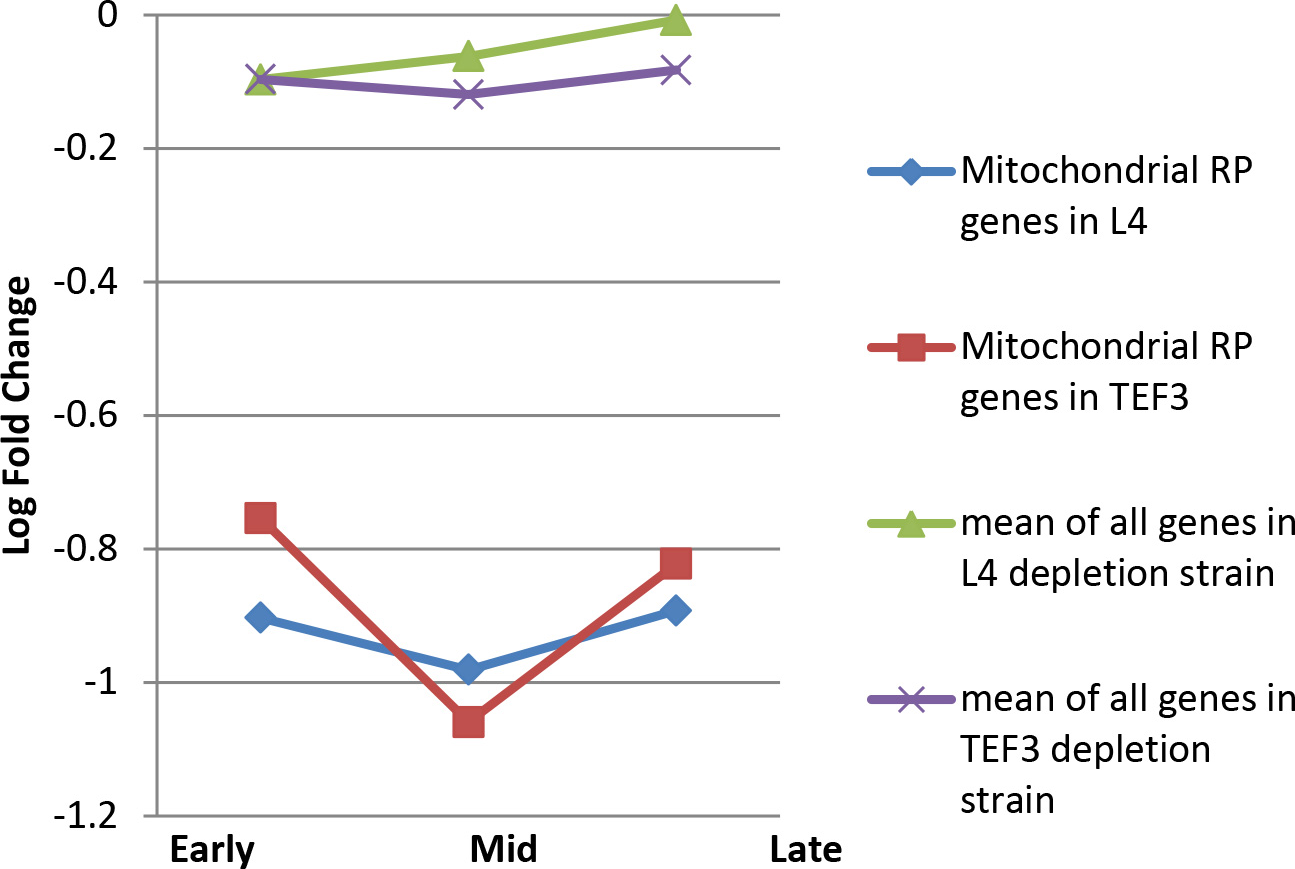
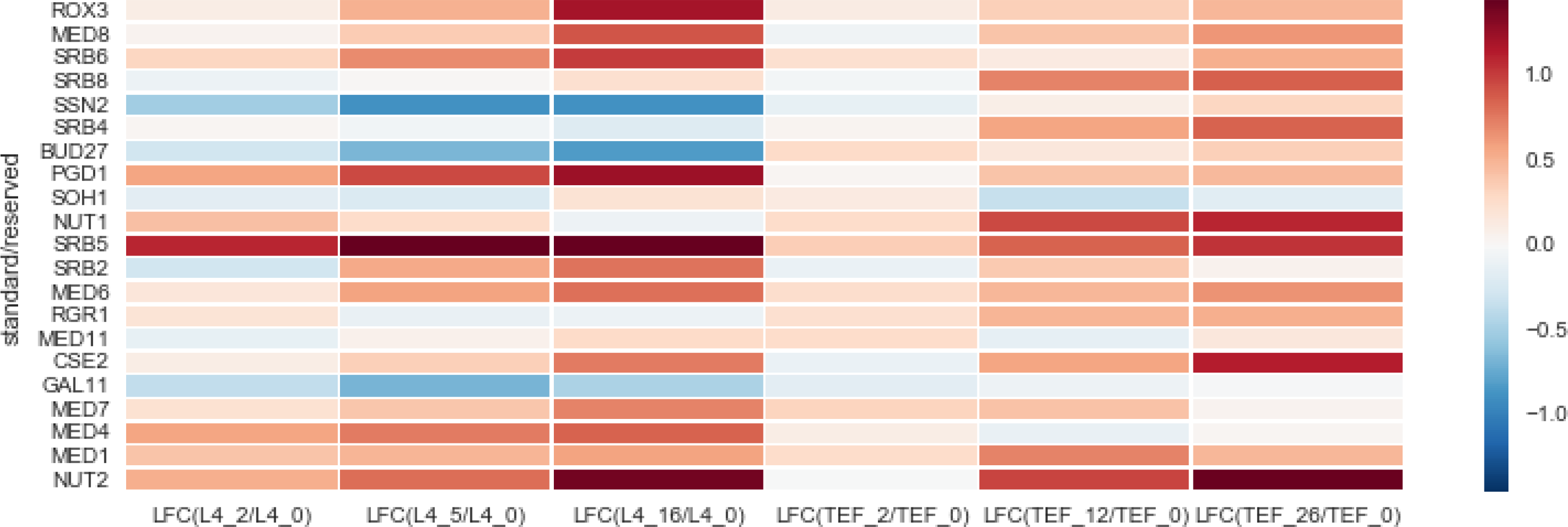
Fold change in expression of genes associated with translation related gene ontology terms. A) Heatmap and scatter plot showing log fold change in expression of large subunit ribosomal protein genes B) Heatmap and scatter plot showing log fold change in expression of small subunit ribosomal protein genes C) Heatmap showing log fold change in the ribosome assembly factor genes D) Heatmap showing log fold change in the GO term translation elongation factor genes E) Heatmap showing log fold change in the mitochondrial ribosomal protein genes. These genes are downregulated at all time points in both of the samples F) Heatmap showing log fold change in the GO term RNA polII transcription cofactor activity genes.

**Fig 9:**
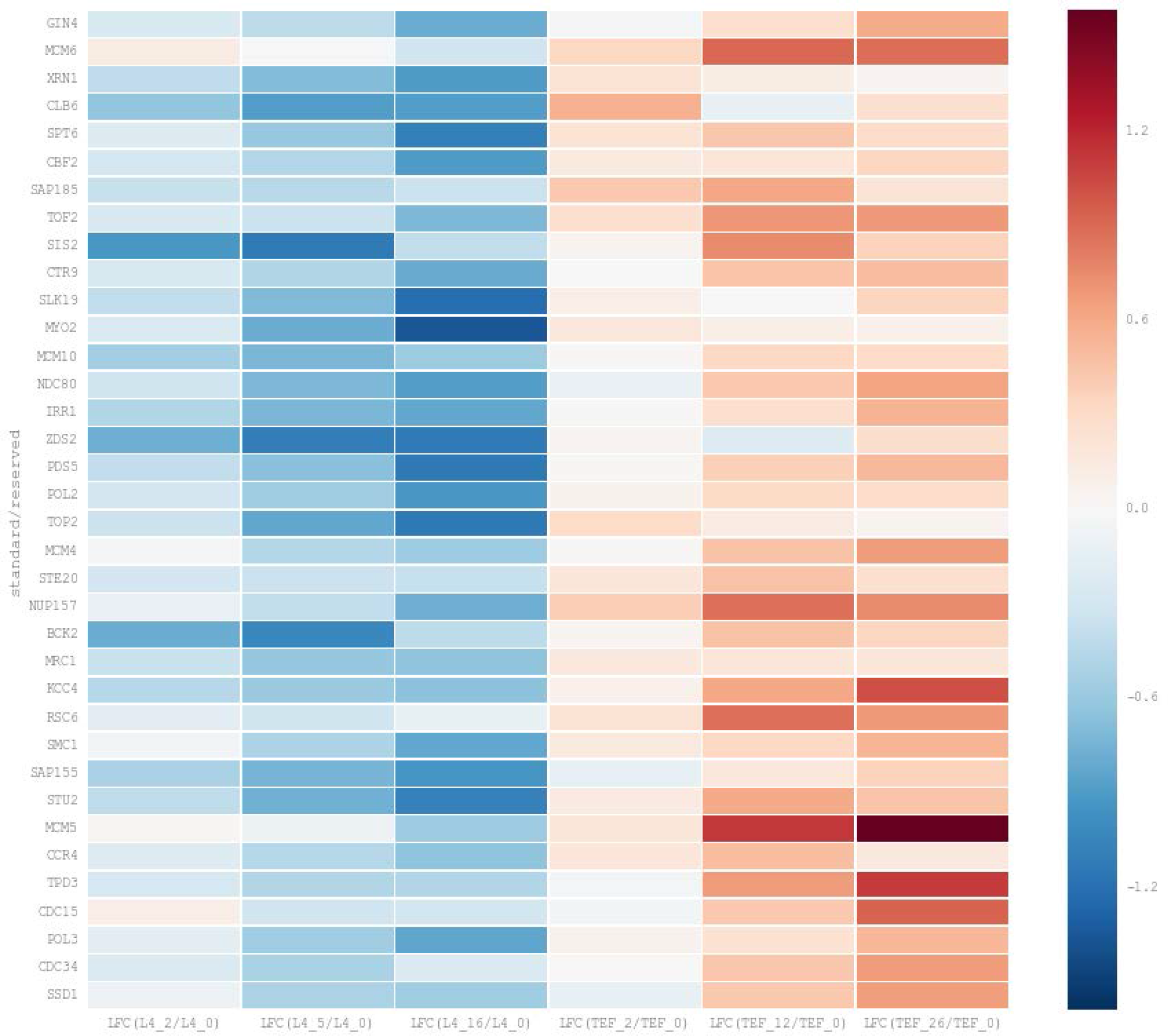
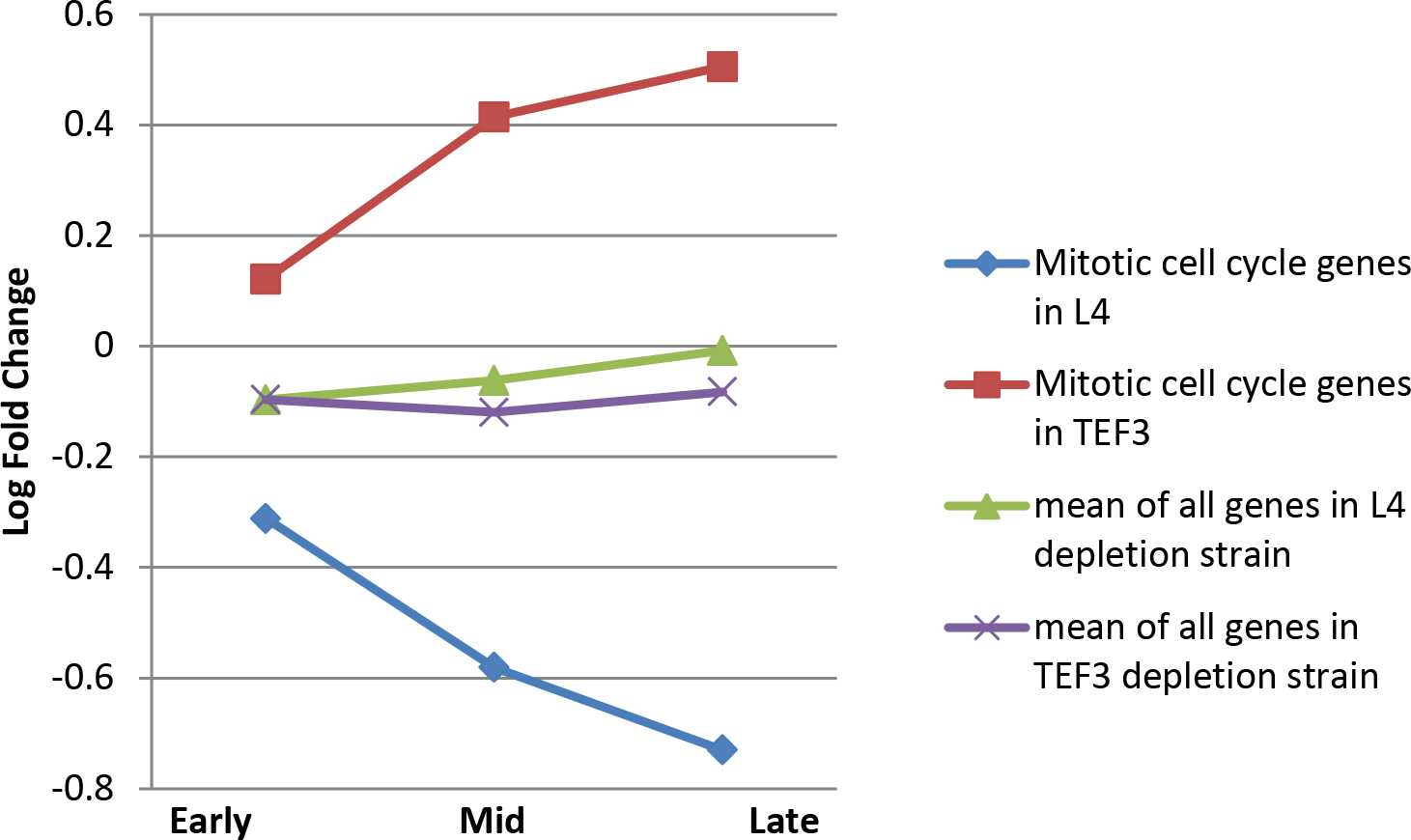
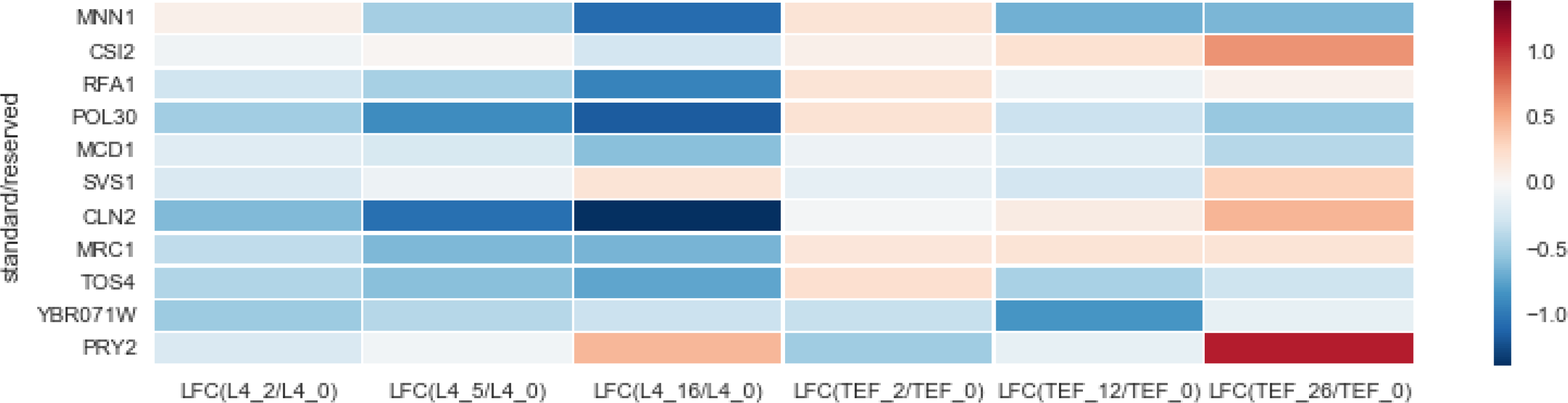
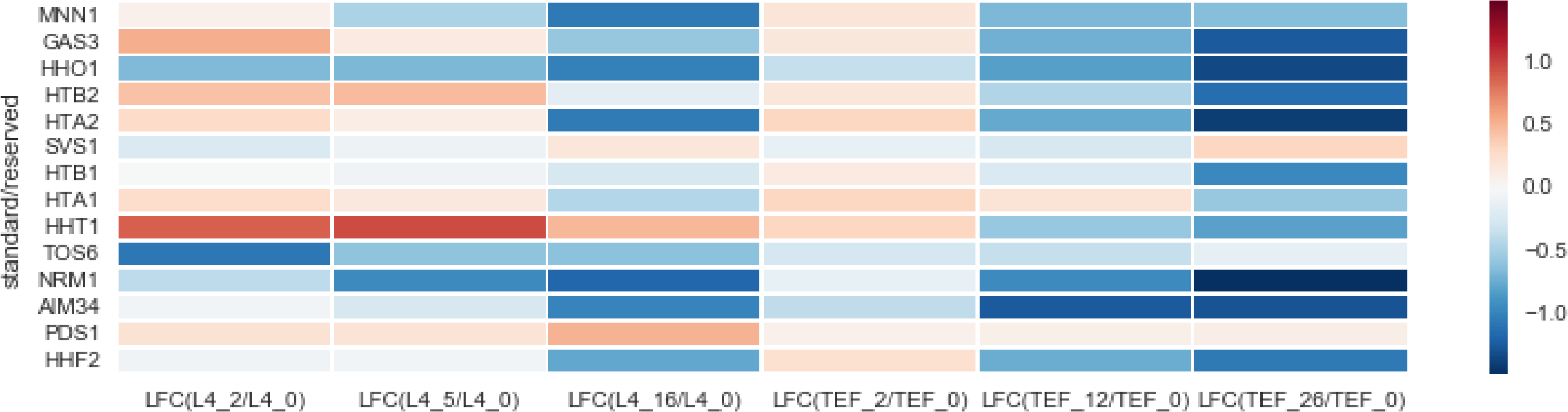
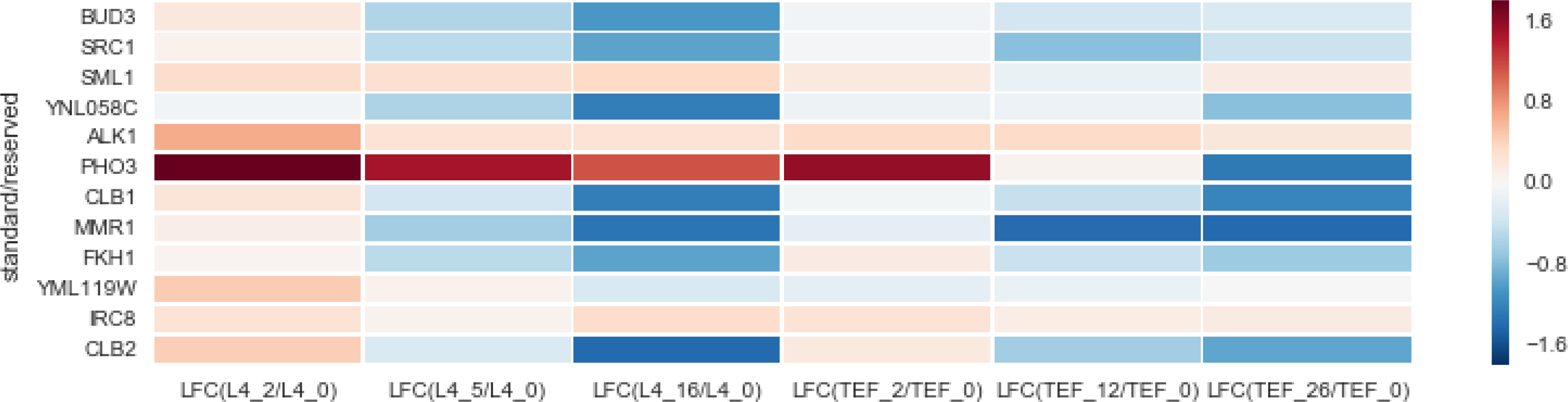
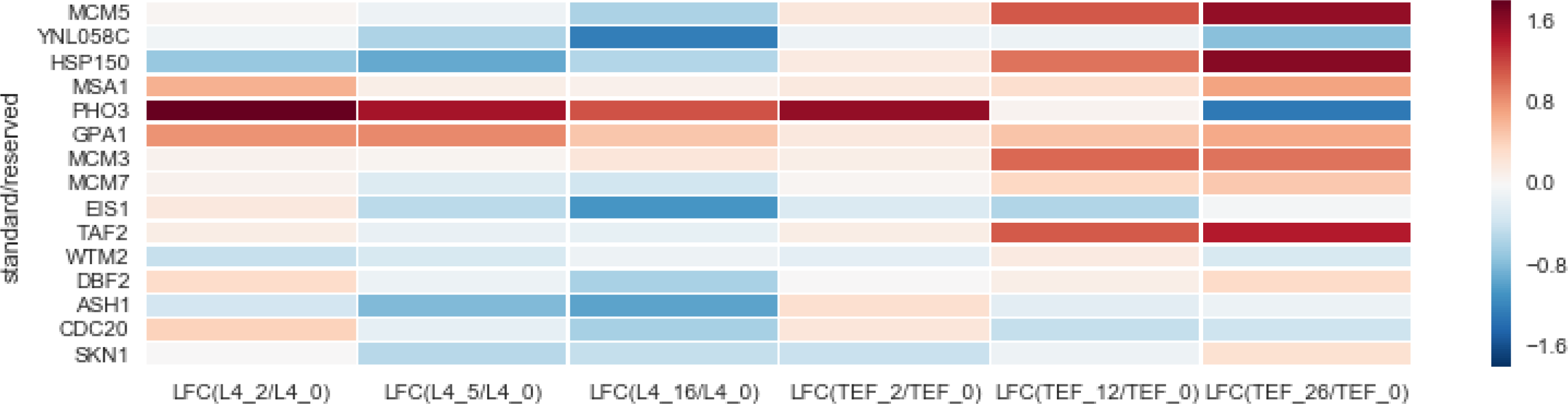
Fold change in expression of genes associated with cell cycle related genes. A) Heatmap showing log fold change in expression of genes associated with the mitotic cell cycle GO term and which are present in the list of top differentiating genes. There is a negative trend in log fold change of expression in L4 depletion strain over time. On the contrary, positive trend in expression is observed in Tef3 samples. Scatter plot shows mean of fold change in expression of this gene set and all the genes identified by RNA-seq in this experiment. B) Heatmap of top genes with G1 specific expression pattern ranked by Cyclebase. There is a negative trend in expression L4 depletion samples and slightly positive trend observed in Tef3 depletion samples C) Heatmap of top genes with S phase specific expression pattern ranked by Cyclebase. Negative trend is observed in both L4 and Tef3 depletion samples D) Heatmap of top genes with G2 specific expression pattern ranked by Cyclebase. Negative trend is observed in both L4 and Tef3 depletion samples E) Heatmap of top genes with M phase specific expression pattern ranked by Cyclebase. There is a negative trend in expression L4 depletion samples and positive trend observed in Tef3 depletion samples

**Fig 10:**
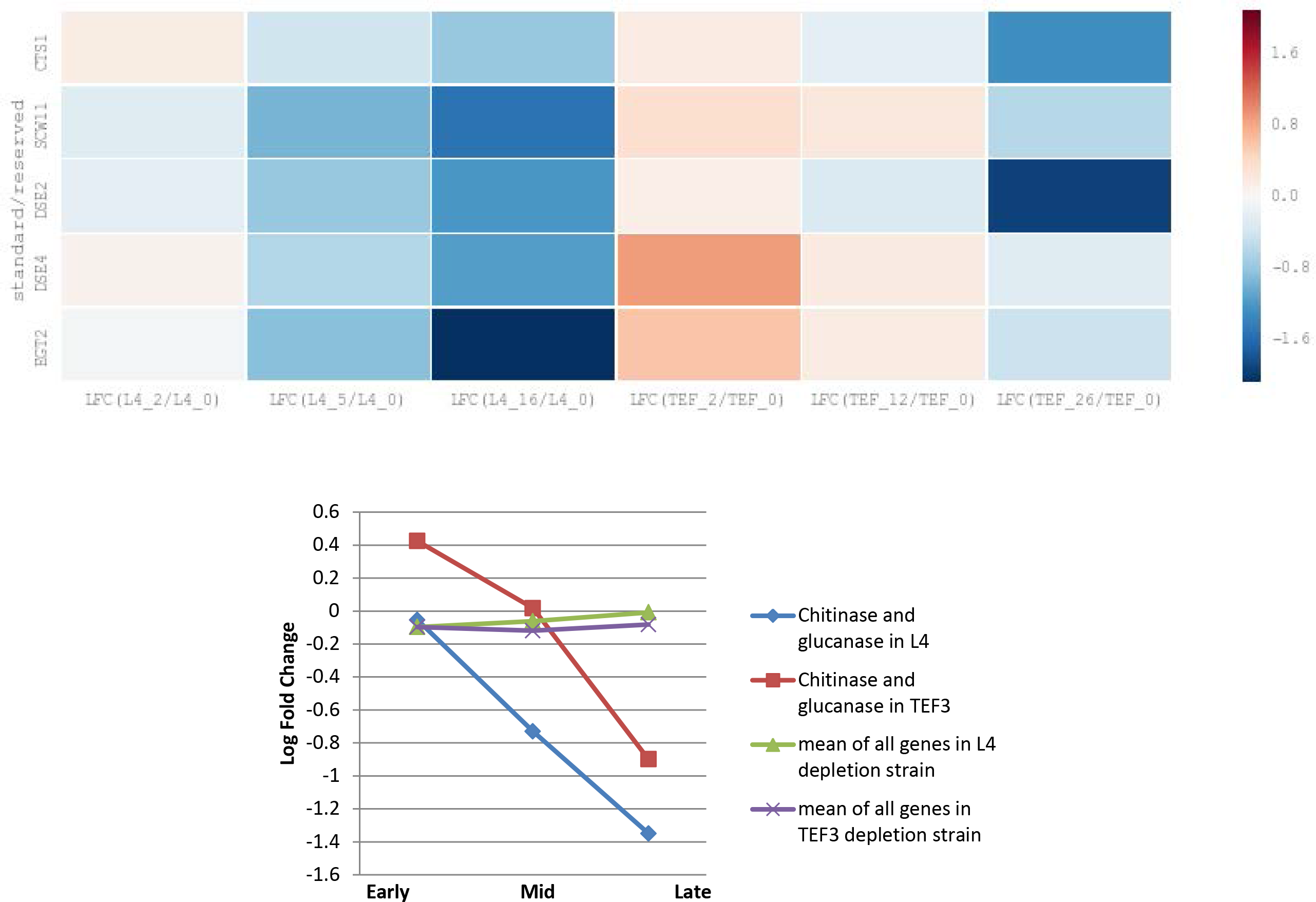
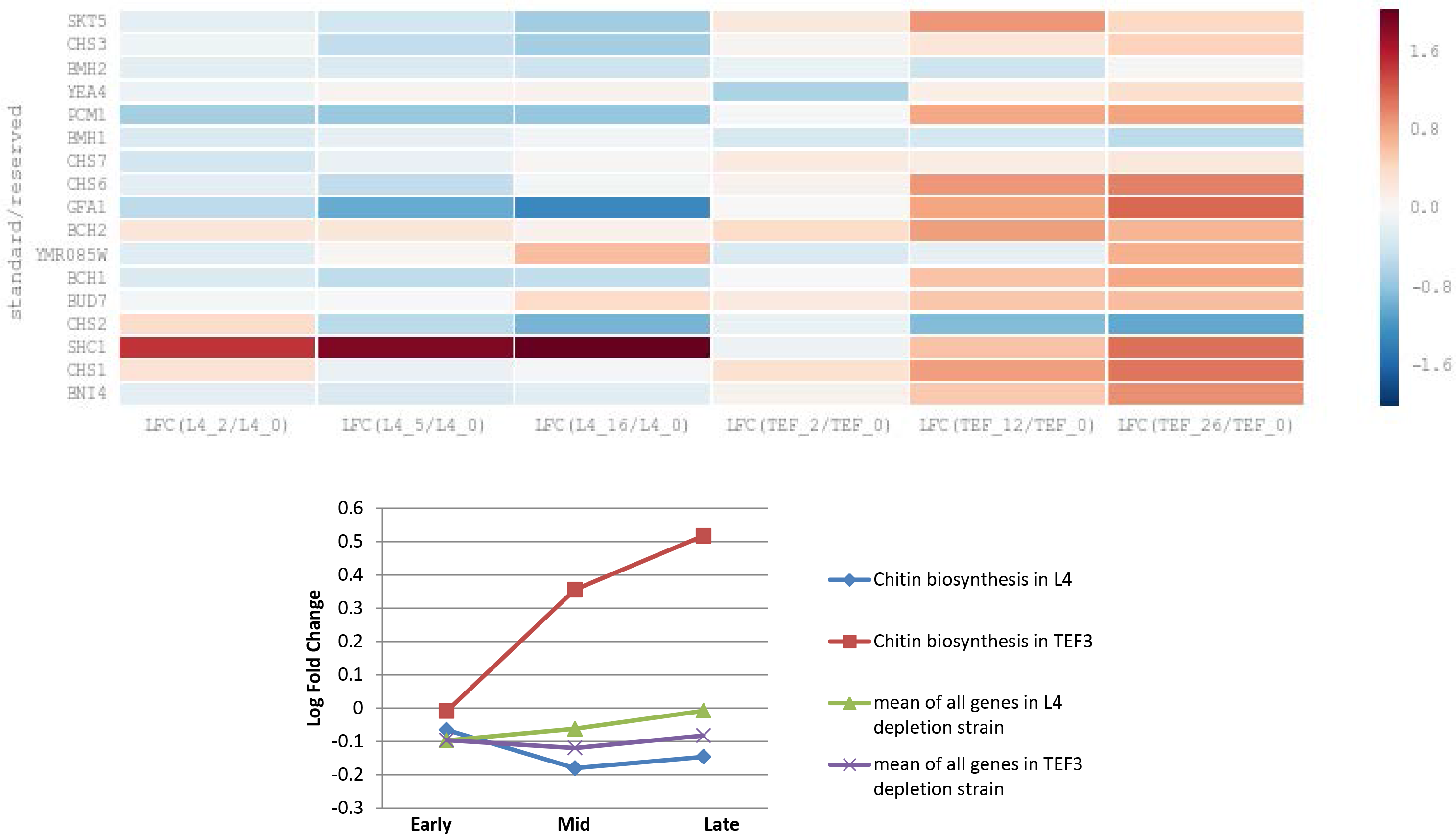
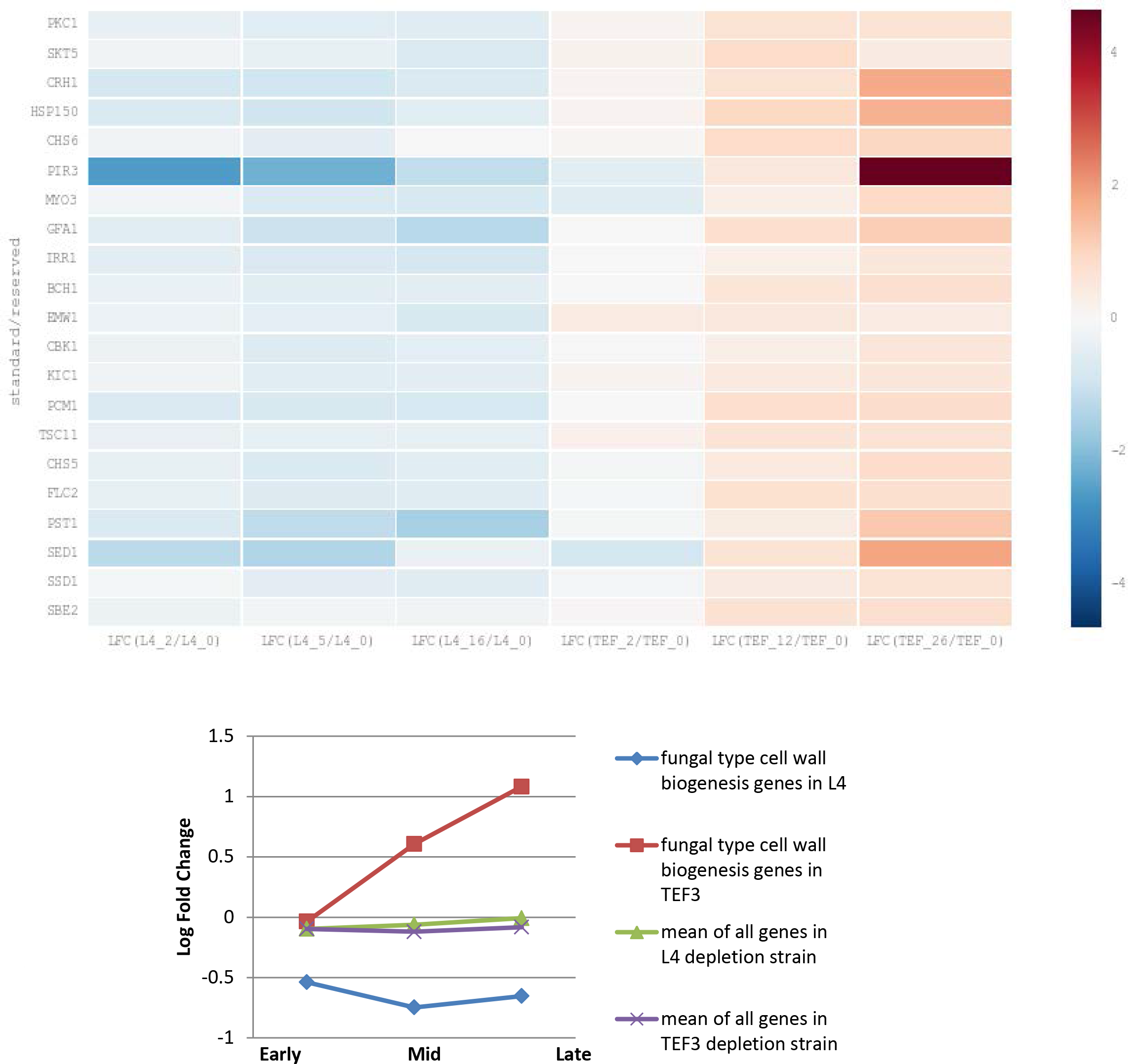
Fold change in expression of genes associated with cell separation and cell wall biosynthesis related gene ontology terms. A) Heatmap showing log fold change in expression of chitinase and glucanase genes associate with cell separation activity. This gene set is down regulated over time in both L4 and Tef3 depletion samples. This gene set is usually expressed in M/G1 boundary by the daughter cells B) Heatmap showing log fold change in expression of chitin biosynthesis genes associated with cell wall repair activity. There is a slightly negative trend in expression L4 depletion samples and positive trend observed in Tef3 depletion samples. This gene set is usually expressed in M/G1 boundary. Expression of these genes indicates either accumulation of cells in this cell cycle stage or need for cell wall repairmen. C) Heatmap showing log fold change in expression of genes associate with the fungal type cell wall biogenesis GO term and which are present in the list of top differentiating genes. There is a slightly negative trend in expression L4 depletion samples and positive trend observed in Tef3 depletion samples.

#### Trend in expression of genes involved in ribosome biogenesis and translation

Since both of ribosomal and translational stress affects translational capacity of the cells, we asked how each of these two stresses affects expression of genes associated with ribosome formation and ribosome function. First, we looked at large and small subunit ribosomal protein genes (RPL and RPS genes, here termed RP genes) (Fig 8A and 8B). As a group, the RP genes for either subunit were first slightly down at 2hr sample in Pgal-L4 strain, but went up at later time points. The opposite trend was seen in Pgal-TEF3 strain (Fig 8A and 8B). Next, we looked at ribosome assembly factor genes. No strong pattern is observed in this gene set (Fig. 8C), showing that RP genes and genes for assembly factors have different expression trends in Pgal-L4 but probably similar trend in Pgal-Tef3.

Next, we identified genes associated with GO term “translation elongation” to determine if there are any compensatory gene expression changes like it is observed for RP genes in Pgal-L4 strain. We did not observe upregulation of genes involved in elongation in Pgal-TEF3 strain (Fig 8D). Note however that (for some mot obvious reason) RPP gene were included in the list returned for a search for elongation factors, but are really ribosome stalk proteins, i.e. structural component of ribosome. These stalk protein genes have a positive trend of expression like other RP genes under ribosomal stress. Also included in the GO term translation elongation are mitochondrial translation elongation factors (TEF1, TEF2, TUF1). These genes are regulated as mitochondrial proteins. The trends for the genes that are actually returned by a search for the term mitochondrial ribosomal protein (MRP) genes are illustrated in (Fig 8E). Interestingly, all the MRP genes are down regulated at all time points in both of the strains. RP genes are transcribed by RNA-pol-II, so we also looked at expression pattern of genes associated with the GO term “RNA-POL-II transcription cofactor activity”. Interestingly, there is a positive trend in expression of these genes under both type of stress (Fig 8F).

#### Trend in expression of genes involved in cell cycle

Multiple studies have shown that ribosomal stress leads to cell cycle arrest in both higher and lower eukaryotes (Oeffinger and Tollervey 2003, Bernstein and Baserga 2004, Gomez-Herreros, Rodriguez-Galan et al. 2013, Thapa, Bommakanti et al. 2013, Polymenis and Aramayo 2015). We observed accumulation of cells in post cytokinesis stage of cell cycle after depleting ribosomal proteins (M. Shamsuzzaman, A. Bommakanti, A. Zapinsky, N. Rahman, C. Pascual, and L. Lindahl, submitted). Searches for (i) GO term enrichment analysis of all genes in the L4 16h sample ranked by log fold change values, and (ii) Tef3 vs L4 cumulative time series based on sum of absolute rank differences, generated a list of genes associated with the GO terms mitotic cell cycle, cell cycle phase and cell division. Hence, we analyzed the pattern of expression of genes associated with the GO term “Mitotic cell cycle” which are present in list of top differentiating genes. We observed a negative trend in cell cycle gene expression in samples under ribosomal stress, but a positive trend in cells under translational stress (Fig 9A). To further analyze the expression of cell cycle genes we examined the expression pattern of genes, which are highly expressed for brief periods specifically in G1 or S or G2 or M phase. All genes, not limited to specific GO terms, have been ranked in the Cyclebase database. The narrower the expression period, the lower the rank number (http://www.cyclebase.org/CyclebaseSearch). We analyzed expression trends during stress of G1, S, G2 and M specific genes with the lowest rank number, i.e. the genes whose expression was most clearly limited to a specific cell cycle phase. Both S and G2 specific gene sets have negative trend of expression with increasing time of ribosomal or translational stress, indicating that an increasing number of cells accumulate beyond S and G2 phase during either type of stress (Fig 9C and 9D). However, the majority of genes specific to M and G1 phase are observed to be down regulated in Pgal-L4 strain, but upregulated during Tef3 depletion beginning at 2hr of stress and continuing through 26hr of translational stress (Fig 9C and 9E).

Cell cycle genes that are not transcribed specifically in any cell cycle phase appear to be down regulated immediately after the repression of L4, as early as 2hr, suggesting that the cells may have exited the cell cycle and have adopted some sort of G0 stage after repression of L4 synthesis and cessation of ribosome formation. However, cells under translational stress actively transcribe M/G1 boundary specific genes and the intensity of expression increases with the time of stress. These cells appear to accumulate in M/G1 boundary rather than exiting the cell cycle (Fig 9B and 9E). This expression pattern supports our observation of cell cycle phenotype we observed under translational stress. Pgal-TEF3 cells under depleting condition increasingly accumulate as mother-daughter cell complex, which is also observed in case of ribosomal protein depletion. However, few daughter cells also go through another round of cell division and form bud on bud phenotype which is absent in cells under ribosomal stress (M. Shamsuzzaman, A. Bommakanti, A. Zapinsky, N. Rahman, C. Pascual, and L. Lindahl, submitted). Daughter cells in translational stress are competent to complete another round of cell cycle whereas cells under ribosomal stress are not.

#### Genes involved in cell separation and cell wall assembly/repair

Cells under either of the stress forms invoked in our work accumulate as mother-daughter complex, which points to a cell separation defect. Hence, we looked at expression pattern of genes involved in post cytokinesis cell separation stage. Four chitinase and glucanase genes are expressed from daughter cell and chew the cell wall from the daughter side (Colman-Lerner, Chin et al. 2001, Weiss 2012). These genes have a negative trend of expression in cells with either of the stresses (Fig 10A). We also looked at the expression pattern of daughter specific genes. These genes also show a negative trend of expression indicating repression of daughter specific genes is a probable cause of cell cycle separation defects (Supplementary fig 5). Interestingly these trends are seen in spite of accumulation of the daughter-specific transcription factor Ace2 in daughter nuclei.

We looked further at expression pattern of genes associated with the enriched GO terms “Chitin biosynthesis” and “Fungal type cell wall biogenesis”, which are present in the list of “top differentiating genes (see above). While there is no trend in change of expression in cells under ribosomal stress, cells under translational stress actively transcribe genes for repairing cell wall (Fig 10B and 10C). These two gene sets are not daughter specific genes and time of expression of these genes coincides with the time of expression of cell separation genes. The constellation of cells in translational stress transcribing chitin biosynthesis genes, but not chitinase and glucanase genes, point to the fact that these cells are incompetent of expressing daughter specific genes only and that is a probable cause of cell separation defect.

#### Other significant GO terms differentially enriched in ribosomal stress and translational stress

Other GO terms that were enriched and have different gene expression pattern in these two strains are ER to Golgi transport, exocytosis, protein transport, protein transport into nucleus and L-amino acid peptidase activity (Fig 7). We looked at the expression pattern of these genes sets, which are present in the list of “top differentiating genes”. All five of these gene sets have positive trend in expression over the time of translational stress, whereas there is negative trend in expression of these genes in cells under ribosomal stress (Fig 11A-E). It appears that the cells under translational stress are compensating for the decreased translational capacity by up regulating the components of the protein delivery system.

**Fig 11:**
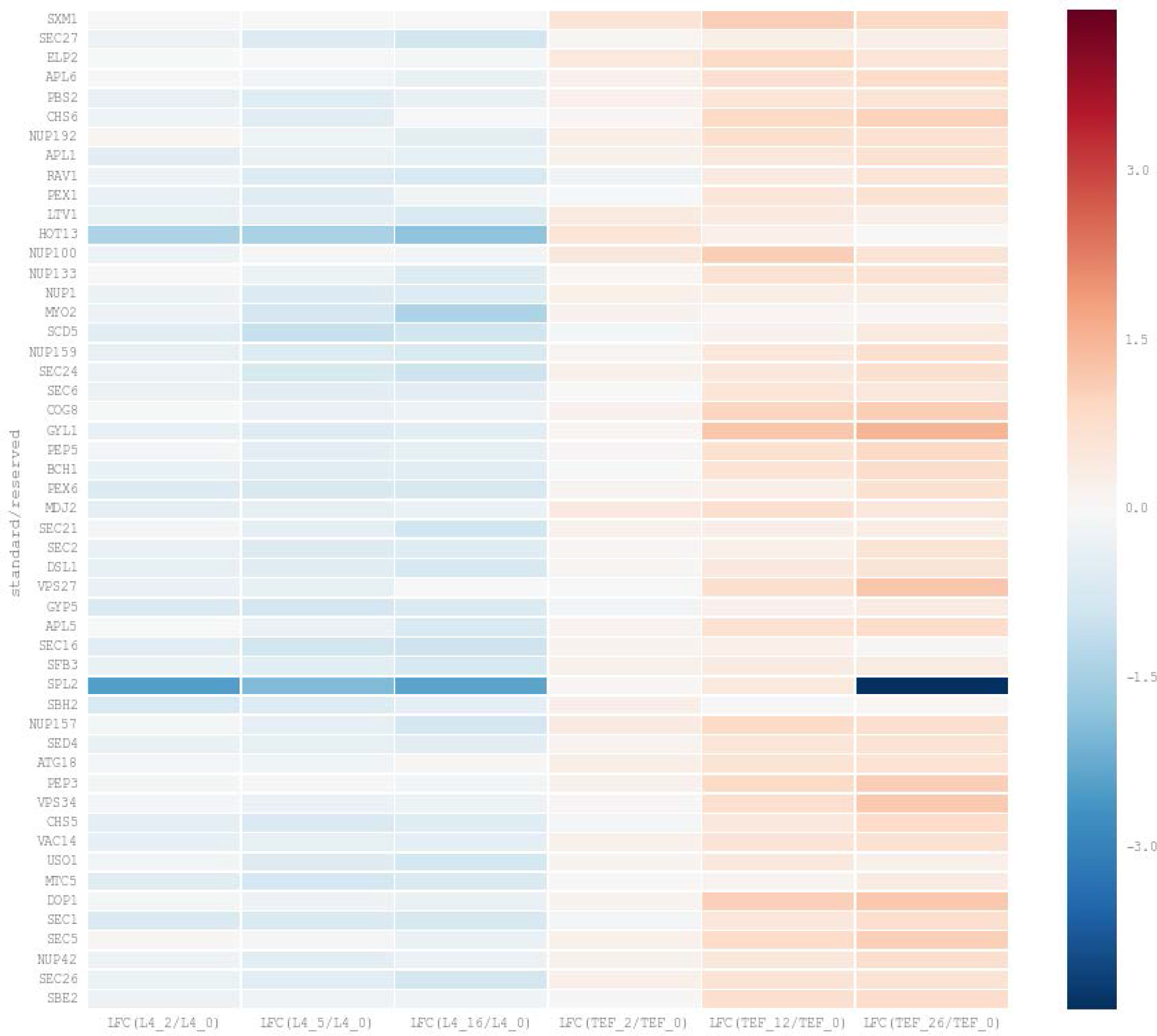
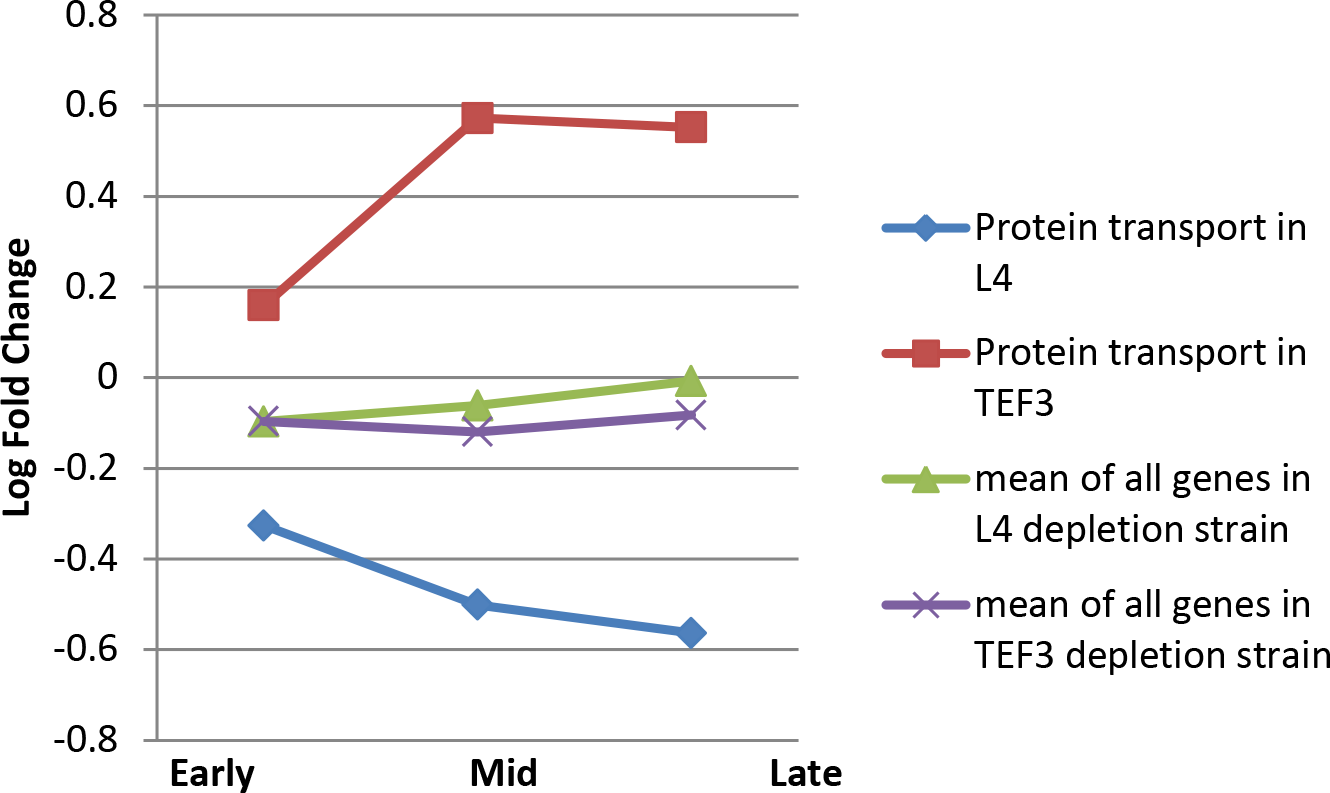
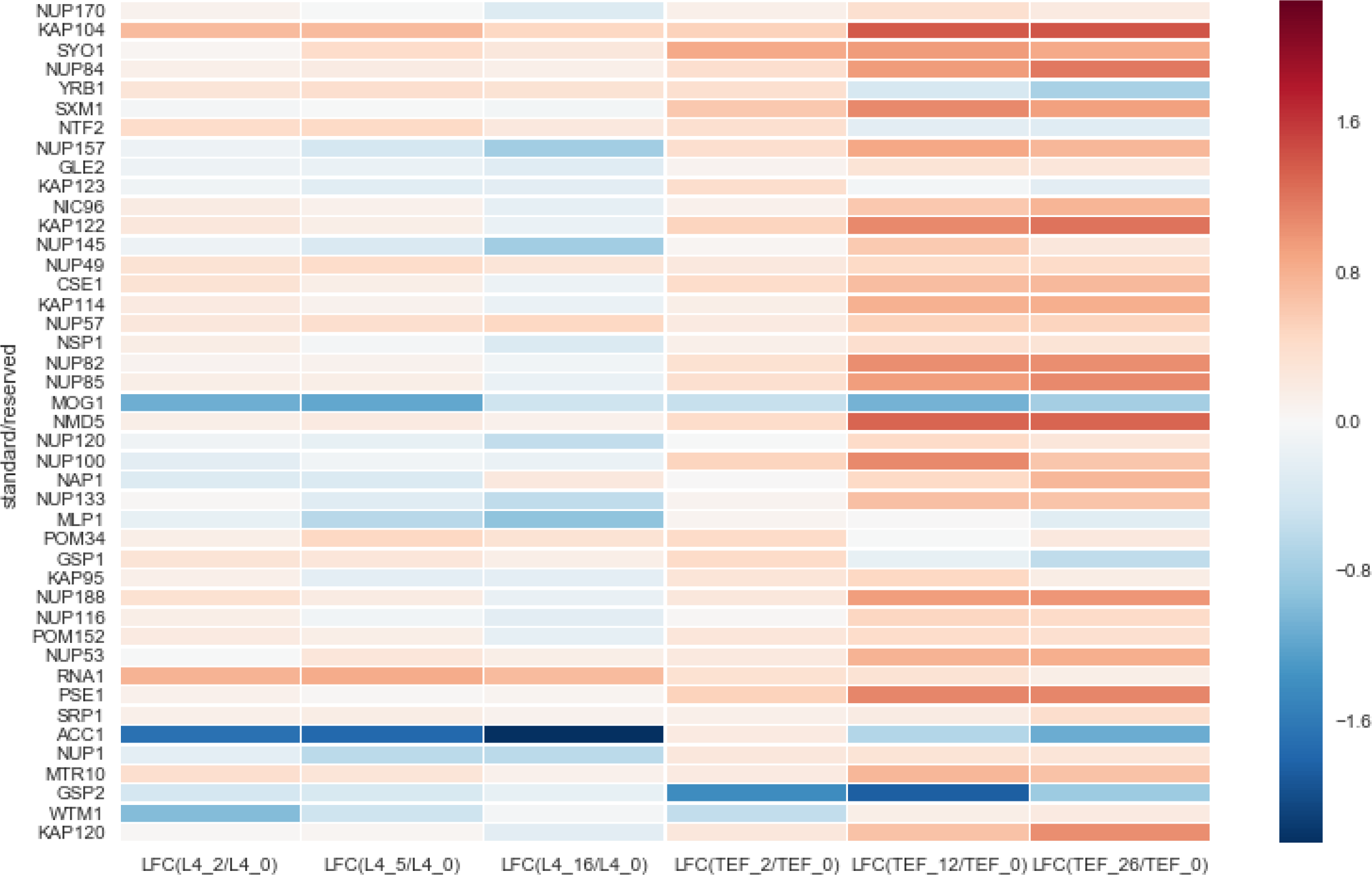
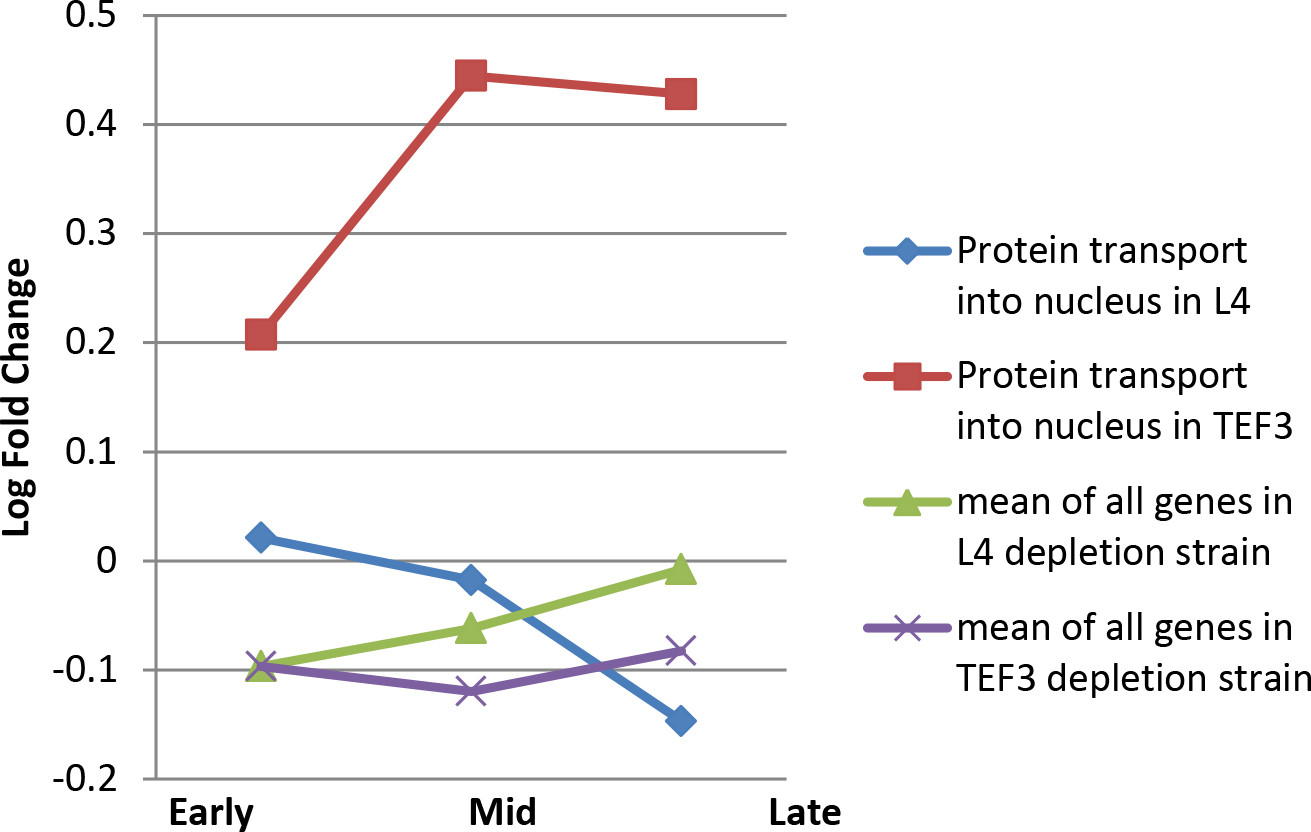
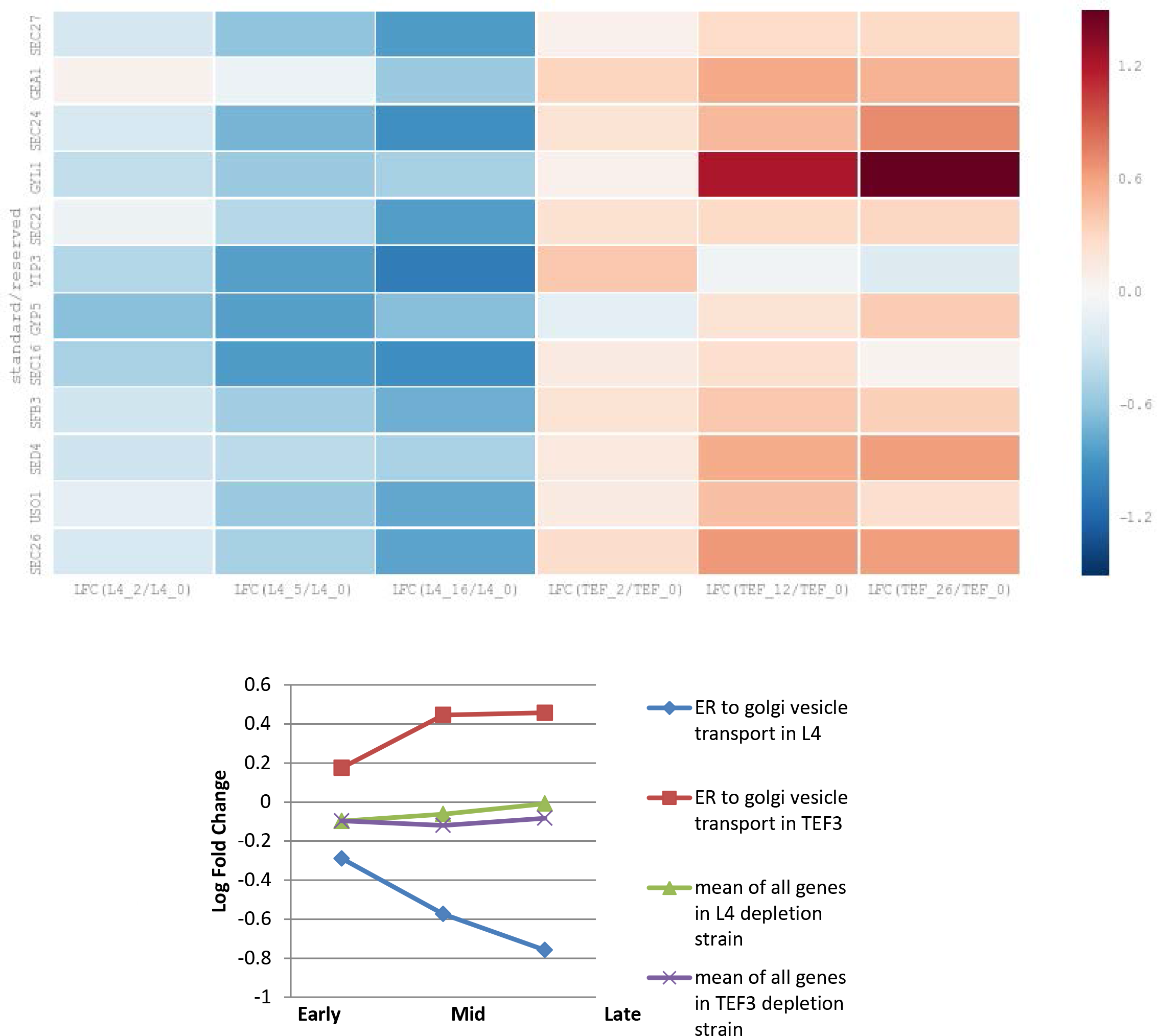
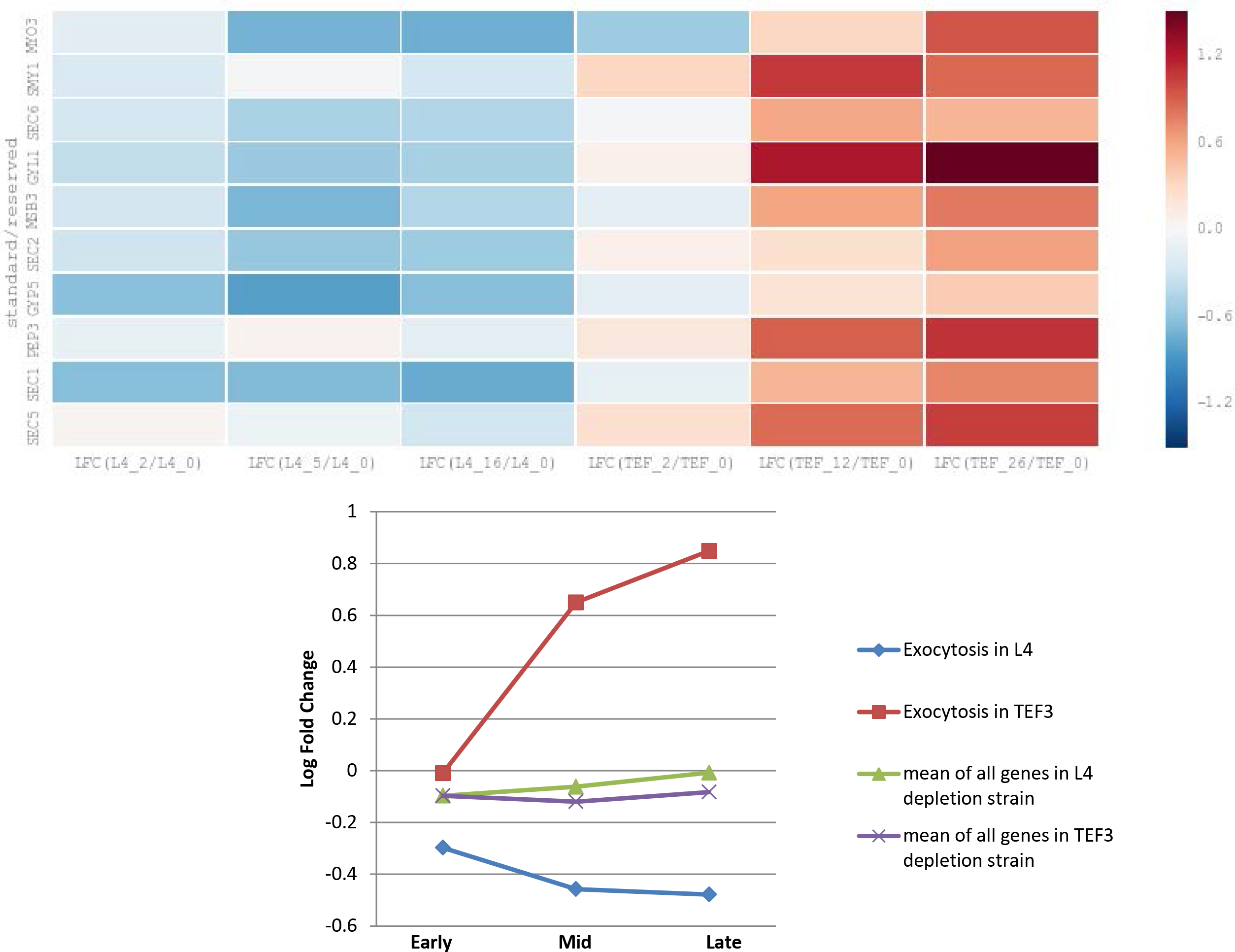
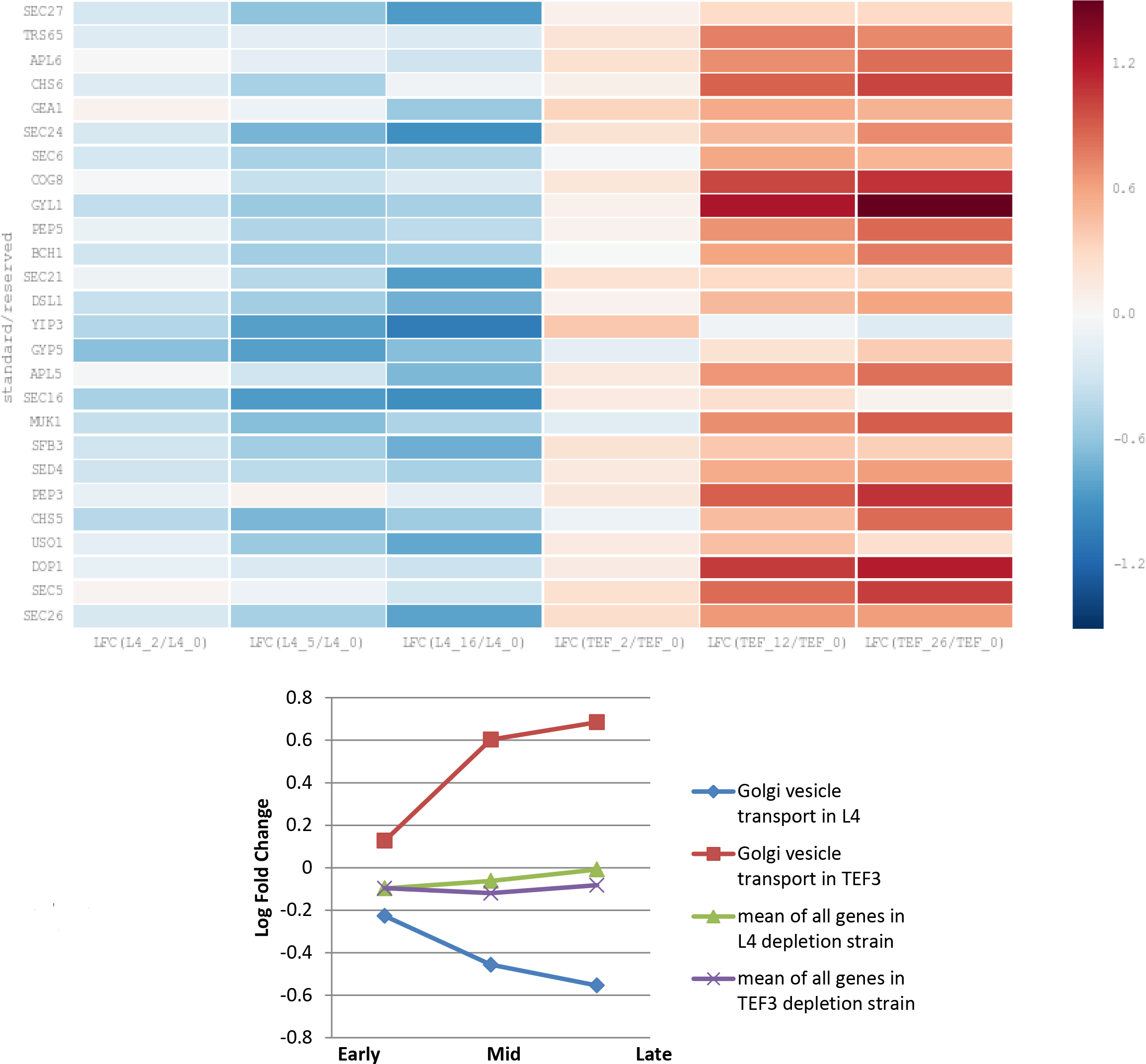
Fold change in expression of genes associated with protein transport and vesicular transport related gene ontology terms. A) Heatmap and scatter plot showing log fold change in expression of DEG genes associated with protein transport gene ontology term and which are present in the list of top differentiating genes. Negative trend is observed in L4 depletion samples but trend is reversed in Tef3 depletion samples. B) Heatmap and scatter plot showing log fold change in expression of genes associate with protein transport into nucleus GO term and which are present in the list of top differentiating genes. There is a slightly negative trend in expression L4 depletion samples and positive trend observed in Tef3 depletion samples. C) Heatmap and scatter plot showing log fold change in expression of DEG genes associated with endoplasmic reticulum to Golgi transport gene ontology term and which are present in the list of top differentiating genes. Negative trend is observed in L4 depletion samples but trend is reversed in Tef3 depletion samples. D) Heatmap and scatter plot showing log fold change in expression of genes associate with exocytosis GO term and which are present in the list of top differentiating genes. There is a slightly negative trend in expression L4 depletion samples and strong positive trend observed in Tef3 depletion samples. E) Heatmap and scatter plot showing log fold change in expression of genes associate with Golgi vesicle transport GO term and which are present in the list of top differentiating genes. There is a negative trend in expression L4 depletion samples and positive trend observed in Tef3 depletion samples.

### GO terms with similar trend in ribosomal stress and translational stress

Multiple gene sets associated with mitochondrion organization, oxidative phosphorylation and mitochondrial transport were enriched in both Pgal-TEF3 and Pgal-L4 strains (Supplementary Fig 1-4). We looked at these gene sets for trend in expression over the time of stress. There is a negative trend in expression in all of these gene sets. Thus, the genes associated with oxidative phosphorylation are more than 3 fold down starting from as early as 2hr of stress (Fig 12A). Cells sense these stress very early and shuts down powerhouse of cell and stop producing energy.

**Fig 12:**
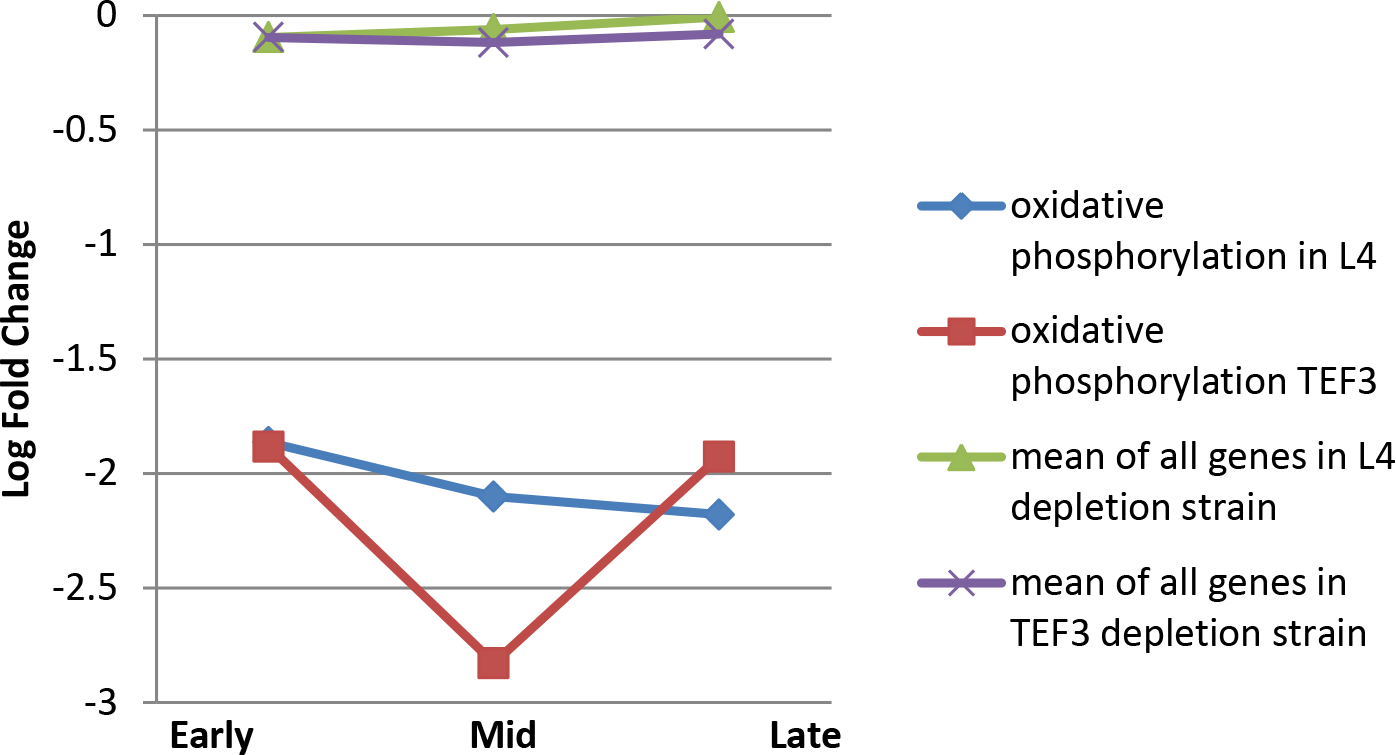
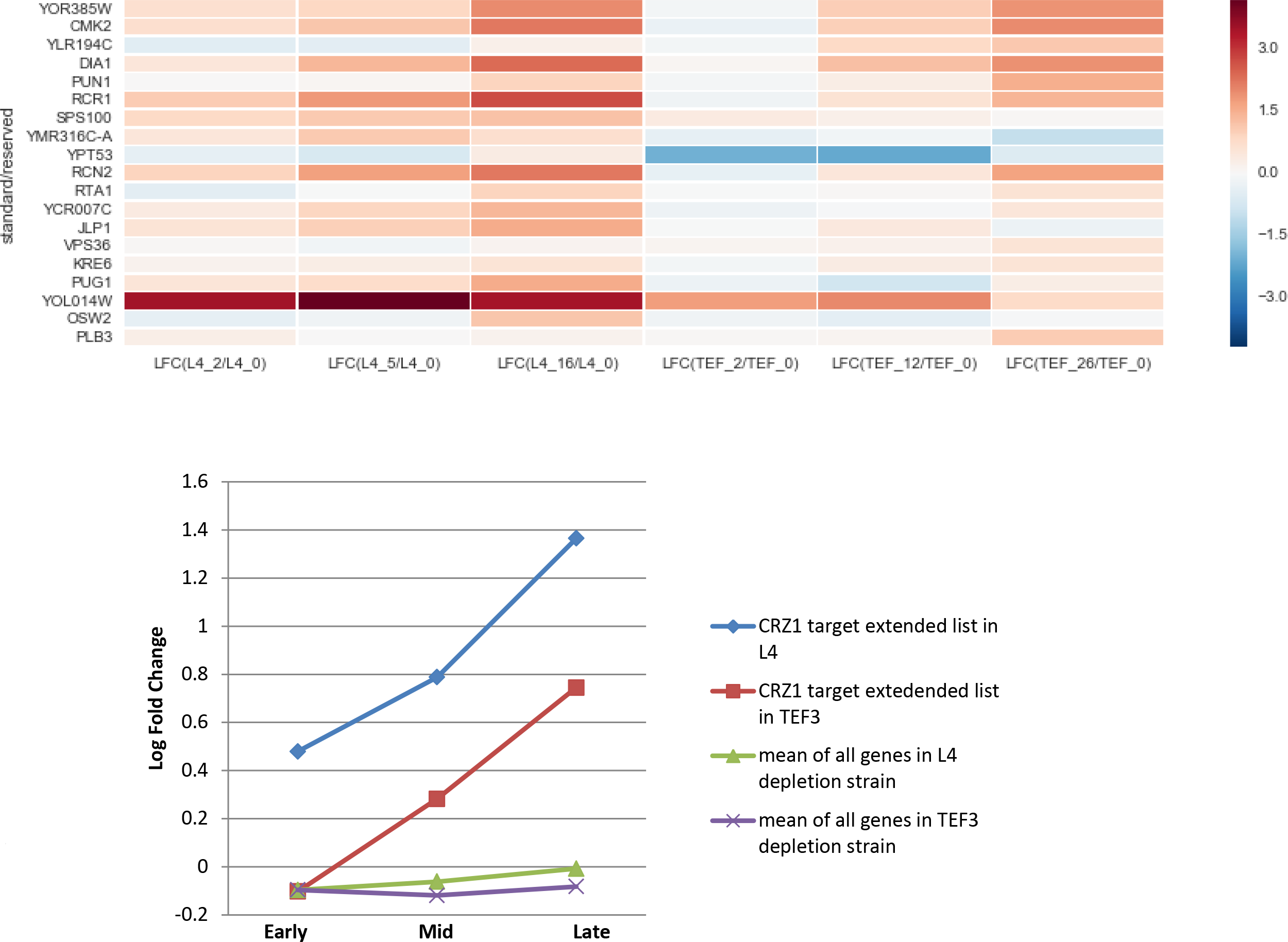
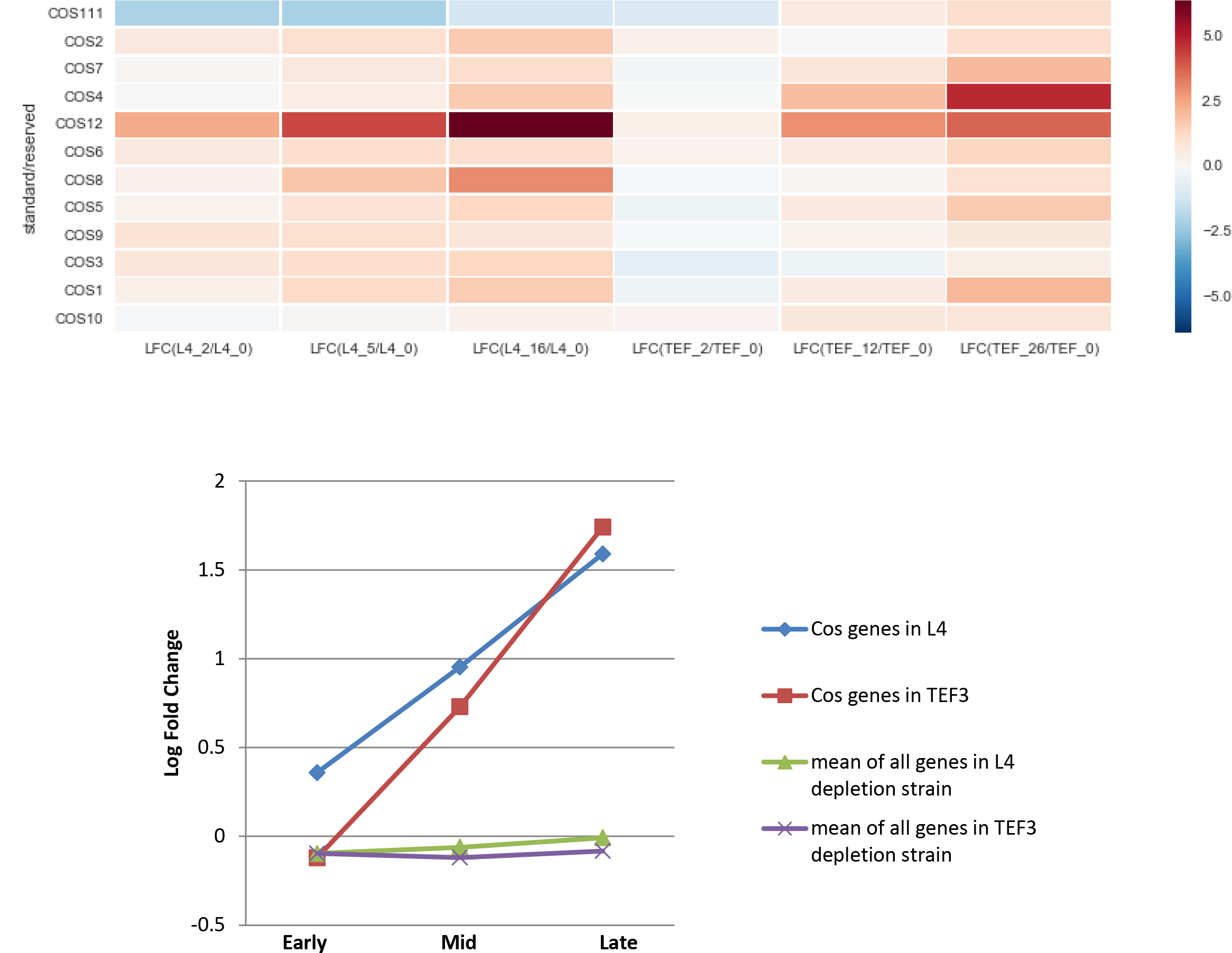
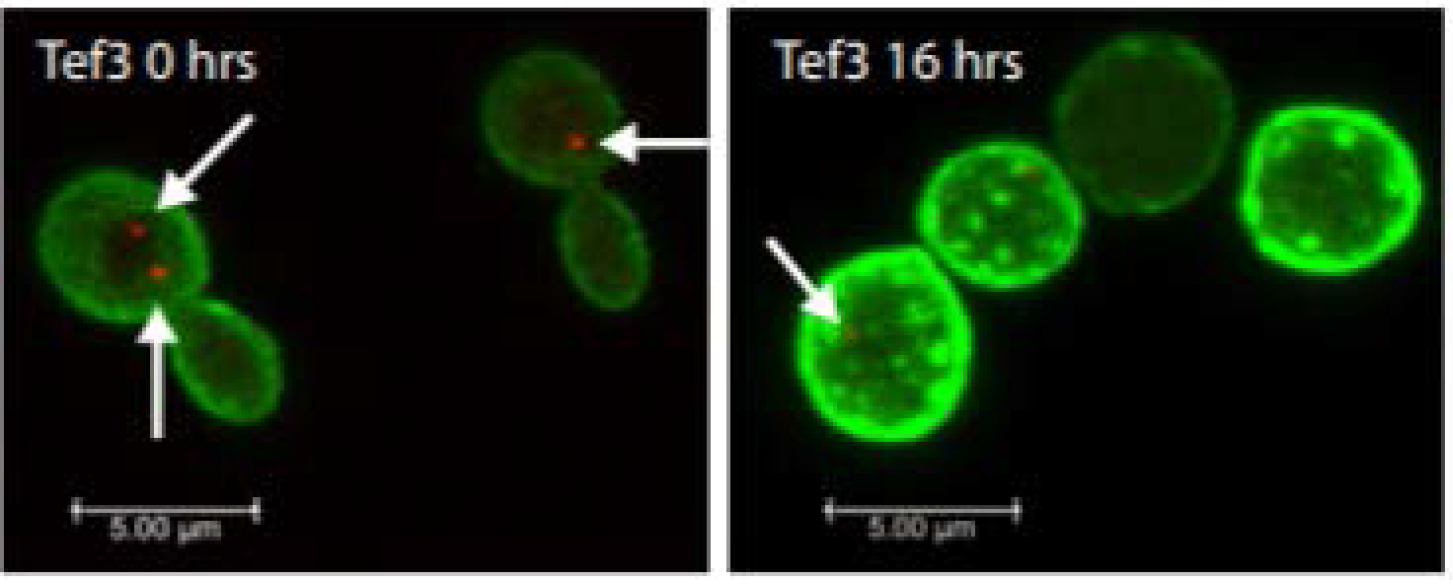
Gene sets with similar trend in fold change of expression in both L4 and Tef3 depletion samples. A) Scatter plot showing mean of log fold change in expression of the gene set associated with oxidative phosphorylation GO term and of all the genes identified by RNA-seq in this experiment. This gene set is downregulated early in both depletion samples and continued to be down regulated over time of depletion. B) Heatmap showing log fold change in expression of genes regulated by CRZ1 transcription factor under multiple stress condition. Very similar strong positive trend is observed in both L4 and Tef3 depletion samples. C) Heatmap showing log fold change in expression of COS genes involved in sorting of membrane proteins into multi-vesicular bodies. Strong positive trend is observed in both L4 and Tef3 depletion samples. D) Fluorescence microscopy images showing engulfing and sorting of GFP tagged Ras2 membrane protein Tef3 depletion sample.

A very common stress signaling pathway is Calcineurin-Crz1 mediated upregulation of stress responsive genes. These genes are expressed in heat shock, oxidative stress, osmotic stress (Yoshimoto, Saltsman et al. 2002). Genes include protein needed to reinforce cell wall synthesis, lipid and sterol metabolism, Ion transport and homeostasis, vesicle transport (Yoshimoto, Saltsman et al. 2002). Genes reported to be most influenced by Calcineurin-Crz1 pathway (Yoshimoto, Saltsman et al. 2002). Had a positive trend both during translational stress and ribosomal stress (Fig 12B). It appears that the cells in translational stress lags behind the ribosomal stress in terms of eliciting the Crz1 mediated stress response. Whereas some of the stress responses like shutting down mitochondria were very immediate, the Crz1 stress response is very gradual, possibly reflecting gradual increase in Ca^++^ concentration in response to stress rather than instant activation or inactivation of any kinase.

Finally, we looked at expression pattern of a set of the 12 Cos genes, which are induced during starvation and are involved in sensing high level of cellular NAD+ concentration by NAD+/Sir2 dependent mechanism (Macdonald 2015). Cos proteins are important in sorting non-ubiquitinated proteins into multivesicular bodies (MBV) for degradation. Most of the membrane proteins are sorted into MBV and are taken to vacuole and lysozyme for degradation in this pathway. Cells under both ribosomal stress and translational stress significantly upregulate COS genes expression (Fig 12C).

A characteristic phenotype of overexpression of COS genes is engulfing membrane proteins and formation of visible large endosomes resulting from engulfing the membrane proteins (Macdonald 2015). Accordingly, we examined the vesicles in GFP-Ras2 (a marker of plasma membranes) tagged cells by projecting the images in the Z-stacks from confocal microscopy into a single plane to produce a view of the vesicles in the entire cell. After the shift to glucose medium Pgal-eL43 developed large complex Ras2-bounded vesicles (Supplementary Fig 6), while vesicles remained small, but increased in number in Pgal-Tef3 (Fig 12D). It is probable that cells, under oxidative stress condition, membrane proteins are oxidized and needs to be replaced frequently.

### Transcription factor (TF) over-representation analysis of up and down regulated DEGs

To understand the transcription factors activity under these two stresses, we analyzed transcription factor over-representation analysis using Yeastract database (Monteiro, Mendes et al. 2008). We did TF over-representation analysis of up and down regulated DEGs separately on both L4 and Tef3 early and late depletion samples. The top 5 significantly enriched TFs during early and late ribosomal and translational stress are shown in Table 2. Among the transcription factors activated, Gat4 is the most significantly enriched in both stresses in three out of the four samples (Table 2). Gat4 is a stress sensitive transcription factor involved in outer spore wall assembly in *Yeast* (Lin, Kim et al. 2013). Enrichment of Gat4 activated genes increases from 8.7% to 22% from 2h to 16h of ribosomal stress, indicating gradual buildup of stress condition and consequent increase in Gat4 target gene expression. 5.5% of Gat4 target genes were also activated in Pgal-Tef3, much less than percentage of target genes activated in Pgal-L4.

**Table 2:** Over representation analysis of TFs.

|  | TFs as<br>activator | % of target genes in<br>this list to number of<br>genes in this list | % of target genes in this list<br>to number of target genes in<br>the genome | P-value |
| --- | --- | --- | --- | --- |
| **L4 2h** | Gat4 | 7.36% | 8.72% | 0.000111422 |
|  | Upc2 | 8.91% | 7.23% | 0.000475682 |
|  | Lys14 | 0.78% | 28.57% | 0.001469326 |
|  | Gzf3 | 6.98% | 6.23% | 0.008335715 |
|  | Eds1 | 3.88% | 7.35% | 0.010047086 |
| **L4 16h** | Gat4 | 6.79% | 22.02% | 2.22064E-08 |
|  | Gis1 | 9.90% | 17.86% | 2.20301E-07 |
|  | Upc2 | 7.07% | 15.72% | 0.000317092 |
|  | Pdc2 | 0.99% | 26.92% | 0.002764135 |
|  | msa2 | 0.28% | 50.00% | 0.003592735 |
| **Tef3 2h** | Gat4 | 17.39% | 5.50% | 1.20824E-07 |
|  | Mig2 | 17.39% | 3.91% | 5.91506E-06 |
|  | Pdc2 | 2.90% | 7.69% | 0.001920704 |
|  | Mot3 | 17.39% | 2.04% | 0.003750499 |
|  | Gis1 | 13.04% | 2.30% | 0.004144943 |
| **Tef3 26h** | Rlm1 | 17.87% | 9.40% | 5.32365E-07 |
|  | Wtm2 | 5.07% | 11.24% | 0.000522297 |
|  | Crz1 | 13.07% | 7.67% | 0.002478779 |
|  | Rgm1 | 8.80% | 8.33% | 0.002952907 |
|  | Gat4 | 5.33% | 9.17% | 0.005103723 |

Table 2
|  | TFs as<br>repressor | % of target genes in<br>this list to number of<br>genes in this list | % of target genes in this<br>list to number of target<br>genes in the genome | P-value |
| --- | --- | --- | --- | --- |
|  | Sut1 | 19.40% | 18.91% | 0 |
|  | Mig2 | 12.50% | 21.82% | 0 |
|  | Bas1 | 52.05% | 9.29% | 5.669E-07 |
|  | Rsc30 | 3.92% | 20.59% | 4.848E-06 |
|  | Mig1 | 11.01% | 13.02% | 7.634E-06 |
|  | Sut1 | 16.16% | 18.36% | 3.2E-14 |
|  | Mig2 | 10.56% | 21.50% | 6.66E-13 |
|  | Mot3 | 13.12% | 13.92% | 5.24E-06 |
|  | YPR015C | 4.16% | 18.84% | 5.055E-05 |
|  | Mig1 | 10.08% | 13.91% | 6.392E-05 |
|  | Bas1 | 66.84% | 8.59% | 0 |
|  | Sut1 | 24.61% | 17.27% | 0 |
|  | Mig2 | 17.10% | 21.50% | 0 |
|  | Sok2 | 50.26% | 8.60% | 2E-15 |
|  | Zap1 | 37.05% | 9.29% | 2.89E-13 |
|  | Sut1 | 14.48% | 17.09% | 1.897E-10 |
|  | Bas1 | 52.54% | 11.36% | 7.737E-09 |
|  | Mig2 | 8.47% | 17.92% | 2.243E-07 |
|  | Mot3 | 12.94% | 14.26% | 6.943E-06 |
|  | YPR015C | 4.01% | 18.84% | 9.679E-05 |

Genes activated by another TF, Mig2, are also enriched during both stresses. This is expected due to shift from galactose to glucose media in order to deplete L4 and Tef3 proteins (Lutfiyya and Johnston 1996). Another common TF acting as repressor between two sample types, is Sut1, which positively regulate genes for sterol uptake and negatively regulate gene expression required in filamentous growth in starvation condition (Foster, Cui et al. 2013). We observed elongated bat shaped cells in L4 and few in Tef3 depletion samples, indicating presence of filamentous growth under stress condition (Thapa *et al.*, 2013). Sut1 repressive activity is induced probably to inhibit the filamentous growth. Among the TFs unique to these two types of stresses are Upc2 in L4 depletion and Rlm1 in Tef3 depletion. Upc2 positively regulate ergosterol biosynthesis when sterol uptake from environment is reduced or when cellular sterol level is low (Vik A and Rine 2001). Transcription factor Rlm1 is activated by Pkc1 in cell wall integrity pathway and positively regulate expression of cell wall biogenesis genes (Jung, Sobering et al. 2002). Ergosterol biosynthesis genes are up regulated in L4 depletion sample (Supplementary fig 7) and cell wall biogenesis genes are upregulated in Tef3 depletion sample (Fig 9C).

## Discussion

Gene expression pattern changes in response to intra-cellular or environmental condition changes and helps cells to adapt better in the changed condition. However, gene expression pattern not necessarily always strongly correlate with changes in total cellular protein expression which ultimately determines cellular phenotypes. Since quantitative study of protein expression is challenging, researchers have always used quantitative gene expression studies to understand the molecular mechanism and signaling pathways behind cellular phenotypes. In budding yeast, we previously concluded using fluorescence microscopy and flow cytometry that abrogation of ribosome biogenesis by depleting an essential ribosomal protein causes G1 arrest and cell separation delay (M. Shamsuzzaman, A. Bommakanti, A. Zapinsky, N. Rahman, C. Pascual, and L. Lindahl, submitted). To understand the systemic gene expression response during ribosomal stress that leads to these phenotypes, we performed time series RNA-seq experiment on yeast cells depleted of ribosomal protein L4 over time.

The cell cycle arrest is a well-studied phenotype observed in cells during ribosomal stress (Bernstein and Baserga 2004, Fumagalli, Ivanenkov et al. 2012, Thapa, Bommakanti et al. 2013, Polymenis and Aramayo 2015) Shamsuzzman et al submitted. However, abrogation of ribosome biogenesis has a negative effect on the cell translational capacity. Consequently, it is not known whether the signaling pathway(s) that cause cell cycle arrest originate from the shut-down of the ribosome assembly pathways or the ensuing decline in translation capacity. To sort out these two types of signaling pathways we therefore compared the global gene expression following r-protein repression with the expression pattern after depletion of a translation elongation factor, Tef3, which inhibits translation, but does not affect ribosome biogenesis. We defined the stress triggering from depletion of Tef3 as translational stress.

### Difference in shift up response in Pgal-L4A and Pgal-TEF3 strains

The genes which differ most in expression pattern between the samples of two different types of stresses are ribosomal protein genes. Expression of both large and small ribosomal subunit protein genes went down in cells after 2hr depletion of ribosomal protein L4. In contrast, expression of both of these gene sets went up in 2hr after depletion of translation elongation factor Tef3. The up regulation of ribosomal protein genes in Pgal-L4 strain immediately after shift to glucose is accompanied with increased growth rate (Fig 1C) in this strain. Thus, the increase in ribosomal protein gene expression in Pgal-Tef3 may be induced by the shift to a more favorable carbon source, especially since the growth rate of Pgal-Tef3 also increased after the shift. It is well known that ribosomal protein gene expression correlate with growth rate (Kief and Warner 1981, Jorgensen, Nishikawa et al. 2002, Jorgensen and Tyers 2004, Castrillo, Zeef et al. 2007). Interestingly, we did not observe the shift up response in Pgal-L4A strain, which correlates with the absence of increased expression of r-protein genes. We measured growth rate of parent strain BY4741, Pgal-L43 and Pgal-S9 strains in galactose media and after shifting to glucose media. We did not observe any shift up response in these strains (Fig 1C). It is possible that under GAL promoter Tef3 protein was produced in excess, but that the growth rate was limited due to lack of favorable carbon source when grown in galactose media. After the shift to glucose, the excess amount of Tef3 protein was utilized under the favorable condition to support the growth increase.

### Trend in ribosomal gene expression differs between early and late hour of stress and also between ribosomal and translational stress

The trend in ribosomal protein gene expression changes in both strains – from down regulated at early to up regulated at late depletion samples in Pgal-L4A strain and reverse in Pgal-TEF3 strain. The downregulation of ribosomal protein genes observed in cells under translational stress is similar to the gene expression response observed in cells under several other types of environmental stresses (Gasch, Spellman et al. 2000, Belli, Molina et al. 2004). Under environmental stress conditions, ribosomal protein genes are downregulated as part of ~600 genes commonly repressed during stress response while the cellular transcriptional machinery is directed towards genes needed to adapt to the stress condition (Gasch, Spellman et al. 2000, Ho and Gasch 2015). This may be caused by TORC1 which under favorable environmental and nutritional conditions positively regulates the expression of ribosomal protein genes by RNA PolII (Martin, Soulard et al. 2004, Urban, Soulard et al. 2007, Fermi, Bosio et al. 2016). In stress conditions, in the absence of active TORC1 signaling pathway, expression of ribosomal protein genes goes down (Martin, Soulard et al. 2004, Schawalder, Kabani et al. 2004).

After Tef3 depletion, expression of ribosomal protein genes does not go up, even though translational capacity goes down, which mimics the common stress response. However, during ribosomal stress the expression of ribosomal protein genes increased, apparently because, cells sense the absence of an essential ribosomal protein. Ribosomal protein genes are regulated by Rap1 and Fhl1 that bind to specific upstream sites on RP genes in association with Ifh1, and positively regulate RP gene transcription in the presence of active TORC1 signaling pathway (Martin, Soulard et al. 2004, Fermi, Bosio et al. 2016). A small group of RP genes are regulated by Abf1, Fhl1 and Ifh1 transcriptional activators instead of Rab1. These genes respond similarly in response to favorable and stress condition. RPS22B gene, an exception in terms of regulation, has an Abf1 binding site, but expression is not affected by the decrease in Abf1 and Fhl1 occupancy (Fermi, Bosio et al. 2016). Possibly, because RPS22B genes has an additional transcription factor, Tbf1 binding site (Fermi, Bosio et al. 2016). RPS22B gene expression like other RP genes is sensitive to rapamycin treatment (Fermi, Bosio et al. 2016), attributing to TORC1 mediated regulation of this gene. Down regulation of RPS22B gene observed in our experiment, points to repressed TORC1 signaling pathways. In this case, existence of a TORC1 independent pathway only, explains up regulation of other RP genes in L4 depletion condition. Alternatively, Tbf1 binding to RPS22B promoter could be sensitive to ribosomal stress causing down regulation of this gene under stress. Interestingly, in both of the strains mitochondrial ribosomal protein genes are down (Fig 8E).

### Genes involved in translation are differentially regulated under ribosomal and translational stress

We looked at gene expression pattern of other cellular entities involved in translation in both if the stress sample types. Ribosome biogenesis (Ribi) genes are up regulated immediately after the shift to glucose both in Pgal-L4 and Pgal-TEF3 strains (Fig 8C). This up regulation can be explained by the activation of stress and nutrient sensitive transcriptional activator Sfp1 (Marion, Regev et al. 2004), which localizes to nucleus in both TORC1 dependent (Marion, Regev et al. 2004) and TORC1 independent manner (Singh and Tyers 2009). The Ribi genes are also regulated positively by Sch9 in a TORC1 dependent manner (Huber, Bodenmiller et al. 2009, Huber, French et al. 2011). Unlike Sch9, a Sfp1 deletion reduces expression of Ribi genes, but does not affect RP gene expression (Cipollina, van den Brink et al. 2008). The initial increase in expression seen at 2hr after shift to glucose could be the result of change in carbon source (up-shift) and Sfp1 localization to the nucleus. The subsequent decrease of expression of many Ribi genes after both L4 and Tef3 depletion may be due to TORC1 inactivation and subsequent Sfp1 re-localization to cytoplasm. We note however, that not all Ribi genes follow a common expression pattern after prolonged repression of RP and Tef3 genes, suggesting that the Ribi genes are not regulated by the exact same mechanism.

Unlike, RP genes in ribosomal stress, no compensatory up regulation of elongation factor genes were observed in Tef3 depleted samples (Fig 8D). Since, RP genes and Ribi genes are transcribed from RNA polII, we looked at expression pattern of RNA polII transcription cofactor activity. Interestingly, this gene set was up regulated in both L4 and Tef3 late depletion samples (Fig 8F). We also observed up regulation of protein transport related gene expression in Tef3 but not in L4 depletion samples. Under translational stress, cells appear to utilize the RNA polII transcription machinery to upregulate the expression of genes to maximize the delivery of the translated protein. On the other hand, under ribosomal stress, cells seemed to direct the RNA polII transcription machinery in maximizing expression of ribosomal protein genes.

### Cell cycle genes are regulated differentially under the two stress conditions examined

Cells after L4 depletion appeared to exit cell cycle and reside in G0 mode as we did not observe accumulation of any cell cycle phase specific transcripts in these cells (Fig 9A-E). Rather, we observed downregulation of most cell cycle genes specifically expressed in all phases (Fig 8A). To identify the upstream regulator of cell cycle phase transition to G0, we looked at over representation of target genes in L4 16h DEGs for transcription factors associated with G0 phase. From this analysis, we found over representation of target genes for Gis1 transcription factor in L4 16h depletion sample (Table 2). Gis1 is important in maintenance of stationary phase under starvation by inhibiting genes of cell division (Zhang, Wu et al. 2009). In contrast, cell cycle genes are differentially regulated in Tef3 depletion samples. Expression of genes specific to the M/G1 boundary were up in Tef3 depleted cells (Fig 9B and 9D). This also matches with up-regulation of cell integrity pathway genes (Fig 10C) observed in Tef3 depletion samples as this pathway also activate SBF transcription factor in M/G1 boundary which turn on expression of genes required for G1 to S progression.

### Translational stress activate cell wall integrity pathway

Cell growth requires continuous protein synthesis as well as cell membrane and cell wall expansion. Interrupting the function or deletion of any component in the secretory pathway, starting from peptide insertion into endoplasmic reticulum to vesicle fusion with cytoplasmic membrane activates cell integrity pathway and represses transcription of genes involved in ribosome synthesis including ribosomal protein gene and rRNA gene expression (Mizuta and Warner 1994, Nierras and Warner 1999). This signaling pathway depends on Pkc1, a kinase required for remodeling of cell wall during growth. However, the downstream Bck1-Mkk1/2–Slt2 mitogen-activated protein kinase cascade and continuous protein synthesis is not required for the link of ribosome gene expression to secretion (Nierras and Warner 1999, Li, Moir et al. 2000). Rather, an unknown branch of Pkc1 signaling interacts with the Rap1 transcription factor C-terminal domain and represses RP and Ribi gene expression (Mizuta, Tsujii et al. 1998). In our experiment, Tef3 depletion samples showed elevated expression of genes of secretory pathway including ER to Golgi transport, Golgi vesicular transport, exocytosis and fungal cell wall biogenesis genes (Fig 11A-E). These genes are induced under cell wall stress in cell integrity pathway (Bulik, Olczak et al. 2003). TF over representation analysis also showed enrichment of Rlm1 transcription factor target genes in Tef3 26hr sample(Table 2), which works as an effector of the Pkc1 signaling pathway and turns on expression of cell wall biogenesis genes (Jung, Sobering et al. 2002). Moreover, RP and Ribi genes were repressed during late hour of Tef3 depletion (Fig 8A and 8B). All these observations indicate activation of cell integrity pathway and Pkc1 mediated signaling in cells under translational stress. Cells with defects in secretory pathway and continuous protein synthesis are thought to build intracellular turgor pressure which in turn causes membrane stress and activate cell integrity pathway. In case of Tef3 depletion, the story is probably other way around. Cells stop synthesizing protein and as a result turgor pressure probably drops significantly. This drop in pressure is likely detected in the membrane and results in activation of cell integrity pathway. In support of this hypothesis, we did not detect activation of cell integrity pathway in cells depleted for L4 protein. Interruption of ribosome biogenesis does not stop protein synthesis function per se which in turn probably does not affect cellular turgor pressure.

### Depletion of ribosomal protein causes oxidative stress and acidification of cell

Genes involved in detoxifying oxidative stress is observed to be induced in cells with L4 depletion. Some of these genes are induced as early as in 2h. We observed the enrichment of the reactive oxygen species GO term in both L4 2h and 16h depletion samples (Supp. fig 1 and Supp. Fig 2). One of the cause of oxidative stress is accumulation of hydroperoxides. Cells metabolize these peroxides with inducing expression of peroxidases such as catalases and glutathione peroxidases. Under oxidative stress, cells can get exposed to small molecule hydroperoxides like H_2_O_2_. These small molecules hydroperoxides are neutralized by catalases CTT1 (cytoplasmic), CTA1 (mitochondrial) and CPA1 (peroxisomal). CTT1 is highly induced in cells with L4 depletion starting from 2h (Supp. Fig 8). In addition, cells can get exposed to large molecules hydroperoxide like *tert-butyl* hydroperoxide (*t*-BHP) and lipid hydroperoxide. Glutathione peroxidases (GRX and GPX proteins) neutralizes these peroxides (Avery and Avery 2001). For neutralizing membrane lipid hydroperoxides, a specific type of glutathione peroxidase - phospholipid hydroperoxide glutathione peroxidase is responsible (Avery and Avery 2001). In cells under ribosomal stress, these glutathione peroxidase genes are upregulated, most notably GPX2 a phospholipid hydroperoxide glutathione peroxidase is induced 7 fold as early as in 2h (Supp. Fig 8). This indicates that the cell membrane components in ribosomal stress are probably oxidized. Increased oxidized glutathione are exported from cell through GEX1 (Dhaoui, Auchère et al. 2011). GEX1 exports glutathione and imports H+ and causes cellular acidification. Glutathione exporter (GEX) function is important for cells to survive in oxidative stress (Dhaoui, Auchère et al. 2011). We observed upregulation of GEX1 in cells under ribosomal stress. GPX2 probably plays predominant role in protecting cells from oxidative stress in the presence of Ca2+ (Tanaka, Izawa et al. 2005).

Cells under ribosomal stress are probably defective in vacuolar acidification. We observed increased expression of Vma3 and Vma13, involved in vacuolar acidifications which are induced under oxidative stress (Supp. Fig 7) (Belli, Molina et al. 2004). We observed upregulation of ergosterol biosynthesis genes in L4 depletion samples and the enrichment of Upc2 transcription factor responsible for ergosterol biosynthesis gene upregulation (Supplementary Fig 7, Table 2). These genes are induced in cells as a result of decrease in sterol uptake (Smith, Crowley et al. 1996). Vacuolar ATPase function, important in vacuolar acidification, requires ergosterol (Zhang, Gamarra et al. 2010). Cells under ribosomal stress probably have ergosterol deficiency which causes abnormality in vacuolar acidification and thus increase cellular pH level and reactive oxygen species (ROS) level (Niles and Powers 2014). Overexpression of Gex1 probably increases pH of cells under ribosomal stress which in turn activates PKA pathway and represses Pkc1 mediated cell integrity pathway (Dechant, Binda et al. 2010, Dhaoui, Auchère et al. 2011).

### Increase in Reactive oxygen species (ROS) probably cause actin depolarization in cells under ribosomal stress

Cells under oxidative stress demonstrate actin depolarization (Vilella, Herrero et al. 2005). Increase amount of ROS in oxidative stress oxidizes specific cysteine residues in actin protein and is thought of as a possible mechanism for actin depolarization (Niles and Powers 2014). Increase in cellular ROS is observed mainly in three ways – first, through cellular respiration and protein kinase A mediated mitochondrial dysfunction; secondly from defects in cellular acidification and finally from hypoactive Pkc1 signaling pathway (Vilella, Herrero et al. 2005, Niles and Powers 2014). We observed actin depolarization in L4 depleted cells as early as at 2hr of depletion. In RNA-seq experiment, we have observed expression of multiple gene sets indicating elevated level of ROS in the cells depleted for L4 depletion and in Tef3 depleted cells these phenotypes appearing later. We speculate that increase reactive oxygen species under oxidative stress probably cause actin depolarization observed in the cells under ribosomal stress. In cells with translational stress, there is also signature of gene expression responsive to oxidative stress. But, that is much weaker than it is in L4 depletion samples. It matches with our delayed observation of actin depolarization in Tef3 depletion samples. In these cells, possible activation of cell integrity pathway may help in downregulating cellular ROS.

### Elevated expression of genes involved in degradation of membrane proteins is detected in both of these stress conditions

We observed a positive trend in up regulation of *COS* genes in cells depleted for either L4 or Tef3 protein (Fig 12C). There are 12 Cos genes in yeast. This gene set is induced in nutrient stress condition and causes down regulation of cell surface proteins, predominantly GIP anchored membrane proteins (MacDonald, Payne et al. 2015). Cos proteins work as trans ubiquitination signal for sorting of membrane proteins into multi vesicular bodies for degradation (MacDonald, Payne et al. 2015). Expression of Cos proteins leads to formation of large endosomal vesicles observed by fluorescence tagging of membrane proteins (MacDonald, Payne et al. 2015). In cells with both ribosomal stress and translational stress, we observed large endosomal vesicles by tagging Ras2 membrane protein with GFP and consequent fluorescence microscopy (Fig 12D, Supplementary fig 6). We speculate that under these two stresses, membrane proteins are downregulated from the cell surface which includes TORC2 and its mediators Rom2, Slm1 and Slm2 (Niles and Powers 2012). That in turn, may lead to down regulation of TORC2-Ypk1/Ypk2 signaling pathway causing actin depolarization and elevation of ROS.

## Concluding remarks

Ribosomal stress is sensed early after ribosomal protein depletion in the cell, as early as 2hr. Up regulation of genes responsive to oxidative stress and over representation of mRNAs for TFs responsive to stress is detected in cell at 2hr of ribosomal protein depletion. Even though, we detected phenotypic similarities in terms of cell separation and G1 cell cycle accumulation in cells with ribosomal and translational stress, they differ in gene expression pattern underlying these phenotypes indicating difference in causalities of these phenotypes. Both ribosomal and translational stress show common stress responsive gene expression like Crz1 target gene expression, signature of oxidative stress response and finally membrane instability. Cell membrane and cell wall acts as a major stress sensor in the cell and adjust cellular metabolism accordingly. Any change in membrane lipid composition, or membrane protein oxidation, or decrease or increase in intracellular turgor pressure causes stress in cell membrane. Cell membrane stress activates and/or inactivates specific signaling pathway which triggers stress responsive gene expression and adaptation of cellular behavior accordingly.

## Supporting Information

**Supplementary fig 1:** Over representation analysis of all differentially expressed genes in L4 2hr sample.

**Supplementary fig 2:** Over representation analysis of all differentially expressed genes in L4 16hr sample.

**Supplementary fig 3:** Treemap showing GO terms enriched in Tef3 2hr gene expression data with respect to 0hr data.

**Supplementary fig 4:** Treemap showing GO terms enriched in Tef3 26hr gene expression data with respect to 0hr data.

**Supplementary fig 5:** Heatmap showing log fold change in expression of genes which are expressed by the daughter cell after cytokinesis.

**Supplementary fig 6:** Fluorescence microscopy images showing engulfing and sorting of GFP tagged Ras2 membrane protein L43 depletion sample.

**Supplementary fig 7:** Heatmap showing log fold change in expression of genes associate with Ergosterol biosynthesis

**Supplementary fig 8:** Heatmap and scatter plot showing log fold change in expression of genes involved in ROS metabolism and acidification of cell

**Supplementary file 1:** List of Go terms from enrichment analysis of L4 and Tef3 genes ranked by sum of absolute rank differences

